# Consensus clustering for Bayesian mixture models

**DOI:** 10.1101/2020.12.17.423244

**Authors:** Stephen Coleman, Paul D.W. Kirk, Chris Wallace

**Author notes:** These authors provided an equal contribution (Paul D.W. Kirk, Chris Wallace).

## Abstract

Cluster analysis is an integral part of precision medicine and systems biology, used to define groups of patients or biomolecules. Consensus clustering is an ensemble approach that is widely used in these areas, which combines the output from multiple runs of a non-deterministic clustering algorithm. Here we consider the application of consensus clustering to a broad class of heuristic clustering algorithms that can be derived from Bayesian mixture models (and extensions thereof) by adopting an early stopping criterion when performing sampling-based inference for these models. While the resulting approach is non-Bayesian, it inherits the usual benefits of consensus clustering, particularly in terms of computational scalability and providing assessments of clustering stability/robustness.

In simulation studies, we show that our approach can successfully uncover the target clustering structure, while also exploring different plausible clusterings of the data. We show that, when a parallel computation environment is available, our approach offers significant reductions in runtime compared to performing sampling-based Bayesian inference for the underlying model, while retaining many of the practical benefits of the Bayesian approach, such as exploring different numbers of clusters. We propose a heuristic to decide upon ensemble size and the early stopping criterion, and then apply consensus clustering to a clustering algorithm derived from a Bayesian integrative clustering method. We use the resulting approach to perform an integrative analysis of three ‘omics datasets for budding yeast and find clusters of co-expressed genes with shared regulatory proteins. We validate these clusters using data external to the analysis. These clusters can help assign likely function to understudied genes, for example *GAS3* clusters with histones active in S-phase, suggesting a role in DNA replication.

Our approach can be used as a wrapper for essentially any existing sampling-based Bayesian clustering implementation, and enables meaningful clustering analyses to be performed using such implementations, even when computational Bayesian inference is not feasible, e.g. due to poor exploration of the target density (often as a result of increasing numbers of features) or a limited computational budget that does not along sufficient samples to drawn from a single chain. This enables researchers to straightforwardly extend the applicability of existing software to much larger datasets, including implementations of sophisticated models such as those that jointly model multiple datasets.

## Background

From defining a taxonomy of disease to creating molecular sets, grouping items can help us to understand and make decisions using complex biological data. For example, grouping patients based upon disease characteristics and personal omics data may allow the identification of more homogeneous subgroups, enabling stratified medicine approaches. Defining and studying molecular sets can improve our understanding of biological systems as these sets are more interpretable than their constituent members (1), and study of their interactions and perturbations may have ramifications for diagnosis and drug targets (2, 3). The act of identifying such groups is referred to as *cluster analysis*. Many traditional methods such as *K*-means clustering (4, 5) condition upon a fixed choice of *K*, the number of clusters. These methods are often heuristic in nature, relying on rules of thumb to decide upon a final value for *K*. For example, different choices of *K* are compared under some metric such as silhouette (6) or the within-cluster sum of squared errors (**SSE**) as a function of *K*. Moreover, *K*-means clustering can exhibit sensitivity to initialisation, necessitating multiple runs in practice (7).

Another common problem is that traditional methods offer no measure of the stability or robustness of the final clustering. Returning to the stratified medicine example of clustering patients, there might be individuals that do not clearly belong to any one particular cluster; however if only a point estimate is obtained, this information is not available. Ensemble methods address this problem, as well as reducing sensitivity to initialisation. These approaches have had great success in supervised learning, most famously in the form of Random Forest (8) and boosting (9). In clustering, consensus clustering (10) is a popular method which has been implemented in `R` (11) and to a variety of methods (12, 13) and been applied to problems such as cancer subtyping (14, 15) and identifying subclones in single cell analysis (16). Consensus clustering uses *W* runs of some base clustering algorithm (such as *K*-means). These *W* proposed partitions are commonly compiled into a *consensus matrix*, the (*i, j*)^*th*^ entries of which contain the proportion of model runs for which the *i*^*th*^ and *j*^*th*^ individuals co-cluster (for this and other definitions see section 1 of the Supplementary Material), although this step is not fundamental to consensus clustering and there is a large body of literature aimed at interpreting a collection of partitions (see, e.g., 17, 18, 19). This consensus matrix provides an assessment of the stability of the clustering. Furthermore, ensembles can offer reductions in computational runtime because the individual members of the ensemble are often computationally inexpensive to fit (e.g, because they are fitted using only a subset of the available data) and because the learners in most ensemble methods are independent of each other and thus enable use of a parallel environment for each of the quicker model runs (20).

Traditional clustering methods usually condition upon a fixed choice of *K*, the number of clusters with the choice of *K* being a difficult problem in itself. In consensus clustering, Monti *et al*. (10) proposed methods for choosing *K* using the consensus matrix and Ünlü *et al*. (21) offer an approach to estimating *K* given the collection of partitions, but each clustering run uses the same, fixed, number of clusters. An alternative clustering approach, mixture modelling, embeds the cluster analysis within a formal, statistical framework (22). This means that models can be compared formally, and problems such as the choice of *K* can be addressed as a model selection problem (23). Moreover, *Bayesian mixture models* can be used to try to directly infer *K* from the data. Such inference can be performed through use of a Dirichlet Process mixture model (24, 25, 26), a mixture of finite mixture models (27, 28) or an over-fitted mixture model (29). The Bayesian model also assesses the uncertainty in the cluster allocations, and if *K* is treated as a random variable uncertainty about the value of *K* propagates through to the clustering. Furthermore, the Bayesian hierarchical modelling framework enables extrapolating the mixture model to capture more complex dependencies, for example, integrative clustering methods tailored for multi-omics analysis (30, 31, 32). Bayesian clustering methods have a history of successful application to a diverse range of biological problems such as finding clusters of gene expression profiles (33), cell types in flow cytometry (34, 35) or scRNAseq experiments (36), and estimating protein localisation (37).

Markov chain Monte Carlo (MCMC) methods are the most common tool for performing computational Bayesian inference. They guarantee an exact description of the posterior distribution in the limit of infinite iterations in contrast to Variational Inference (38). In Bayesian clustering, they are used to draw a collection of clustering partitions from the posterior distribution. In practice MCMC methods may explore the parameter space very inefficiently despite their ergodic properties. As the number of features/measurements increases this inefficiency can become pathological with chains prone to becoming stuck in local posterior modes preventing convergence in any feasible period of runtime (see, e.g., the Supplementary Materials of 39); this problem is frequently referred to as poor mixing within the chain. There is a rich zoo of MCMC methods designed to overcome the different limitations of the most basic samplers. For example, there are MCMC methods that use parallel chains to improve the scalability with an increasing number of observations, such as Consensus Monte Carlo (40, 41, 42). Consensus Monte Carlo methods subsample the original dataset and run separate chains on each smaller dataset. In this way they can use a far smaller quantity of data for each Monte Carlo algorithm and treat each chain as embarrassingly parallel enabling simultaneous model runs across machines, with the samples then averaged across chains. This parallelisation and reduced dataset size offers a significant reduction in runtime for large *N* datasets. Another method designed to improve scaling to large datasets, is stochastic gradient MCMC (SGMCMC 43). This uses a subset of the data in each sampling iteration and has provable guarantees (44). However, SGMCMC converges at a slower rate than traditional MCMC algorithms and remains computationally costly (45, 46). While these methods help in scaling to large N data, they are less helpful in situations where we have high-dimension but only moderate sample size, such as frequently arises in analysis of ‘omics data. Other methods such as coupling (47) use multiple chains to reduce the bias of the Monte Carlo estimate.

Other MCMC methods make efforts to overcome the problem of poor mixing at the cost of increased computational cost per iteration (48). In clustering models introducing split-merge moves into the sampler are the most common examples of such bold exploration moves (see, e.g., 49, 50, 51, 52). However, these methods are difficult to implement and frequently propose many rejected moves, thereby increasing computational cost without necessarily guaranteeing full exploration of the target density in any finite amount of time. Furthermore, most available Bayesian clustering methods are implemented using a basic Gibbs sampler and would require reimplementation to exploit more scalable samplers, a costly investment of time and effort. Ideally these existing implementations could be used despite their simple sampler.

There also exists a range of alternative clustering methods that are designed or have been extended to scale well with increasing sample size, e.g., *k*-means clustering (53, 54), spectral clustering (55, 56), density-based clustering (57, 58), etc., any of which could be used within a consensus clustering wrapper. However, while these methods have better scalability than sampling based clustering methods, they suffer from a lack of flexibility. They do not, in general, have the ability to explore multiple values of *K* in a single model run, it is not easy to extend these methods to the multiple dataset problem and they are often restricted to a specific data type.

As described above, sampling-based Bayesian clustering methods are flexible, capable of modelling complex dependencies and the number of clusters present. However, they suffer from prohibitive runtimes and poor exploration in high-dimensional data (i.e., large *P* data). This limits the consistency of their inference in biomedical applications where data is often very high dimensional with only moderate sample size (for some discussion around stability in clustering, please see 59, 60, 61). Motivated by this, we aim to develop a general and straightforward procedure that exploits the flexibility of Bayesian model based clustering methods but improves their performance under a constrained computational budget without requiring reimplementation. Specifically, we make use of existing sampling-based Bayesian clustering implementations, but only run them for a fixed (and relatively small) number of iterations, stopping before they have converged to their target stationary distribution. We initialise each chain with a random draw from an uninformative prior distribution on the space of partitions and then collect the final sampled partition. Doing this repeatedly, we obtain an ensemble of clustering partitions which has large variety in its initialisation. We use this set to perform consensus clustering, constructing a consensus matrix (thereby avoiding the label-switching problem) which describes uncertainty about the latent structure in the data. This can be used to infer a point estimate clustering. We propose a heuristic for deciding upon the ensemble size (the number of learners used, *W*) and the ensemble depth (the number of iterations, *D*), inspired by the use of scree plots in Principal Component Analysis (**PCA**; 62). We hope to show a way of scaling Bayesian mixture models and their extensions with increasing numbers of features that can explore a range of *K* in a single model run and can be tailored to specific properties of a given dataset. We note that, despite the similarity in names, our consensus clustering approach for Bayesian mixture models is very different to the consensus Monte Carlo approach of Ni *et al*. (41), which was designed to enable Bayesian mixture models to scale to large *N* datasets. Our approach leans into the ensemble framework of Monti *et al*. (10); we consider the case that our individual chains are too short to have converged and in which case the inference is non-Bayesian, in contrast to Consensus Monte Carlo. Our primary aim is to mitigate the problem of poor mixing which tends to emerge when the data has a high numbers of features relative to the sample size and individual chains struggle to converge in a reasonable runtime, and to enable the use of complex models such as arise in multi-view or integrative analyses for which each iteration of the MCMC algorithm is slow even for small sample data and running a long chain might not be feasible.

We show via simulation that our approach can successfully identify meaningful clustering structures even when chains are very short. We then illustrate the use of our approach to extend the applicability of existing Bayesian clustering implementations, using as a case study the Multiple Dataset Integration (**MDI** 30, section 2 of the Supplementary Material) model for Bayesian integrative clustering applied to real data. While the simulation results serve to validate our method, it is important to also evaluate methods on real data which may represent more challenging problems. For our real data, we use three ‘omics datasets relating to the cell cycle of *Saccharomyces cerevisiae* with the aim of inferring clusters of genes across datasets. As there is no ground truth available, we then validate these clusters using knowledge external to the analysis.

## Material and methods

### Consensus clustering for Bayesian mixture models

We apply consensus clustering to MCMC based Bayesian clustering models using the method described in algorithm 1. Our application of consensus clustering has two main parameters at the ensemble level, the chain depth, *D*, and ensemble width, *W*. We infer a point clustering from the consensus matrix using the `maxpear` function (63) from the R package `mcclust` (64) which maximises the posterior expected adjusted Rand index between the true clustering and point estimate if the matrix is composed of samples drawn from the posterior distribution (section 3 of the Supplementary Material for details). There are alternative choices of methods to infer a point estimate which minimise different loss functions (see, e.g., 65, 66, 67).

#### Algorithm 1

Consensus clustering for Bayesian mixture models.

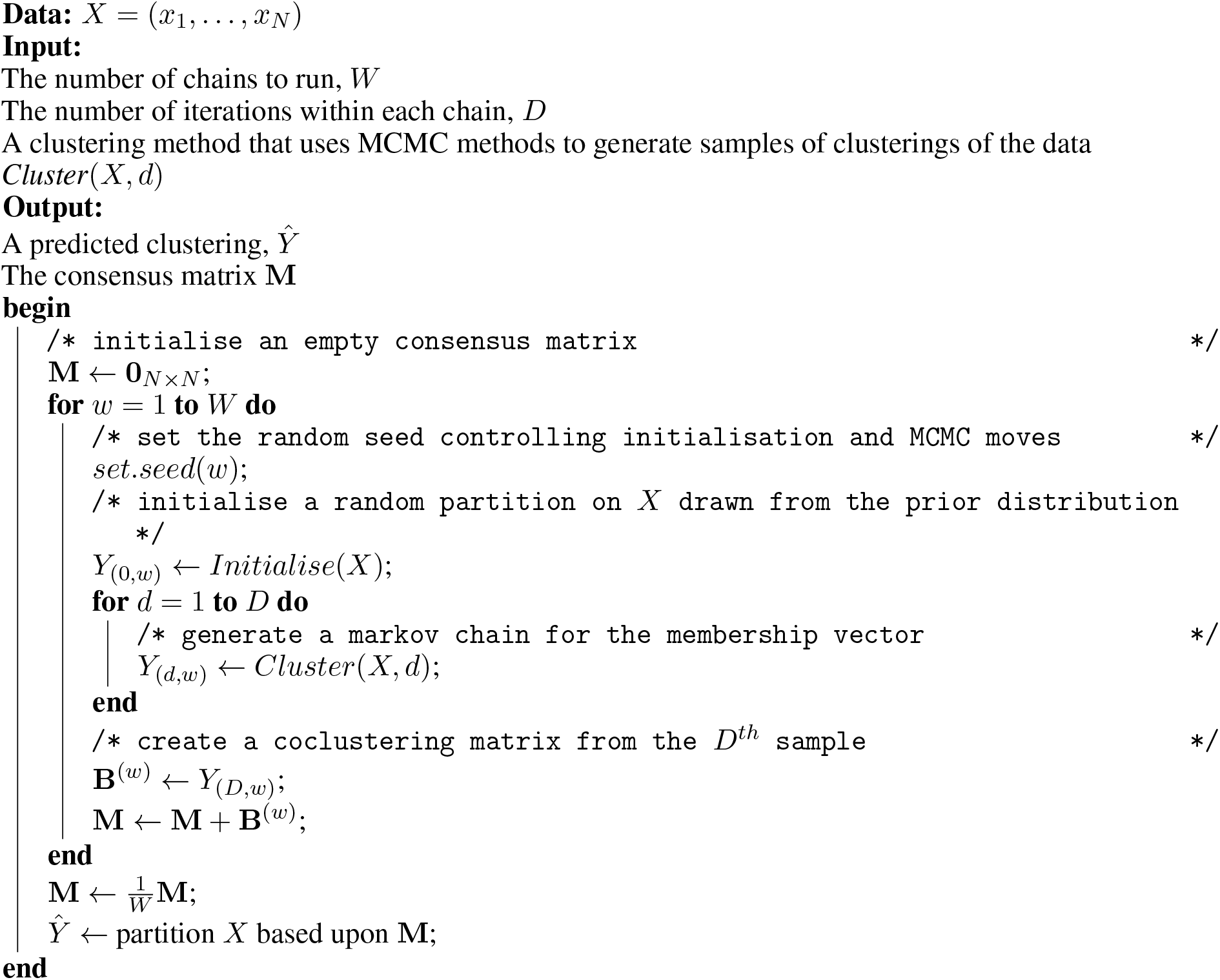

#### Determining the ensemble depth and width

As our ensemble sidesteps the problem of convergence within each chain, we need an alternative stopping rule for growing the ensemble in chain depth, *D*, and number of chains, *W*. We propose a heuristic based upon the consensus matrix to decide if a given value of *D* and *W* are sufficient. We suspect that increasing *W* and *D* might continuously improve the performance of the ensemble, but we observe in our simulations that these changes will become smaller and smaller for greater values, eventually converging for each of *W* and *D*. We notice that this behaviour is analogous to PCA in that where for consensus clustering some improvement might always be expected for increasing chain depth or ensemble width, more variance will be captured by increasing the number of components used in PCA. However, increasing this number beyond some threshold has diminishing returns, diagnosed in PCA by a scree plot. Following from this, we recommend, for some set of ensemble parameters, *D*^′^ = {*d*_1_, …, *d*_*I*_} and *W*^′^ = {*w*_1_, …, *w*_*J*_ }, find the mean absolute difference of the consensus matrix for the 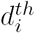 iteration from *w*_*j*_ chains to that for the 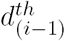 iteration from *w*_*j*_ chains and plot these values as a function of chain depth, and the analogue for sequential consensus matrices for increasing ensemble width and constant depth.

If this heuristic is used, we believe that the consensus matrix and the resulting inference should be stable (see, e.g., 59, 60), providing a robust estimate of the clustering. In contrast, if there is still strong variation in the consensus matrix for varying chain length or number, then we believe that the inferred clustering is influenced significantly by the random initialisation and that the inferred partition is unlikely to be stable for similar datasets or reproducible for a random choice of seeds.

### Simulation study

We use a finite mixture with independent features as the data generating model within the simulation study. Within this model there exist “irrelevant features” (68) that have global parameters rather than cluster specific parameters. The generating model is

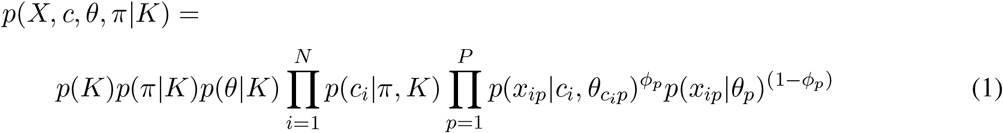

for data *X* = (*x*_1_, …, *x*_*N*_), cluster label or allocation variable *c* = (*c*_1_, …, *c*_*N*_), cluster weight *π* = (*π*_1_, …, *π*_*K*_), *K* clusters and the relevance variable, *ϕ* ∈ {0, 1} with *ϕ*_*p*_ = 1 indicating that the *p*^*th*^ feature is relevant to the clustering. We used a *Gaussian* density, so 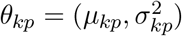. We defined three scenarios and simulated 100 datasets in each (figure 1 and table 1). Additional details of the simulation process and additional scenarios are included in section 4.1 of the Supplementary Materials.

**Figure 1:**
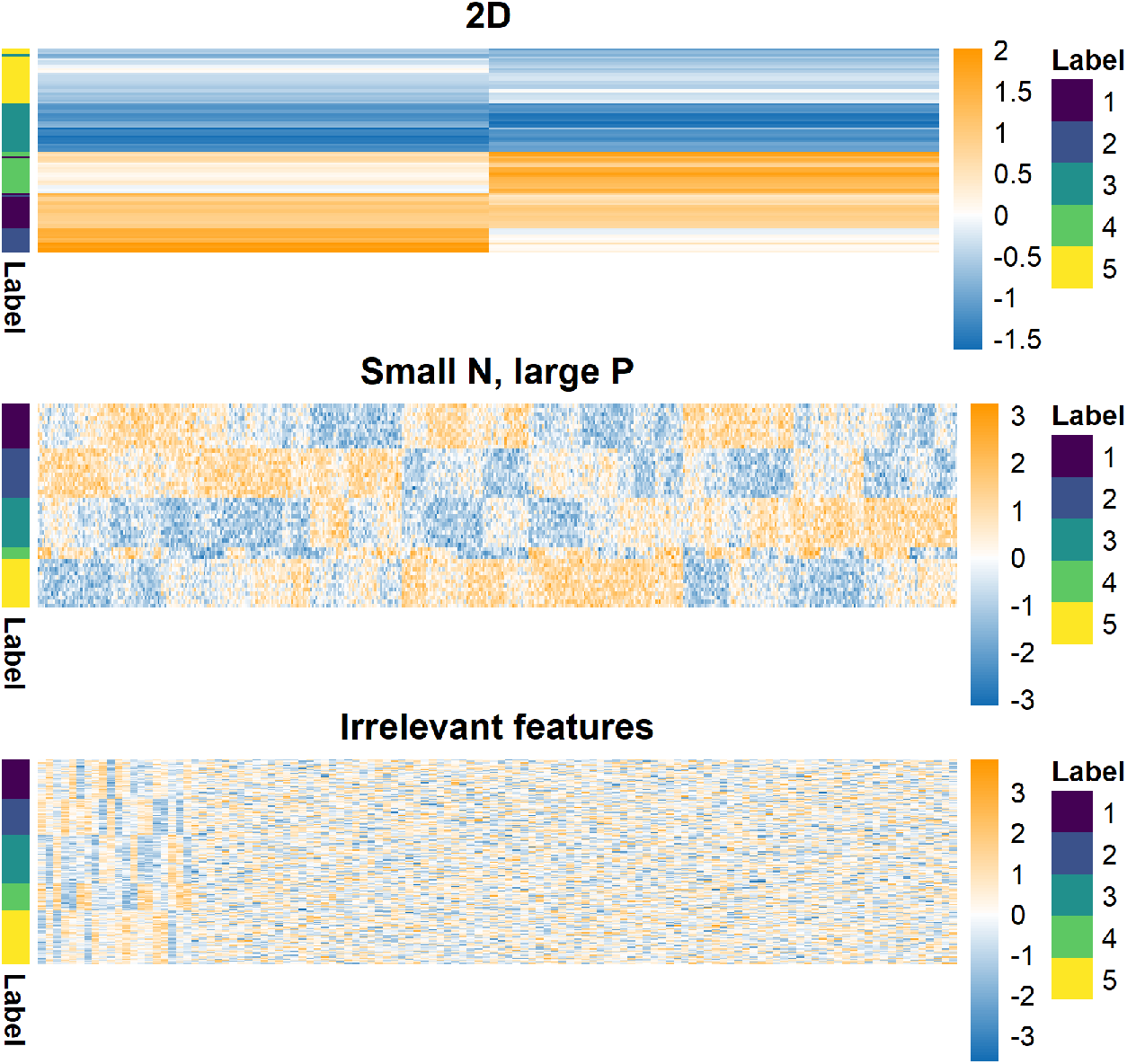
Example of generated datasets. Each row is an item being clustered and each column a feature of generated data. The 2D dataset (which is ordered by hierarchical clustering here) should enable proper mixing of chains in the MCMC. The small *N*, large *P* case has clear structure (observable by eye). This is intended to highlight the problems of poor mixing due to high dimensions even when the generating labels are quite identifiable. In the irrelevant features case, the structure is clear in the relevant features (on the left-hand side of this heatmap). This setting is intended to test how sensitive each approach is to noise.

**Table 1:** Parameters defining the simulation scenarios as used in generating data and labels. Δ*µ* is the distance between neighbouring cluster means within a single feature. The number of relevant features (*P*_*s*_) is∑_*p*_ *ϕ*_*p*_, and *P*_*n*_ = *P* − *P*_*s*_.

| Scenario | $N$ | $P_s$ | $P_n$ | $K$ | $\Delta\mu$ | $\sigma^2$ | $\pi$ |
| --- | --- | --- | --- | --- | --- | --- | --- |
| 2D | 100 | 2 | 0 | 5 | 3.0 | 1 | $(\frac{1}{5}, \frac{1}{5}, \frac{1}{5}, \frac{1}{5}, \frac{1}{5})$ |
| Small N, large P | 50 | 500 | 0 | 5 | 1.0 | 1 | $(\frac{1}{5}, \frac{1}{5}, \frac{1}{5}, \frac{1}{5}, \frac{1}{5})$ |
| Irrelevant features | 200 | 20 | 100 | 5 | 1.0 | 1 | $(\frac{1}{5}, \frac{1}{5}, \frac{1}{5}, \frac{1}{5}, \frac{1}{5})$ |

In each of these scenarios we apply a variety of methods (listed below) and compare the inferred point clusterings to the generating labels using the Adjusted Rand Index (**ARI**, 69).

- `Mclust`, a maximum likelihood implementation of a finite mixture of Gaussian densities (for a range of modelled clusters, *K*),
- 10 chains of 1 million iterations, thinning to every thousandth sample for the overfitted Bayesian mixture of Gaussian densities, and
- A variety of consensus clustering ensembles defined by inputs of *W* chains and *D* iterations within each chain (see algorithm 1) with *W* ∈ {1, 10, 30, 50, 100} and *D* ∈ {1, 10, 100, 1000, 10000} where the base learner is an overfitted Bayesian mixture of Gaussian densities.

Note that none of the applied methods include a model selection step and as such there is no modelling of the relevant variables. This and the unknown value of *K* is what separates the models used and the generating model described in equation 1. More specifically, the likelihood of a point *X*_*n*_ for each method is

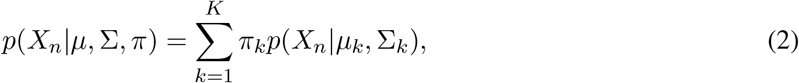

where *p*(*X*_*n*_|*µ*_*k*_, Σ_*k*_) is the probability density function of the multivariate Gaussian distribution parameterised by a mean vector, *µ*_*k*_, and a covariance matrix, `Σ`_*k*_, and *π*_*k*_ is the component weight such that 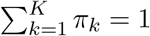. The implementation of the Bayesian mixture model restricts Σ_*k*_ to be a diagonal matrix while `Mclust` models a number of different covariance structures. Note that while we use the overfitted Bayesian mixture model, this is purely from convenience and we expect that a true Dirichlet Process mixture or a mixture of mixture models would display similar behaviour in an ensemble.

The ARI is a measure of similarity between two partitions,*c*_1_, *c*_2_, corrected for chance, with 0 indicating *c*_1_ is no more similar to *c*_2_ than a random partition would be expected to be and a value of 1 showing that *c*_1_ and *c*_2_ perfectly align. Details of the methods in the simulation study can be found in sections 4.2, 4.3 and 4.4 of the Supplementary Material.

`Mclust`

`Mclust` (70) is a function from the R package `mclust`. It estimates Gaussian mixture models for *K* clusters based upon the maximum likelihood estimator of the parameters. It initialises upon a hierarchical clustering of the data cut to *K* clusters. A range of choices of *K* and different covariance structures are compared and the “best” selected using the Bayesian information criterion, (71) (details in section 4.2 of the Supplementary Material).

#### Bayesian inference

To assess within-chain convergence of our Bayesian inference we use the Geweke *Z*-score statistic (72). Of the chains that appear to behave properly we then asses across-chain convergence using 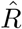 (73) and the recent extension provided by (74). If a chain has reached its stationary distribution the Geweke *Z*-score statistic is expected to be normally distributed. Normality is tested for using a Shapiro-Wilks test (75). If a chain fails this test (i.e., the associated *p*-value is less than 0.05), we assume that it has not achieved stationarity and it is excluded from the remainder of the analysis. The samples from the remaining chains are then pooled and a posterior similarity matrix (**PSM**) constructed. We use the `maxpear` function to infer a point clustering. For more details see section 4.3 of the Supplementary Material.

### Analysis of the cell cycle in budding yeast

#### Datasets

The cell cycle is crucial to biological growth, repair, reproduction, and development (76, 77, 78) and is highly conserved among eukaryotes (78).. This means that understanding of the cell cycle of *S. cerevisiae* can provide insight into a variety of cell cycle perturbations including those that occur in human cancer (79, 77) and ageing (80). We aim to create clusters of genes that are co-expressed, have common regulatory proteins and share a biological function. To achieve this, we use three datasets that were generated using different ‘omics technologies and target different aspects of the molecular biology underpinning the cell cycle process.

- Microarray profiles of RNA expression from (81), comprising measurements of cell-cycle-regulated gene expression at 5-minute intervals for 200 minutes (up to three cell division cycles) and is referred to as the **time course** dataset. The cells are synchronised at the START checkpoint in late G1-phase using alpha factor arrest (81). We include only the genes identified by (81) as having periodic expression profiles.
- Chromatin immunoprecipitation followed by microarray hybridization (**ChIP-chip**) data from (82). This dataset discretizes *p*-values from tests of association between 117 DNA-binding transcriptional regulators and a set of yeast genes. Based upon a significance threshold these *p*-values are represented as either a 0 (no interaction) or a 1 (an interaction).
- Protein-protein interaction (**PPI**) data from BioGrid (83). This database consists of of physical and genetic interactions between gene and gene products, with interactions either observed in high throughput experiments or computationally inferred. The dataset we used contained 603 proteins as columns. An entry of 1 in the (*i, j*)^*th*^ cell indicates that the *i*^*th*^ gene has a protein product that is believed to interact with the *j*^*th*^ protein.

The datasets were reduced to the 551 genes with no missing data in the PPI and ChIP-chip data, as in (30).

#### Multiple dataset integration

We applied consensus clustering to MDI for our integrative analysis. Details of MDI are in section 2.2 of the Supplementary Material, but in short MDI jointly models the clustering in each dataset, inferring individual clusterings for each dataset. These partitions are informed by similar structure in the other datasets, with MDI learning this similarity as it models the partitions. The model does not assume global structure. This means that the similarity between datasets is not strongly assumed in our model; individual clusters or genes that align across datasets are based solely upon the evidence present in the data and not due to strong modelling assumptions. Thus, datasets that share less common information can be included without fearing that this will warp the final clusterings in some way.

The datasets were modelled using a mixture of Gaussian processes in the time course dataset and Multinomial distributions in the ChIP-chip and PPI datasets.

## Results

### Simulated data

We use the ARI between the generating labels and the inferred clustering of each method to be our metric of predictive performance. In figure 2, we see `Mclust` performs very well in the 2D and Small *N*, large *P* scenarios, correctly identifying the true structure. However, the irrelevant features scenario sees a collapse in performance, `Mclust` is blinded by the irrelevant features and identifies a clustering of *K* = 1.

**Figure 2:**
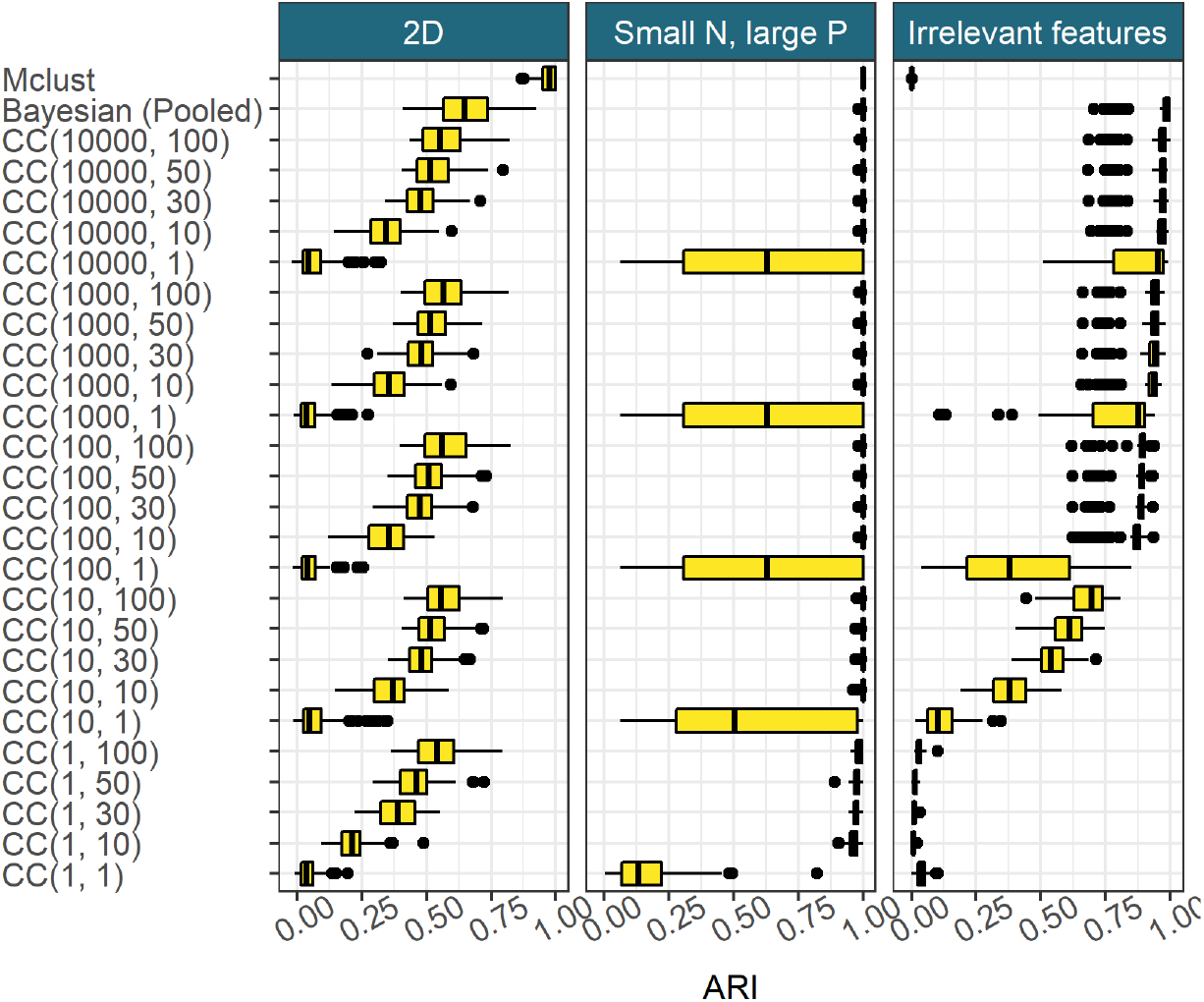
Model performance in the 100 simulated datasets for each scenario, defined as the ARI between the generating labels and the inferred clustering. *CC*(*d, w*) denotes consensus clustering using the clustering from the *d*^*th*^ iteration from *w* different chains.

The pooled samples from multiple long chains performs very well across all scenarios and appears to act as an upper bound on the more practical implementations of consensus clustering.

Consensus clustering does uncover some of the generating structure in the data, even using a small number of short chains. With sufficiently large ensembles and chain depth, consensus clustering is close to the pooled Bayesian samples in predictive performance. It appears that for a constant chain depth increasing the ensemble width used follows a pattern of diminishing returns. There are strong initial gains for a greater ensemble width, but the improvement decreases for each successive chain. A similar pattern emerges in increasing chain length for a constant number of chains (figure 2).

For the PSMs from the individual chains, all entries are 0 or 1 (figure 3). This means only a single clustering is sampled within each chain, implying very little uncertainty in the partition. However, three different clustering solutions emerge across the chains, indicating that each individual chain is failing to explore the full support of the posterior distribution of the clustering. In general, while MCMC convergence theorems hold as the number of iterations tend to infinity, any finite chain might suffer in representing the full support of the posterior distribution, as we observe here. Moreover, the mixing of each chain can be poor as well (i.e. it may take a long time to reach the stationary distribution from an arbitrary initialisation). In our empirical study, we find that using many short runs provide similar point and interval estimates to running a small number of long chains (figure 3), while being computationally less expensive (figure 4), and hence more convenient for our applications.

**Figure 3:**
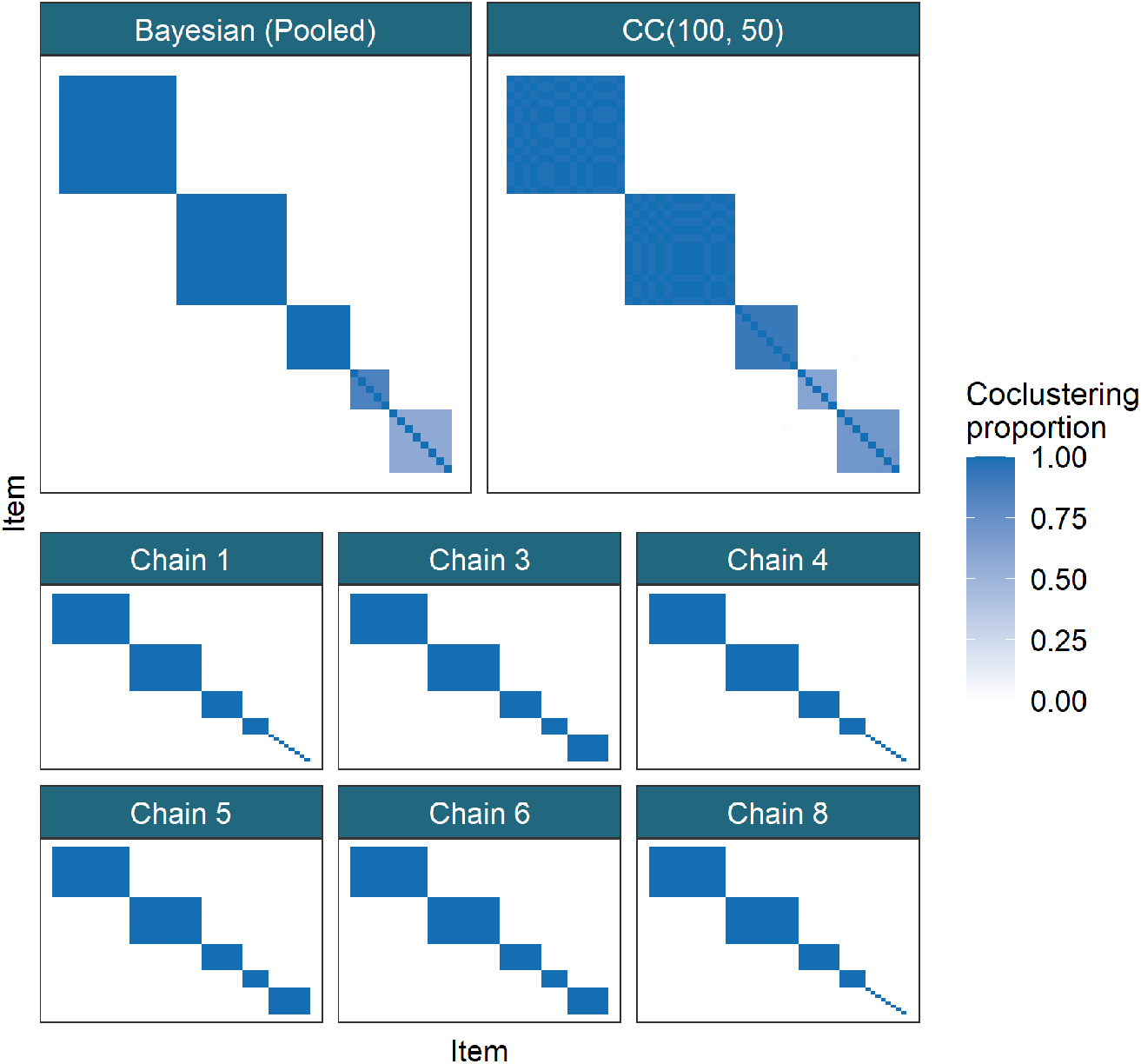
Comparison of similarity matrices from a dataset for the Small *N*, large *P* scenario. In each matrix, the (*i, j*)^*th*^ entry is the proportion of clusterings for which the *i*^*th*^ and *j*^*th*^ items co-clustered for the method in question. In the first row the PSM of the pooled Bayesian samples is compared to the CM for CC(100, 50), with a common ordering of rows and columns in both heatmaps. In the following rows, 6 of the long chains that passed the tests of convergence are shown.

**Figure 4:**
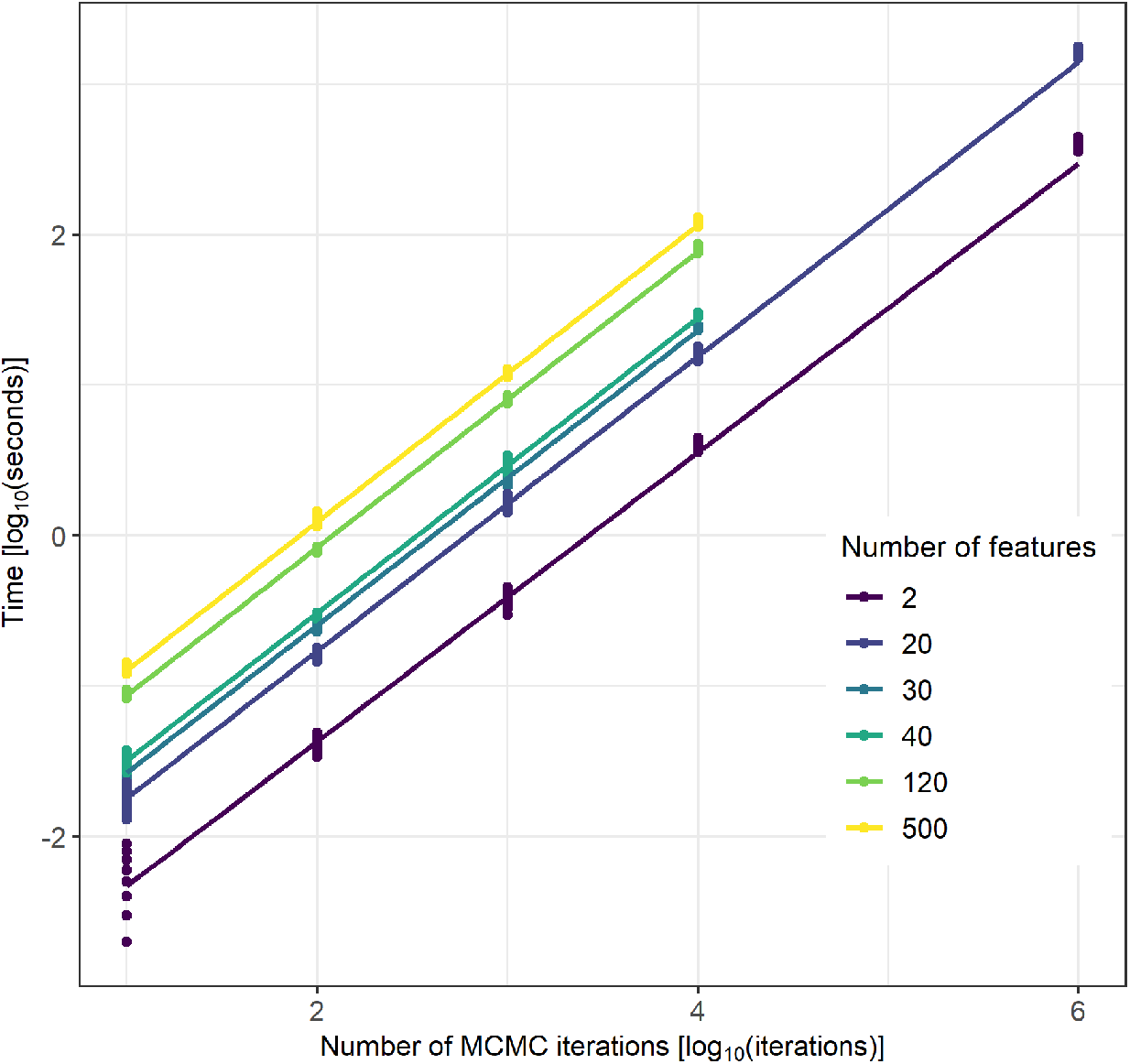
The time taken for different numbers of iterations of MCMC moves in log_10_(*seconds*). The relationship between chain length, *D*, and the time taken is linear (the slope is approximately 1 on the log_10_ scale), with a change of intercept for different dimensions. The runtime of each Markov chain was recorded using the terminal command `time`, measured in milliseconds.

Figure 4 shows that chain length is directly proportional to the time taken for the chain to run. This means that using an ensemble of shorter chains, as in consensus clustering, can offer large reductions in the time cost of analysis when a parallel environment is available compared to standard Bayesian inference. Even on a laptop of 8 cores running an ensemble of 1,000 chains of length 1,000 will require approximately half as much time as running 10 chains of length 100,000 due to parallelisation, and the potential benefits are far greater when using a large computing cluster.

Additional results for these and other simulations are in section 4.4 of the Supplementary Material.

### Multi-omics analysis of the cell cycle in budding yeast

We use the stopping rule proposed in to determine our ensemble depth and width. In figure 5, we see that the change in the consensus matrices from increasing the ensemble depth and width is diminishing in keeping with results in the simulations. We see no strong improvement after *D* = 6, 000 and increasing the number of learners from 500 to 1,000 has small effect. We therefore use the largest ensemble available, a depth *D* = 10001 and width *W* = 1000, believing this ensemble is stable (additional evidence in section 5.1 of the Supplementary Material).

**Figure 5:**
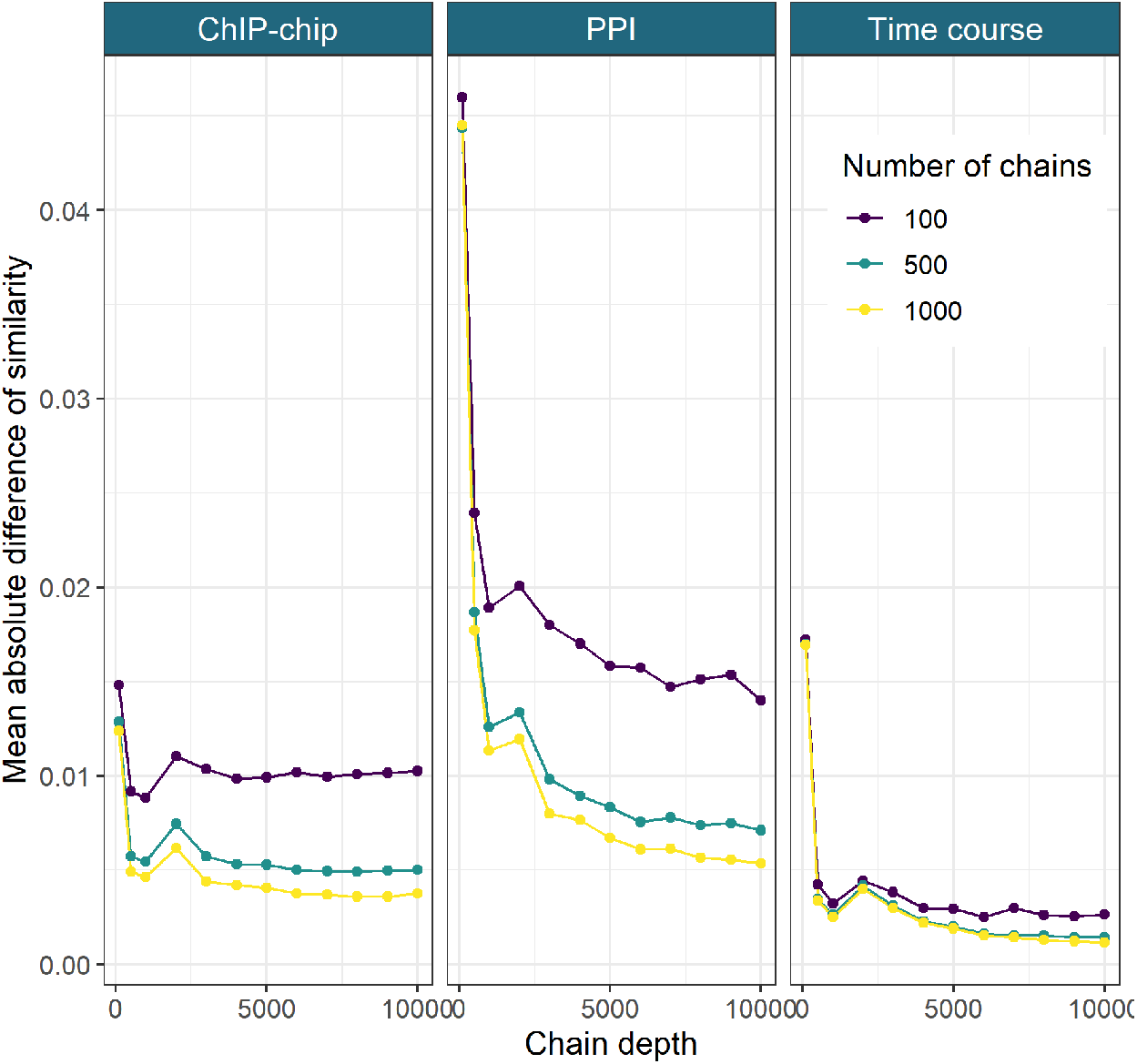
The mean absolute difference between the sequential Consensus matrices. For a set of chain lengths, *D*^′^ = {*d*_1_, …, *d*_*I*_} and number of chains, *W*^′^ = {*w*_1_, …, *w*_*J*_ }, we take the mean of the absolute difference between the consensus matrix for (*d*_*i*_, *w*_*j*_) and (*d*_*i−*1_, *w*_*j*_) (here *D*^′^ = {101, 501, 1001, 2001, …, 10001} and *W*^′^ = {100, 500, 1000}).

We focus upon the genes that tend to have the same cluster label across multiple datasets. More formally, we analyse the clustering structure among genes for which 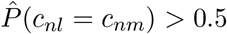, where *c*_*nl*_ denotes the cluster label of gene *n* in dataset *l*. In our analysis it is the signal shared across the time course and ChIP-chip datasets that is strongest, with 261 genes (nearly half of the genes present) in this pairing tending to have a common label, whereas only 56 genes have a common label across all three datasets. Thus, we focus upon this pairing of datasets in the results of the analysis performed using all three datasets. We show the gene expression and regulatory proteins of these genes separated by their cluster in figure 6. In figure 6, the clusters in the time series data have tight, unique signatures (having different periods, amplitudes, or both) and in the ChIP-chip data clusters are defined by a small number of well-studied transcription factors (**TFs**) (see table 2 of the Supplementary Material for details of these TFs, many of which are well known to regulate cell cycle expression, 84).

**Figure 6:**
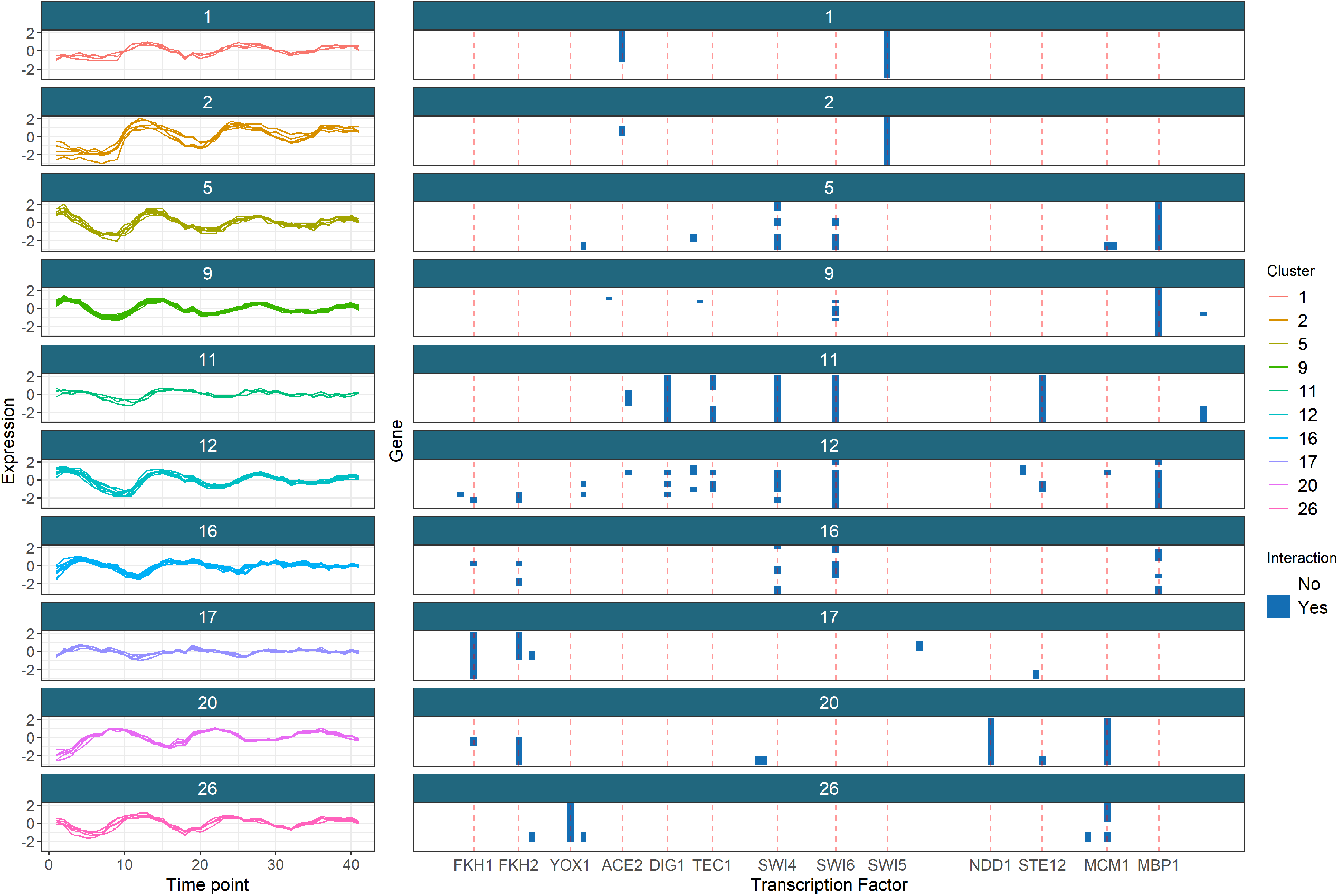
The gene clusters which tend to have a common label across the time course and ChIP-chip datasets, shown in these datasets. We include only the clusters with more than one member and more than half the members having some interactions in the ChIP-chip data. Red lines for the most common transcription factors are included.

As an example, we briefly analyse clusters 9 and 16 in greater depth. Cluster 9 has strong association with MBP1 and some interactions with SWI6, as can be seen in figure 6. The Mbp1-Swi6p complex, MBF, is associated with DNA replication (85). The first time point, 0 minutes, in the time course data is at the START checkpoint, or the G1/S transition. The members of cluster 9 begin highly expressed at this point before quickly dropping in expression (in the first of the 3 cell cycles). This suggests that many transcripts are produced immediately in advance of S-phase, and thus are required for the first stages of DNA synthesis. These genes’ descriptions (found using `org.Sc.sgd.db`, 86, and shown in table 3 of the Supplementary Material) support this hypothesis, as many of the members are associated with DNA replication, repair and/or recombination. Additionally, *TOF1, MRC1* and *RAD53*, members of the replication checkpoint (87, 88) emerge in the cluster as do members of the cohesin complex. Cohesin is associated with sister chromatid cohesion which is established during the S-phase of the cell cycle (89) and also contributes to transcription regulation, DNA repair, chromosome condensation, homolog pairing (90), fitting the theme of cluster 9.

Cluster 16 appears to be a cluster of S-phase genes, consisting of *GAS3, NRM1* and *PDS1* and the genes encoding the histones H1, H2A, H2B, H3 and H4. Histones are the chief protein components of chromatin (91) and are important contributors to gene regulation (92). They are known to peak in expression in S-phase (81), which matches the first peak of this cluster early in the time series. Of the other members, *NRM1* is a transcriptional co-repressor of MBF-regulated gene expression acting at the transition from G1 to S-phase (93, 94). Pds1p binds to and inhibits the Esp1 class of sister separating proteins, preventing sister chromatids separation before M-phase (95, 89). *GAS3*, is not well studied. It interacts with *SMT3* which regulates chromatid cohesion, chromosome segregation and DNA replication (among other things). Chromatid cohesion ensures the faithful segregation of chromosomes in mitosis and in both meiotic divisions and is instantiated in S-phase (89). These results, along with the very similar expression profile to the histone genes in the time course data, suggest that *GAS3* may be more directly involved in DNA replication or chromatid cohesion than is currently believed.

We attempt to perform a similar analysis using traditional Bayesian inference of MDI, but after 36 hours of runtime there is no consistency or convergence across chains. We use the Geweke statistic and 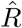 to reduce to the five best behaved chains (none of which appear to be converged, see section 5.2 of the Supplementary Material for details). If we then compare the distribution of sampled values for the *ϕ* parameters for these long chains, the final ensemble used (D = 10001, W = 1000) and the pooled samples from the 5 long chains, then we see that the distribution of the pooled samples from the long chains (which might be believed to sampling different parts of the posterior distribution) is closer in appearance to the distributions sampled by the consensus clustering than to any single chain (figure 7). Further disagreement between chains is shown in the Gene Ontology term over-representation analysis in section 5.3 of the Supplementary Material.

**Figure 7:**
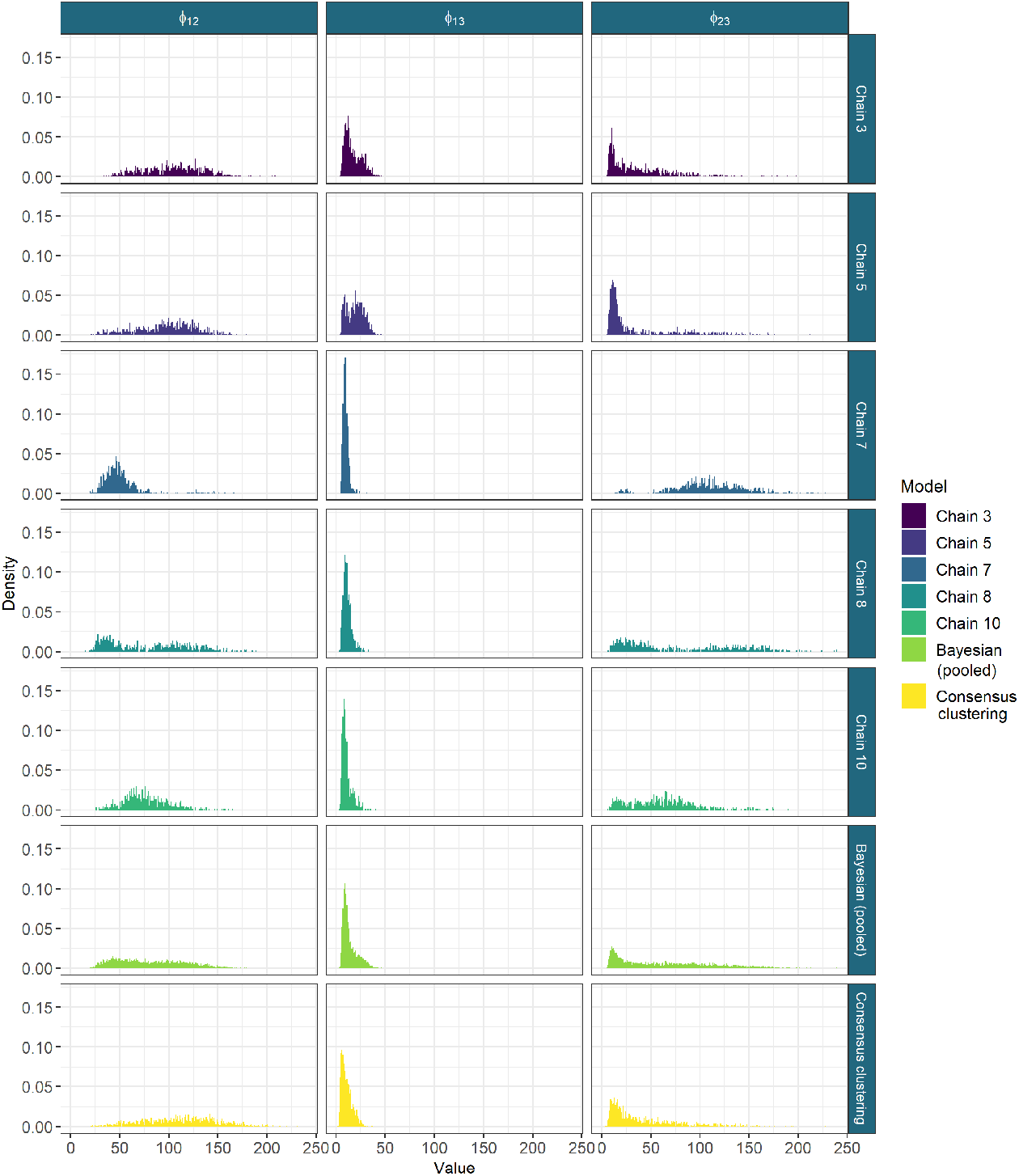
The sampled values for the *ϕ* parameters from the long chains, their pooled samples and the consensus using 1000 chains of depth 10,001. The long chains display a variety of behaviours. Across chains there is no clear consensus on the nature of the posterior distribution. The samples from any single chain are not particularly close to the behaviour of the pooled samples across all three parameters. It is the consensus clustering that most approaches this pooled behaviour.

## Discussion

Our proposed method has demonstrated good performance on simulation studies, uncovering the generating structure in many cases and performing comparably to `Mclust` and long chains in many scenarios. We saw that when the chains are sufficiently deep that the ensemble approximates Bayesian inference, as shown by the similarity between the PSMs and the CM in the 2D scenario where the individual chains do not become trapped in a single mode. We have shown cases where many short runs are computationally less expensive than one long chain and give meaningful point and interval estimates; estimates that are very similar to those from the limiting case of a Markov chain. Thus if individual chains are suffering from mixing problems or are too computationally expensive to run, consensus clustering may provide a viable option. We also showed that the ensemble of short chains is more robust to irrelevant features than `Mclust`.

We proposed a method of assessing ensemble stability and deciding upon ensemble size which we used when performing an integrative analysis of yeast cell cycle data using MDI, an extension of Bayesian mixture models that jointly models multiple datasets. We uncovered many genes with shared signal across several datasets and explored the meaning of some of the inferred clusters using data external to the analysis. We found biologically meaningful results as well as signal for possibly novel biology. We also showed that individual chains for the existing implementation of MDI do not converge in a practical length of time, having run 10 chains for 36 hours with no consistent behaviour across chains. This means that Bayesian inference of the MDI model is not practical on this dataset with the software currently available.

However, consensus clustering does lose the theoretical framework of true Bayesian inference. We attempt to mitigate this with our assessment of stability in the ensemble, but this diagnosis is heuristic and subjective, and while there is empirical evidence for its success, it lacks the formal results for the tests of model convergence for Bayesian inference.

More generally, we have benchmarked the use of an ensemble of Bayesian mixture models, showing that this approach can infer meaningful clusterings and overcomes the problem of multi-modality in the likelihood surface even in high dimensions, thereby providing more stable clusterings than individual long chains that are prone to becoming trapped in individual modes. We also show that the ensemble can be significantly quicker to run. In our multi-omics study we have demonstrated that the method can be applied as a wrapper to more complex Bayesian clustering methods using existing implementations and that this provides meaningful results even when individual chains fail to converge. This enables greater application of complex Bayesian clustering methods without requiring re-implementation using more clever MCMC methods, a process that would involve a significant investment of human time.

We expect that researchers interested in applying some of the Bayesian integrative clustering models such as MDI and Clusternomics (32) will be enabled to do so, as consensus clustering overcomes some of the unwieldiness of existing implementations of these complex models. More generally, we expect that our method will be useful to researchers performing cluster analysis of high-dimensional data where the runtime of MCMC methods becomes too onerous and multi-modality is more likely to be present.

## Abbreviations

ARI: Adjusted Rand Index
ChIP-chip: Chromatin immunoprecipitation followed by microarray hybridization
CM: Consensus Matrix
MCMC: Markov chain Monte Carlo
MDI: Multiple Dataset Integration
PCA: Principal Component Analysis
PPI: Protein-Protein Interaction
PSM: Posterior Similarity Matrix
SSE: Sum of Squared Errors
TF: Transcription Factor

## Availability of data and materials

The code and datasets supporting the conclusions of this article are available in the github repository, https://github.com/stcolema/ConsensusClusteringForBayesianMixtureModels.

## Competing interests

The authors declare that they have no competing interests.

## Funding

This work was funded by the MRC (MC UU 00002/4, MC UU 00002/13) and supported by the NIHR Cambridge Biomedical Research Centre (BRC-1215-20014). The views expressed are those of the author(s) and not necessarily those of the NHS, the NIHR or the Department of Health and Social Care. This research was funded in whole, or in part, by the Wellcome Trust [WT107881]. For the purpose of Open Access, the author has applied a CC BY public copyright licence to any Author Accepted Manuscript version arising from this submission.

## Authors’ contributions

SC designed the simulation study with contributions from PK and CW, performed the analyses and wrote the manuscript. PK and CW provided an equal contribution of joint supervision, directing the research and provided suggestions such as the stopping rule. All contributed to interpreting the results of the analyses. All authors revised and approved the final manuscript.

## Additional Files

### Additional file 1 — Supplementary materials

Additional relevant theory, background and results. This includes some more formal definitions, details of Bayesian mixture models and MDI, the general consensus clustering algorithm, additional simulations and the generating algorithm used, steps in assessing Bayesian model convergence in both the simulated datasets and yeast analysis, a table of the transcription factors that define the clustering in the ChIP-chip dataset, a table of the gene descriptions for some of the clusters that emerge across the time course and ChIP-chip datasets and Gene Ontology term over-representation analysis of the clusterings from the yeast datasets. (PDF, 10MB)

## 1 Definitions

### Definition 1

**(Coclustering matrix)** *The* coclustering matrix *describes a clustering or partition in a binary matrix with the* (*i, j*)^*th*^ *entry indicating if items i and j are allocated to the same cluster*.

### Definition 2

**(Consensus matrix)** *Given W clusterings for a dataset of N items, c*_*s*_ = (*c*_*s*1_, …, *c*_*sN*_), *the* consensus matrix *is a N* ×*N matrix where the* (*i, j*)^*th*^ *entry records the proportions of clusterings for which items i and j are allocated the same label. More formally, it is the matrix* ℂ *such that*

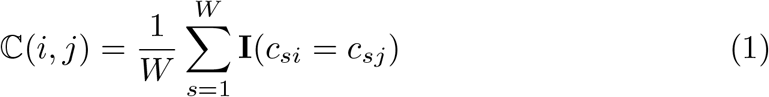

*where 𝕀* (·) *is the indicator function taking a value of 1 if the argument is true and 0 otherwise*.

### Definition 3

**(Posterior similarity matrix)** *A* consensus matrix *for which all the clusterings are generated from a converged Markov chain for some Bayesian clustering model. Sometimes abbreviated to* PSM.

### Definition 4

**(Partition *or* Clustering)** *For a dataset of items X* = (*x*_1_, …, *x*_*N*_), *a* partition *or* clustering *is a set of disjoint sets covering X, normally indicated by a N-vector of integers indicating which set each item is associated with. Note that these labels only have meaning relative to each other, they are symbolic. Each set within the clustering is referred to as a* cluster.

## 2 The models

### 2.1 Individual dataset

In the simulations (see section 4) where individual datasets are modelled a *Bayesian mixture model* is used. We write the basic mixture model for independent items *X* = (*x*_1_, …, *x*_*N*_) as

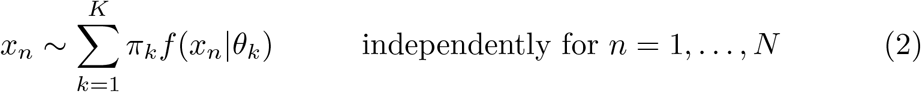

where *f* (· | *θ*) is some family of densities parametrised by *θ*. A common choice is the Gaussian density function, with *θ* = (*µ, σ*^2^) (as in our simulation study). *K*, the number of subgroups in the population, 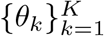, the component parameters, and *π* = (*π*_1_, …, *π*_*K*_), the component weights are the objects to be inferred. In the context of *clustering*, such a model arises due to the belief that the population from which the random sample under analysis has been drawn consists of *K* unknown groups proportional to *π*. In this setting it is natural to include a latent *allocation variable, c* = (*c*_1_, …, *c*_*N*_), to indicate which group each item is drawn from, with each non-empty component of the mixture corresponds to a cluster. The model is

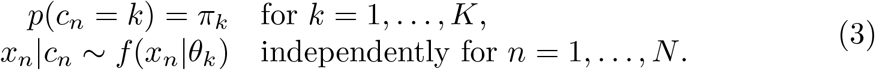

The joint model can then be written

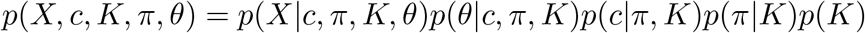

We assume conditional independence between certain parameters such that the model reduces to

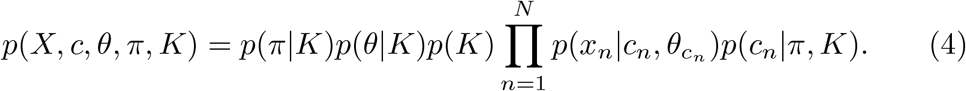

Additional flexibility is provided by the inclusion of hyperparameters on the priors for *π* and *θ*, denoted *α* and *η* respectively. In our context where *θ* = (*µ, σ*^2^), we use

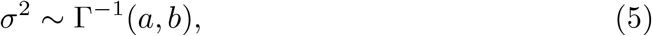

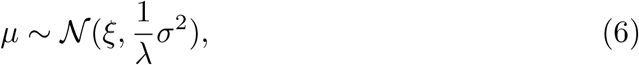

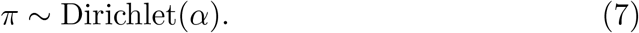

**Figure 1:**
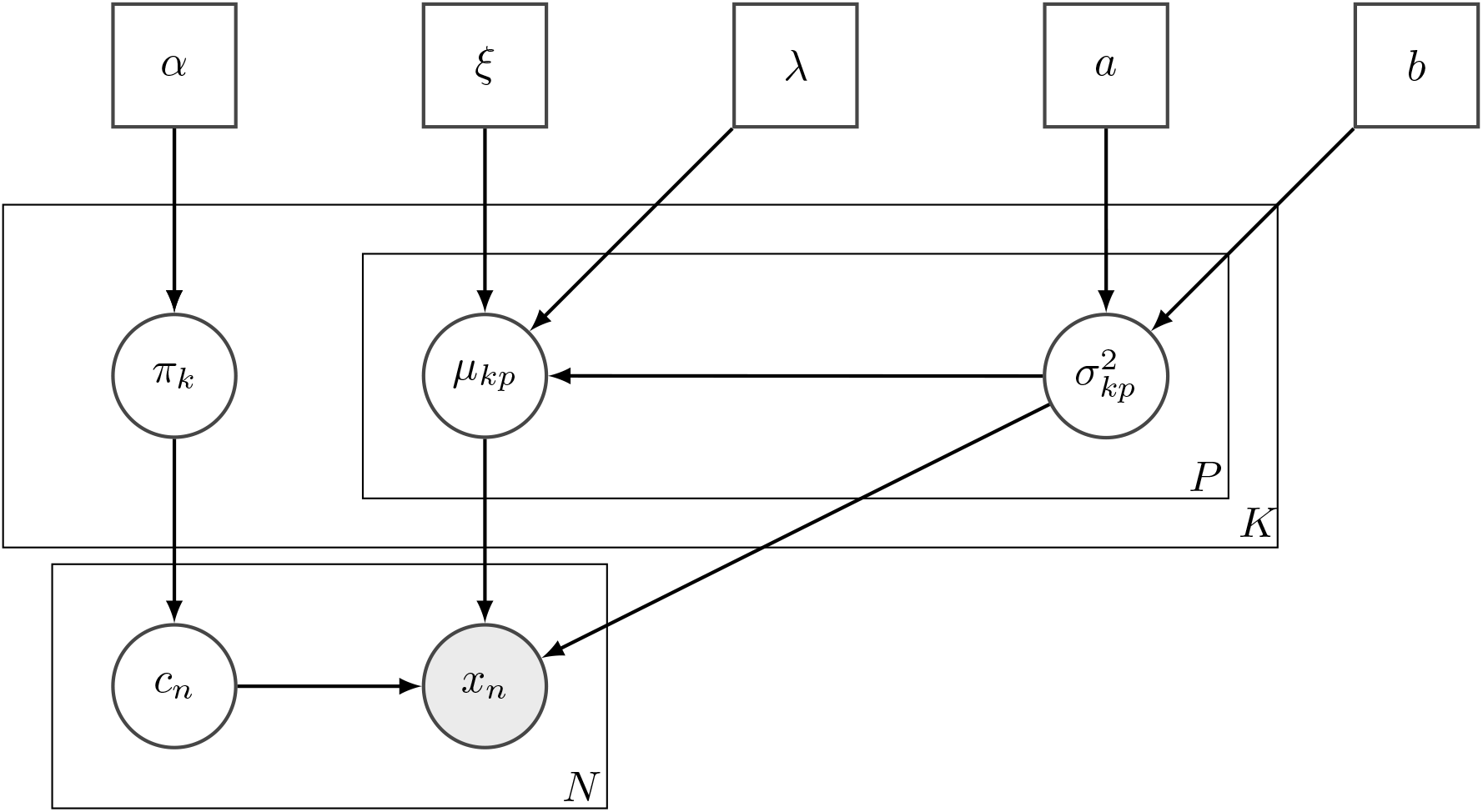
Directed acyclic graph for a mixture of Gaussians with independent features, as used in the simulation study.

The directed acyclic graph (**DAG**) for this model is shown in figure 1. The value of the hyperparameters we use are

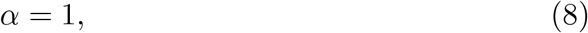

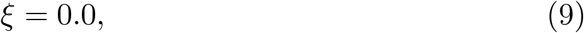

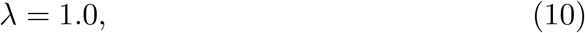

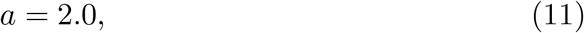

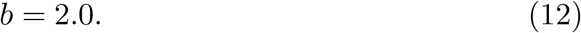

### 2.2 Integrative clustering

We are interested in the use of consensus clustering for integrative methods. We used Multiple Dataset Integration (**MDI**, Kirk et al., 2012) as an example of a Bayesian integrative clustering method. MDI models dataset specific clusterings, in contrast to, for example, Clusternomics (Gabasova et al., 2017) in which a *global clustering* is inferred.

The defining aspect of MDI is the prior on the allocation of the *n*^*th*^ item across the *L* datasets

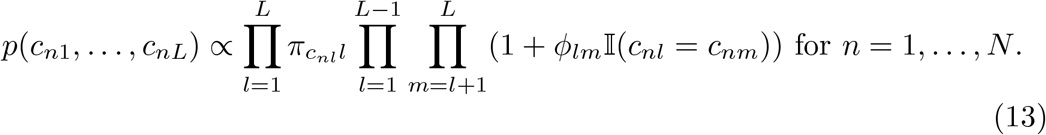

*ϕ*_*lm*_ is the parameter defined by the similarity of the clusterings for the *l*^*th*^ and *m*^*th*^ datasets and is also sampled in each iteration. As *ϕ*_*lm*_ increases more mass is placed on the common partition for these datasets. Conversely, in the limit *ϕ*_*lm*_ →0 we have independent mixture models. In other words, MDI allows datasets with similar clustering of the items to inform the clustering in each other more strongly than the clustering for an unrelated dataset. The DAG for this model for three datasets is shown in figure 2.

## 3 Consensus clustering

Consensus clustering as described by Monti et al. (2003) applies *W* independent runs of the underlying clustering algorithm to perturbed versions of the dataset and combines the *W* final partitions in a *consensus matrix* which can be used to infer a final clustering. An outline of this is described in algorithm 1.

The consensus matrix is a symmetric matrix with the (*i, j*)^*th*^ entry being the proportions of model runs for which the *i*^*th*^ and *j*^*th*^ items are clustered together.

### Algorithm 1

Consensus clustering algorithm

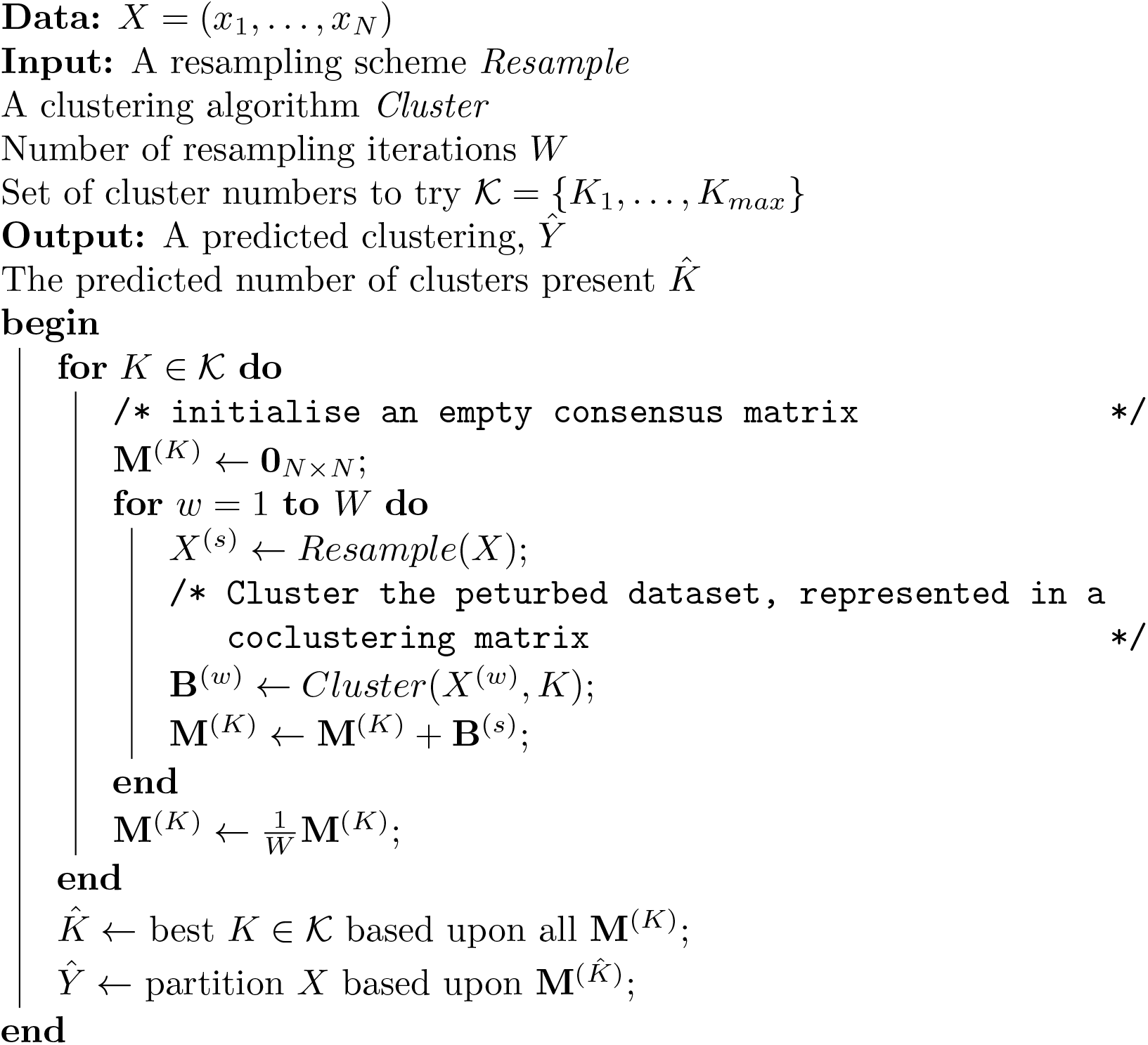

To partition *X* based upon the consensus matrix, we use the R function `maxpear` (Fritsch, 2012). `maxpear` uses a sample average clustering, inferring this by maximising the quantity

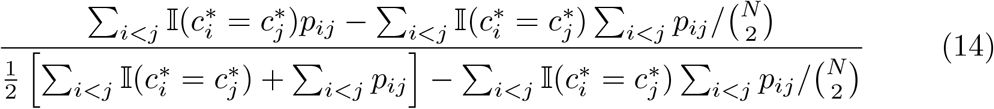

where *p*_*ij*_ is the (*i, j*)^*th*^ entry of the consensus matrix (Fritsch et al., 2009).

**Figure 2:**
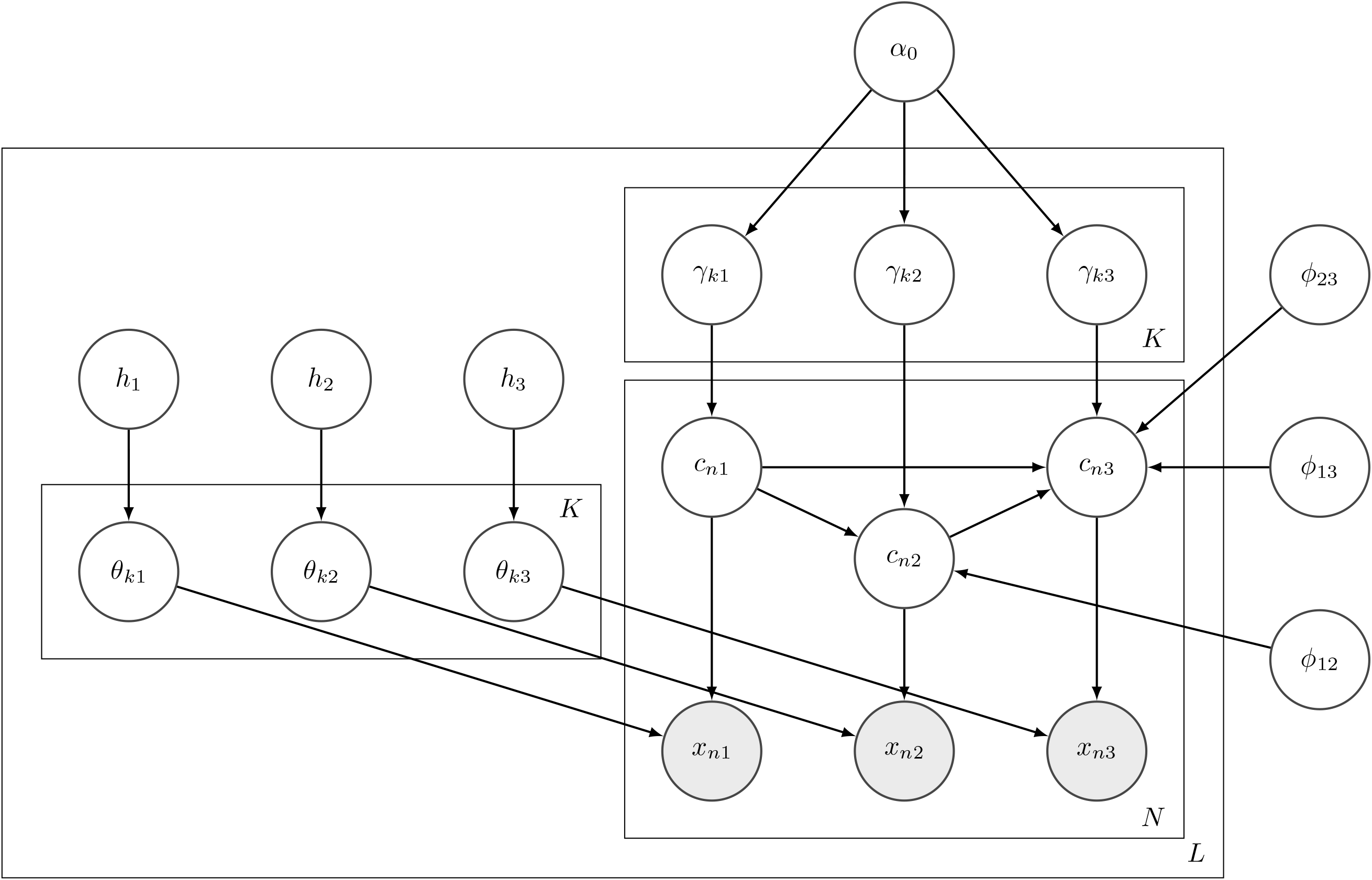
Directed acyclic graph for the Multiple Dataset Integration model for *L* = 3 datasets. *h*_*l*_ is the choice of hyperpriors for the *l*^*th*^ dataset.

## 4 Simulated data

### 4.1 Scenario description

We defined 12 scenarios to simulate data within to test consensus clustering and to compare it to some alternative tools. Table 1 describes the parameters defining these scenarios and algorithm 2 describes how individual simulations were generated.

**Table S1:**
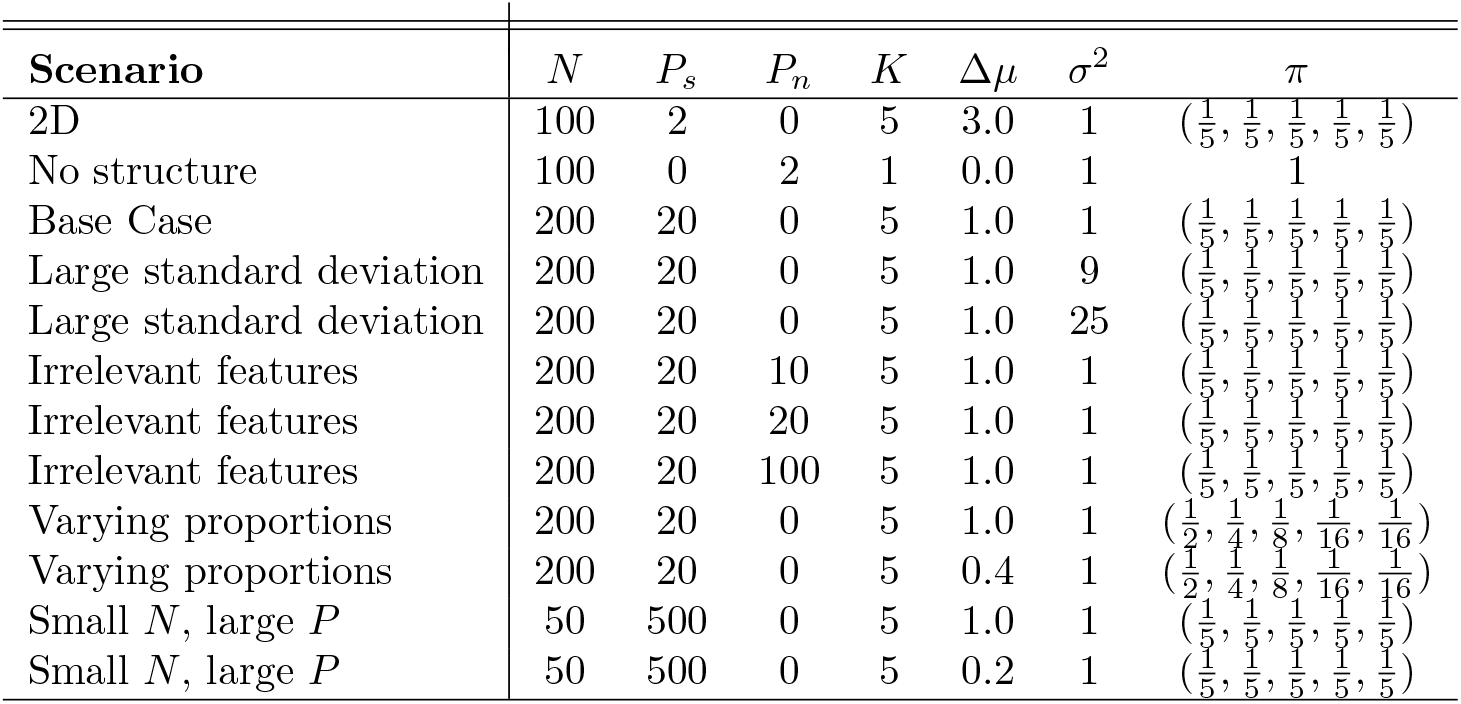
Parameters defining the simulation scenarios as used in generating data and labels.

We intend the scenarios to test different aspects of real data or to benchmark performance for comparison in the more challenging situations.

- *2D* : a low dimensional scenario within which we expected `Mclust` to perform well and the long chains to converge and explore the full support of the posterior distribution.
- *No structure*: we included this scenario to reassure fears that consensus clustering has a predilection to finding clusters where none exist (S, enbabaoğlu et al., 2014a,b).
- *Base case*: highly informative datasets within which we expected methods to find the true generating labels quite easily. We included this scenario to benchmark the others that are variations of this setting.

#### Algorithm 2

Data generation for a mixture of Gaussian with independent features. This algorithm is implemented in the `generateSimulationDataset` function from the `mdiHelpR` package available at www.github.com/stcolema/mdiHelpR.

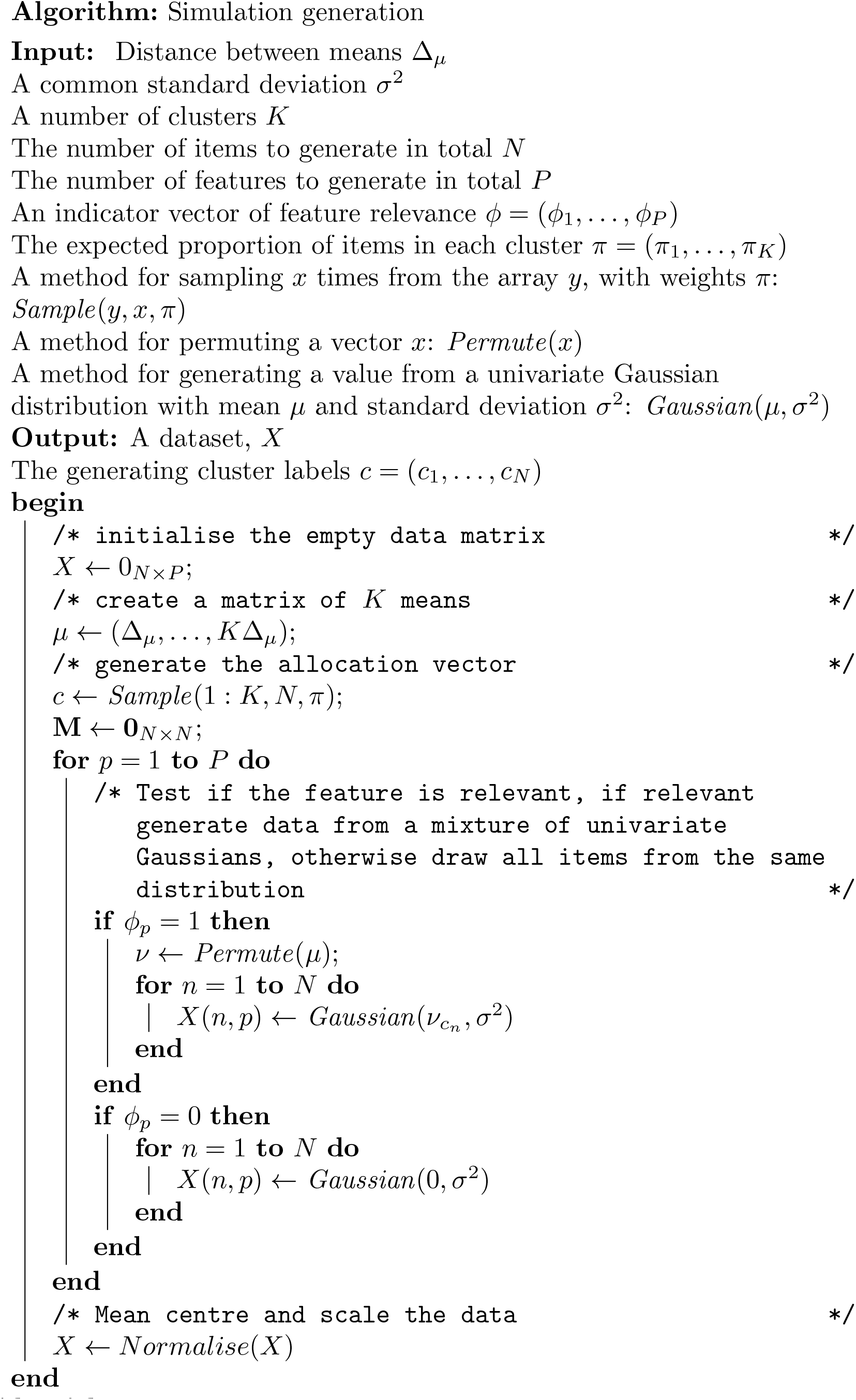

- *Large standard deviation*: these two scenarios investigated the degree of distinction required between clusters for the methods to uncover their structure.
- *Irrelevant features*: we included these scenarios to investigate how robust the methods are to irrelevant features.
- *Varying proportions* : these scenarios investigated how well each method uncovers clusters when the clusters have significantly different membership counts.
- *Small N, large P* : an investigation of behaviour when the number of features is far greater than the number of items.

### 4.2 Mclust

We called `Mclust` using the default settings and a range of inputs for the choice of *K*. We used 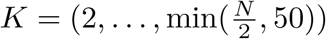 to mirror the choice of *K*_*max*_ = 50 used for the overfitted mixture models (the default in the software we used), with the bound of 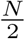 to avoid fitting 50 clusters in the *Small N, large P* scenario where *N* = 50 = *K*_*max*_. In the *No structure* scenarios we extended to range to *K* = (1 …, 50) to include the correct structure as an option. The model choice was performed using the Bayesian Information Criterion (Schwarz et al., 1978, as implemented in `Mclust`). `Mclust` tries different covariance matrices and thus the model choice is not just between different values of *K*.

### 4.3 Bayesian analysis

We use the implementation of Bayesian mixture models in C++ provided by Mason et al. (2016). Rather than directly using a Dirichlet process (Ferguson, 1973) to infer the number of clusters or a mixture that grows and shrinks (Richardson and Green, 1997), this implementation follows the logic of Rousseau and Mengersen (2011) and Van Havre et al. (2015) using an overfitted mixture model to approximate a Dirichlet process. In overfitted mixture models, the number of components, *K*_*max*_, included in the model is set to number far larger than the true number of clusters, *K*, present.

For each simulation we ran 10 chains for 1 million iterations, keeping every thousandth sample. We discarded the first 10,000 iterations to account for burn-in bias, leaving 990 samples per chain. To check if the chains were converged we used

- the Geweke convergence diagnostic (Geweke et al., 1991) to investigate within-chain stationarity, and
- the potential scale reduction factor (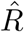, Gelman et al., 1992) and the Vats-Knudson extension (*stable* 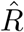, Vats and Knudson, 2018) to check across-chain convergence.

The Geweke convergence diagnostic is a standard Z-score; it compares the sample mean of two sets of samples (in this case buckets of samples from the first half of the samples to the sample mean of the entire second half of samples). It is calculated under the assumption that the two parts of the chain are asymptotically independent and if this assumption holds (i.e. the chain is sampling the same distribution in both samples) than the scores are expected to be standard normally distributed. If a chain’s Geweke convergence diagnostic passed a Shapiro-Wilks test for normality (Shapiro and Wilk, 1965) (based upon a threshold of 0.05), we considered it to have achieved stationarity and included it in the model performance analysis.

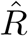 is expected to approach 1.0 if the set of chains are converged. Low 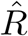 is not sufficient in itself to claim chain convergence, but values above 1.1 are clear evidence for a lack of convergence (Gelman et al., 2013). Vats and Knudson (2018) show that this threshold is significantly too high (1.01 being a better choice) and propose extensions to 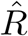 that enable a more formal rule for a threshold. We use their method as implemented in the R package `stableGR` (Knudson and Vats, 2020) as the final check of convergence. An example of the 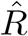 series across the 100 simulations for a scenario where chains are well-behaved is shown in figure 3.

We focused upon stationarity of the continuous variables as assesing convergence of the allocation labels is difficult due to label-switching. In our simulations the only recorded continuous variable is the concentration parameter of the Dirichlet distribution for the component weights.

We pooled the samples from the stationary chains and used these to form a PSM. This and the point estimate clustering found by applying the R function `maxpear`. In Bayesian inference, `maxpear` attempts to find the clustering that maximises the Adjusted Rand Index to the true clustering by using an approximation of the expected clustering under the posterior, 𝔼 (*c*| *X*), believing that this converges to the true clustering. A sample average clustering is used to approximate the expected clustering. This is estimated from the PSM by maximizing

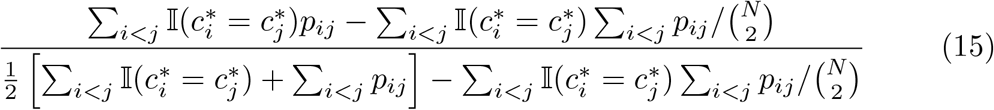

where *p*_*ij*_ is the (*i, j*)^*th*^ entry of the PSM (Fritsch et al., 2009). When the chain has converged this maximises the posterior expected ARI to the true clustering. There are three possibilities to consider the decision to pool the samples across chains under:

- The chains are converged and agree upon the distribution sampled (see figure 4 for an example).
- The chains are not in agreement upon the partition sampled, becoming trapped in different modes. However, a mode does dominate being the mode present in a majority of chains (see figure 5 for an example of this behaviour).

**Figure 3:**
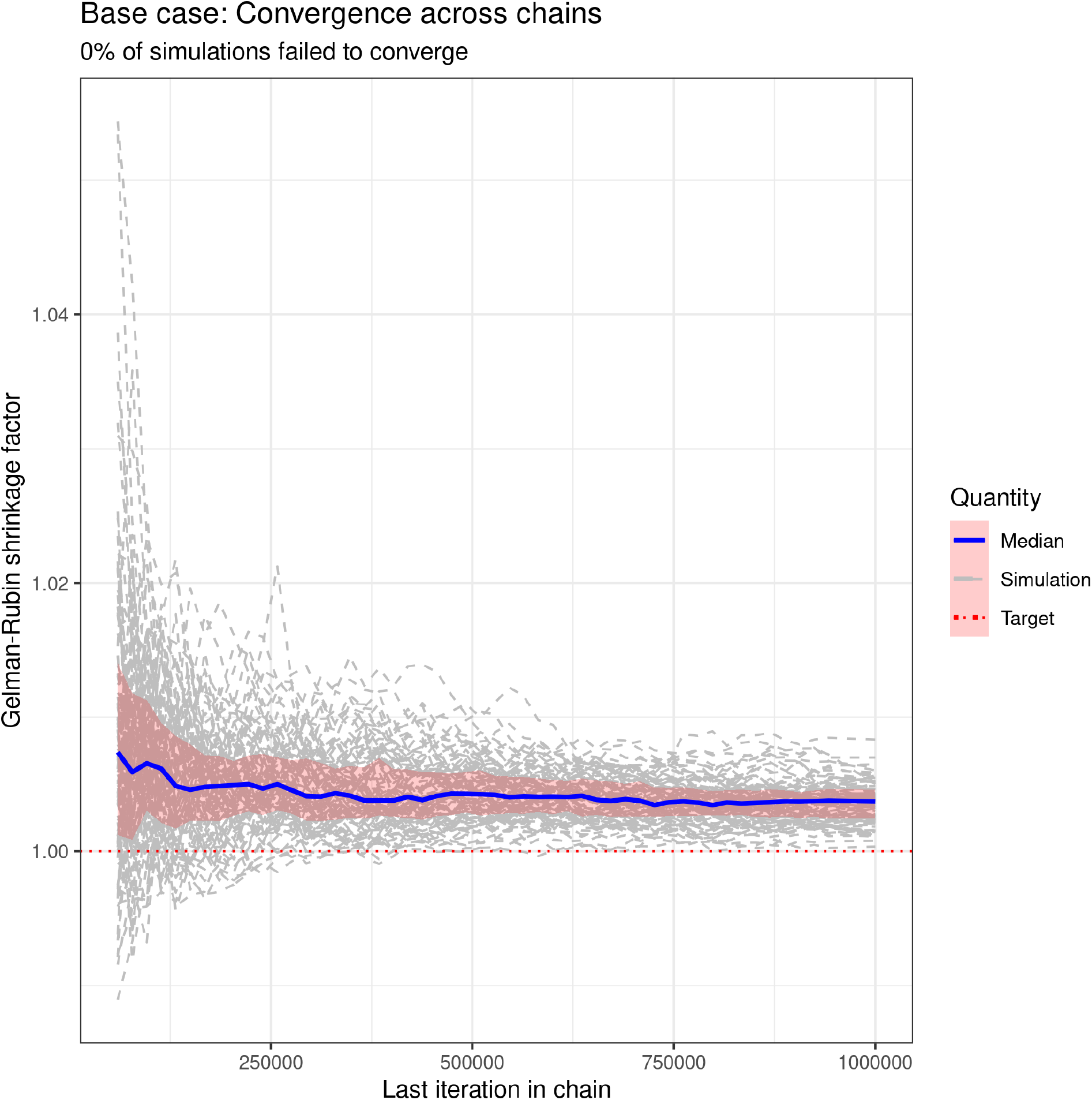
The 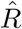 values for each simulation (in dotted grey), the median value and the interquartile range across simulations. One can see that 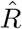 approaches 1.0, being below 1.01 for every simulation by the end of the chains. The “0% of simulations failed to converge” is a statement based upon the percentage of simulations which passed the test of stable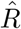.

**Figure 4:**
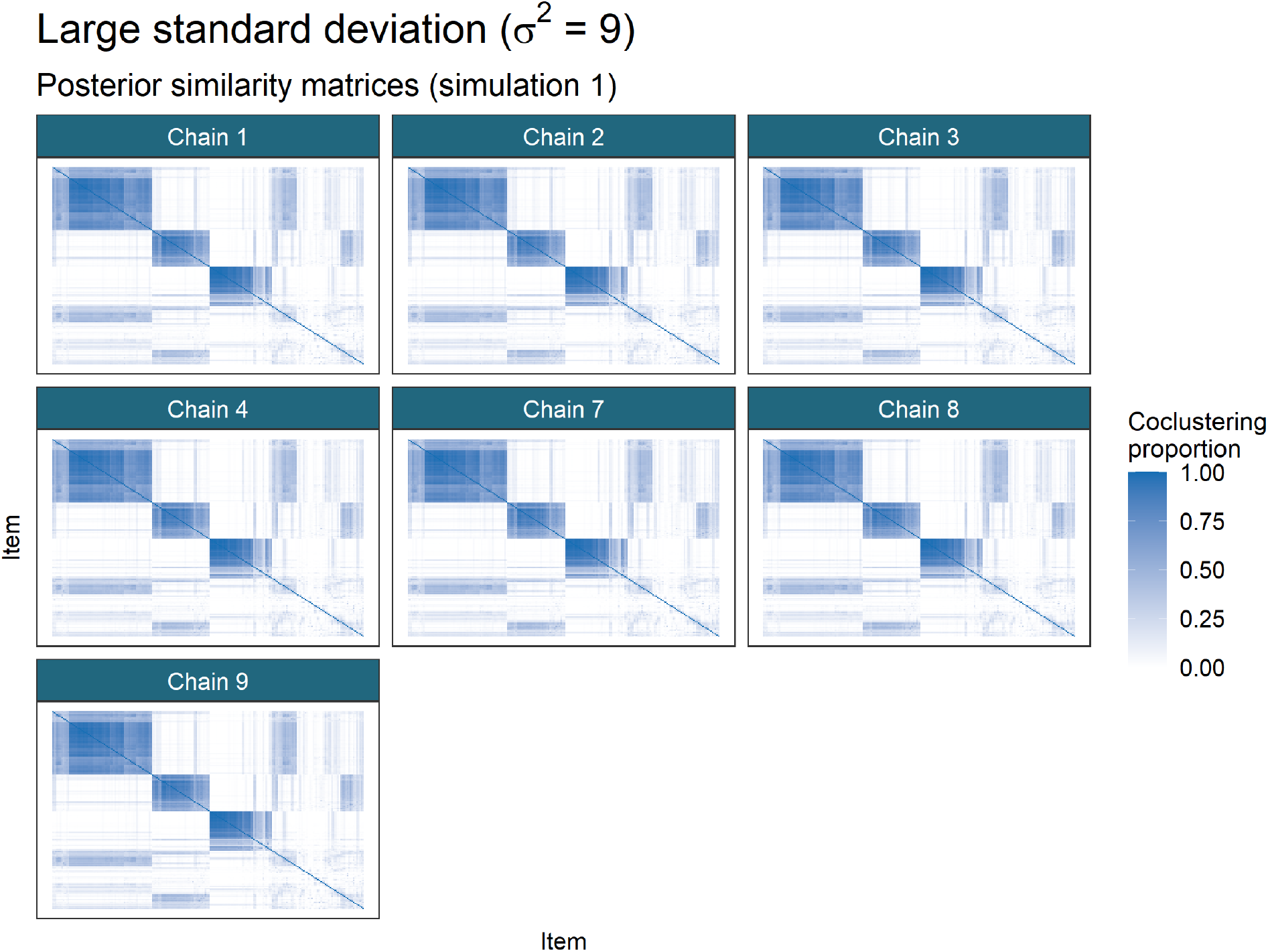
Posterior similarity matrices for the simulation generated using a random seed set to 1 for the first large standard deviation scenario from table This is an example of all stationary chains agreeing in a simulation (and thus pooling of samples is no different to using any choice of chain for the performance analysis). Ordering of rows and columns is defined by hierarchical clustering of the first matrix in the series, in this case that from Chain 1.
- The chains are not in agreement and no one mode dominates among chains (see figure 6 for an example of this behaviour).

**Figure 5:**
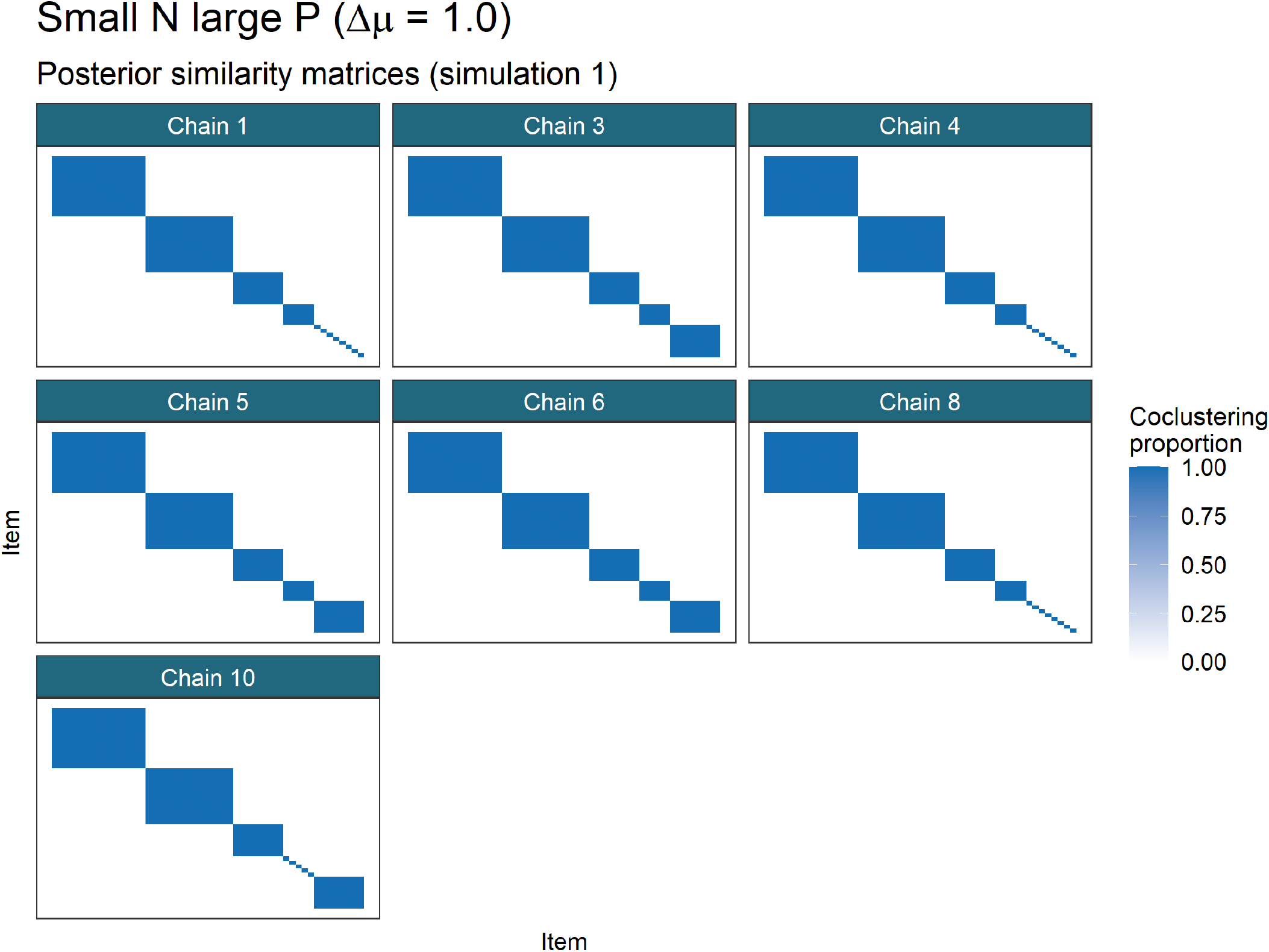
Posterior similarity matrices for the simulation generated using a random seed set to 1 for the first small *N*, large *P* scenario from table 1. This is an example of different chains becoming trapped in different modes, but one mode (which does represent the generating structure well) is dominant, being fully present in 3 of the 6 chains, with the two other modes present having significant overlap. Ordering of rows and columns is defined by hierarchical clustering of the first matrix in the series, in this case that from Chain 1.

**Figure 6:**
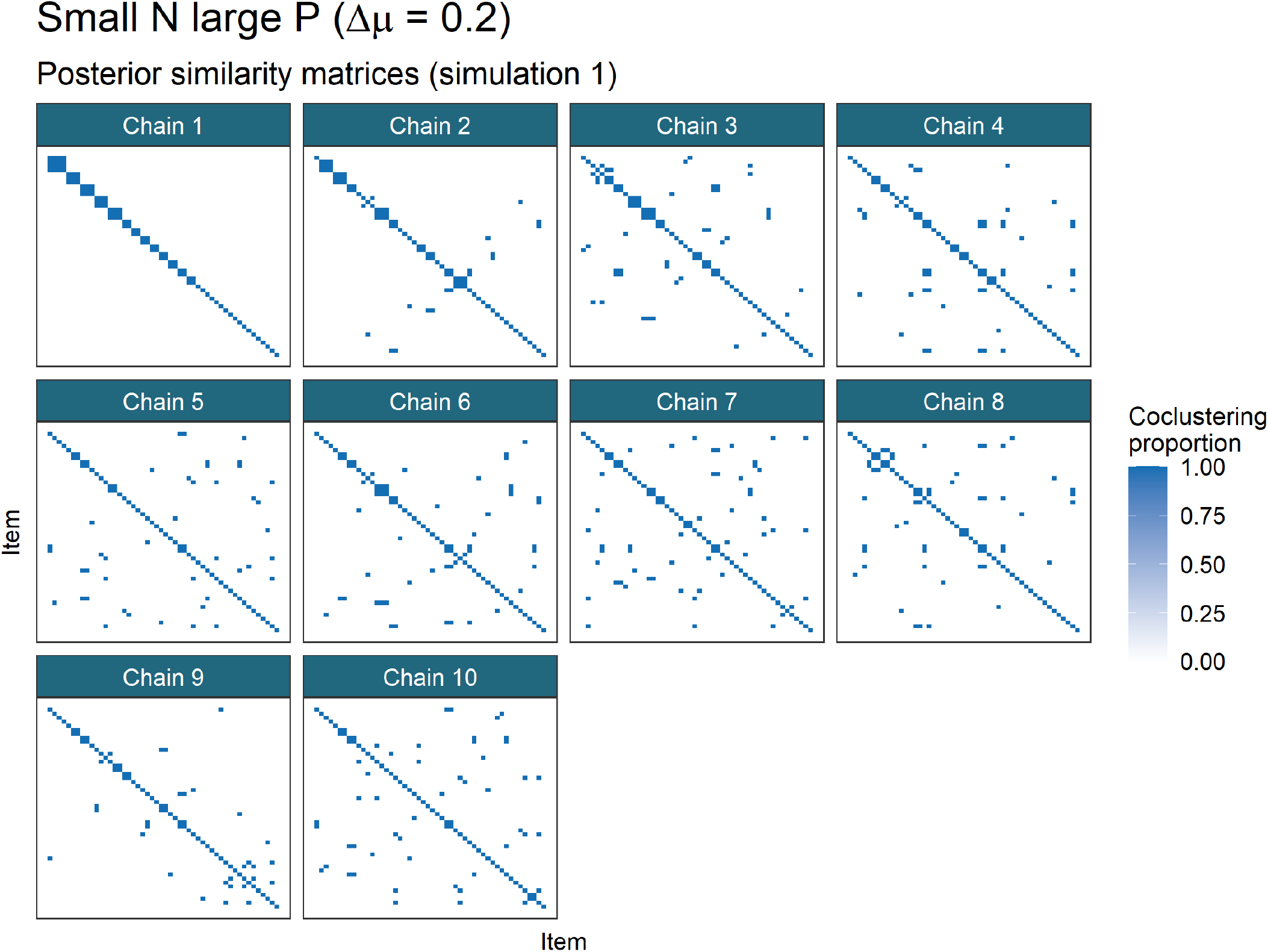
Posterior similarity matrices for the simulation generated using a random seed set to 1 for the second small *N*, large *P* scenario from table 1. This is an example of different chains becoming trapped in different modes with no mode being dominant. In this scenario each chain remains trapped in initialisation. Ordering of rows and columns is defined by hierarchical clustering of the first matrix in the series, in this case that from Chain 1.

**Figure 7:**
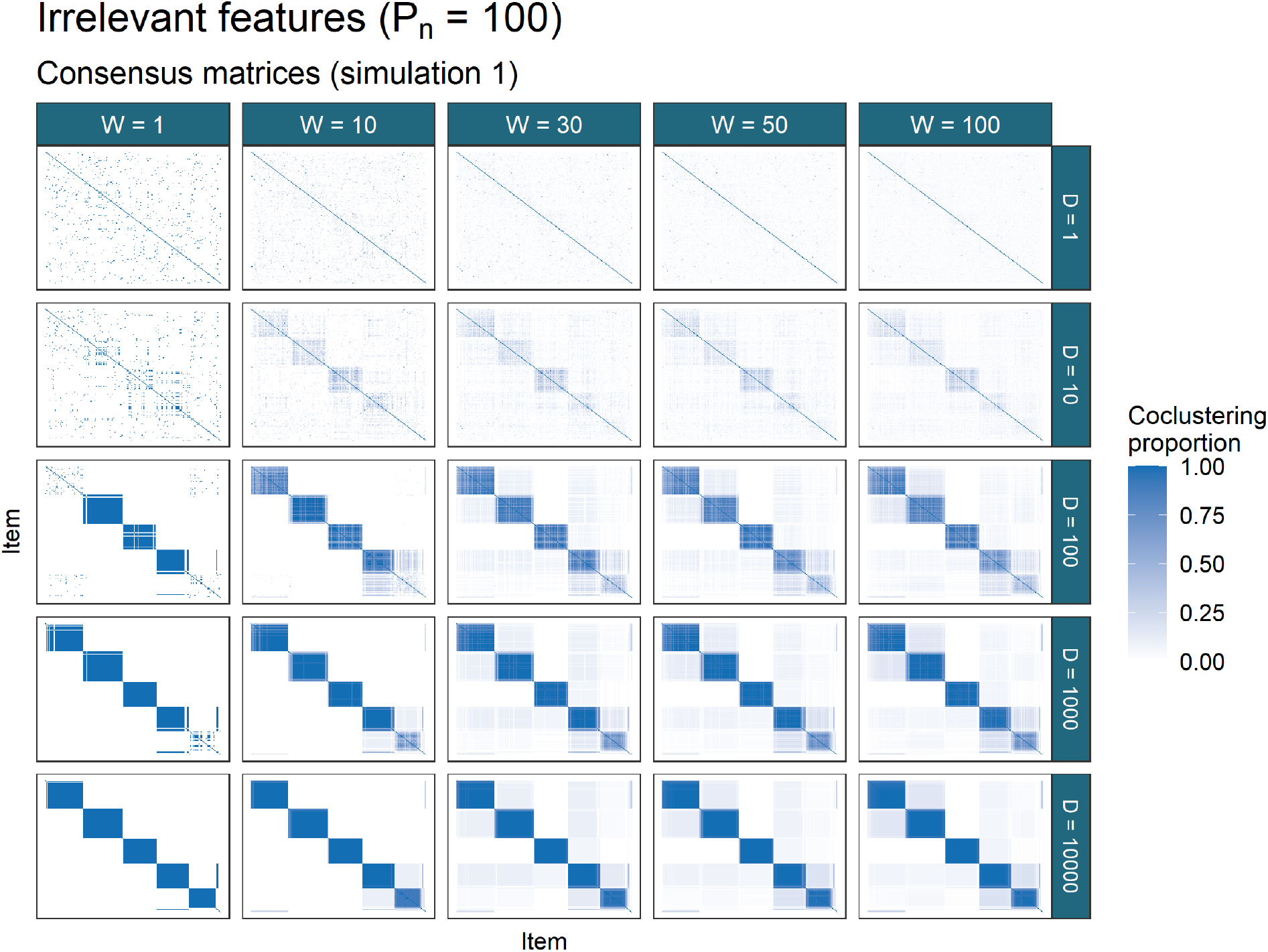
Consensus matrices for the simulation generated using a random seed set to 1 for the third irrelevant features scenario from table 1. *D* is the individual chain length and *W* is the number of chains used. In this example there are several modes present (as seen in the entries with values between 0 and 1) but one mode is clearly dominant (the 5 dark squares along the diagonal which correspond closely to the generating labels).

**Figure 8:**
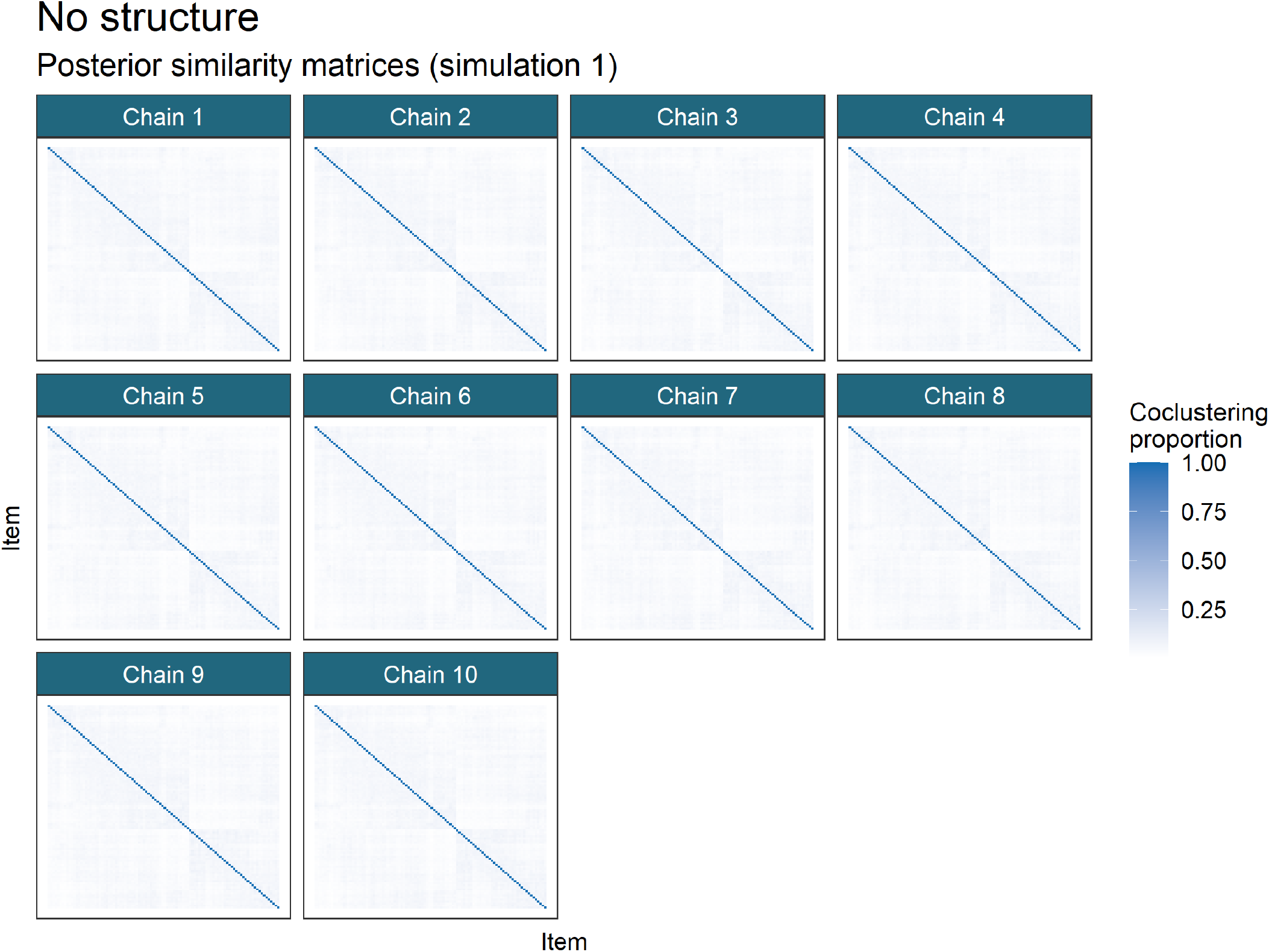
Posterior similarity matrices for simulation 1 of the *No structure* scenario. Each item is allocated to a singleton.

In the first case pooling has no effect upon the predicted clustering compared to using any one chain. In the second case it feels natural that one would use the mode that dominates. Pooling the samples effectively does this for the predictive performance of the method as the mode with the greatest number of samples across the chains dominates; however, the uncertainty for this mode is increased. In the third case the analysis is non-trivial and further thought, chains and samples would be required. In our simulations this case only arises in the most pathological form in the second *Large N, small P* scenario, where each chain remains trapped in the initial partition. The clustering inferred from any chain is not meaningful being a random clustering; thus the clustering predicted by pooling the PSMs is no more or less relevant as it too is random.

### 4.4 Consensus clustering analysis

We investigated a range of ensembles, using all combinations of chain depth, *D* = {1, 10, 100, 1000, 10000 }, and the number of chains, *W* = {1, 10, 30, 50, 100}. This gave a total of 25 different ensembles. A consensus matrix was constructed from the samples generated by each ensemble by finding the proportion of samples within which any pair of items are coclustered. An example of the Consensus matrices for each ensemble in a given simulation is shown in figure 7.

### 4.5 Model performance

The different models (Bayesian (pooled), `Mclust` and the 25 consensus clustering ensembles) were compared under their ability to uncover the generating clustering.

In figure 11 the ARI between the generating labels and the point estimate clustering from each method is shown. For two partitions *c*_1_, *c*_2_,

- *ARI*(*c*_1_, *c*_2_) = 1.0: a perfect match between the two partitions,
- *ARI*(*c*_1_, *c*_2_) = 0.0: *c*_1_ is no more similar to *c*_2_ than is expected for a random partition of the data.

In several scenarios `Mclust` performs the best under this metric (e.g. in the scenarios *2D, Small N, large P (*Δ*µ* = 0.2)). However when the number of irrelevant features is large `Mclust` performs less well (see *Irrelevant features (P*_*n*_ = 20) and *(P*_*n*_ = 100)) than the other methods. In the scenario that *P*_*n*_ = 100 failing to find structure is not inherently wrong as a majority of the features suggest that there are no subpopulations.

For the ensembles there are two parameters changing between each model, the iteration used to provide the clustering in the ensemble, *D*, and the number of chains (and hence samples) used, *W*. In many of the scenarios we find that the benefit of increasing *D* stabilises by approximately *D* = 10. We believe that in a low-dimensional dataset (such as *2D*), or a highly informative dataset (such as *Base case* or any of the higher dimensional scenarios with no irrelevant features where 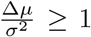) the chains quickly find a “sensible” partition of the data and thus increasing the depth within the chain does not increase the probability that any partition sampled will be closer to the generating partition. For example in figure 11 in the *Small N, large P* case, the distribution of the ARI across the ensembles for which *D* ≥10 and *W* = 1 is nearly identical; this suggests that the chain is sampling a very similar partition again and again for 9,990 iterations (and possibly beyond based upon the PSMs shown in figure 5) and it is through adding more chains rather than using particularly long chains that we improve the ability to uncover the generating structure.

We also notice that even if the behaviour has not stabilised for *D* that the ensemble can uncover meaningful structure. The ARI for the ensembles of short chains can be quite high (as is the case in many of the scenarios). The behaviour of the consensus matrices also shows that low *D* is not a disqualifier from meaningful inference even if longer chains would be ideal, a result that might be useful in real applications with large datasets and complex models. Consider the consensus matrices in figure 7, it can be seen that the behaviour has not stabilised before *D* = 10000 (and possibly there is still some benefit in increasing *D* beyond this value), but the structure being uncovered when there is a sufficient number of chains and *D* is small does correspond to the structure uncovered in the largest and deepest ensemble. We believe that the order in which components merge and items are co-clustered varies depending on initialisation, and thus if the chain is not sufficiently deep that all of the final mergings have occured that a sufficiently large ensemble can still perform meaningful inference of the subpopulation structure despite the poor performance of any individual model. Even though each learner probably has too many clusters for small *D* the consensus among them will have less if the individual learners have low correlation between their partitions (something we might expect if the chains are stopped very early). This is why the entries of the consensus matrix for *D* = 100 and *W* = 100 in figure 7 are more pale than in deeper ensembles; very few items correctly (possibly none) cocluster in every partition, it is only in observing the consensus that the global structure of interest emerges. Thus if there is some limit to the length of chains available for an analysis (e.g. computational or temporal constraints) than the inference obtained from the shorter chains can still be meaningful, with the caveat that the point clustering might have more clusters than the same analysis with longer chains would provide. Additional post-hoc merging of some clusters might be necessary in this case.

In contrast, when the dataset is sparse or contains many irrelevant features, we believe that deeper chains are required to reach this steady-state sampling where no single sample is expected to be better than any other (see the *Irrelevant features (P*_*n*_ = 100) facet of figure 11).

In some scenarios no method is successful in uncovering the generating labels. In the *Large standard deviation (σ*^2^ = 25) and *Small N, large P (*Δ*µ* = 0.2) this is due to the lack of signal - the clusters overlap so significantly that it is not possible for any of these methods to uncover much of the generating structure. In the *No structure* case it is different (although `Mclust` does perform well here). In this case all items are generated from a common distributions. For the Bayesian chains and the ensembles, a clustering of singletons is predicted; each item is allocated a unique label (see figures 8 and 9). While failing to perform well under the ARI, this is a sensible result. Rather than indicating (as we did with the shared label) that no item is particularly distinct from the others and thus all share a common label, this clustering of singletons states that no item is more similar to any other and thus no two items should cluster together. It is an alternative statement of the same result, i.e. that there is no evidence for subpopulation structure. We consider this evidence that an ensemble of Bayesian mixture models is not as susceptible to predicting labels than an ensemble based upon *K*-means clustering as in S, enbabaoğlu et al. (2014a,b).

Increasing *W* is also required when the dimensionality of the dataset is large. In this case it is due to individual chains exploring only a single mode (as can be seen in figure 5 where each chain appears to sample only a single partition). In this example where each sample is a partition that appears to be a mode in the posterior distribution of the allocation vector from very early in the chain (based upon the stable performance for *D* ≥10), increasing *W* allows each chain to “vote” on which mode is the global mode, as we believe that the mode that attracts the most chains is the global mode (although in real datasets the number of chains required might be greater than in our simulations). An example of this behaviour may be seen in figure 10.

In figure 11, limiting behaviour for increases of *W* and *D* can be seen for the ensemble. For most simulations there is no change in performance for greater choices of *W* and *D* after some stabilising values.

**Figure 9:**
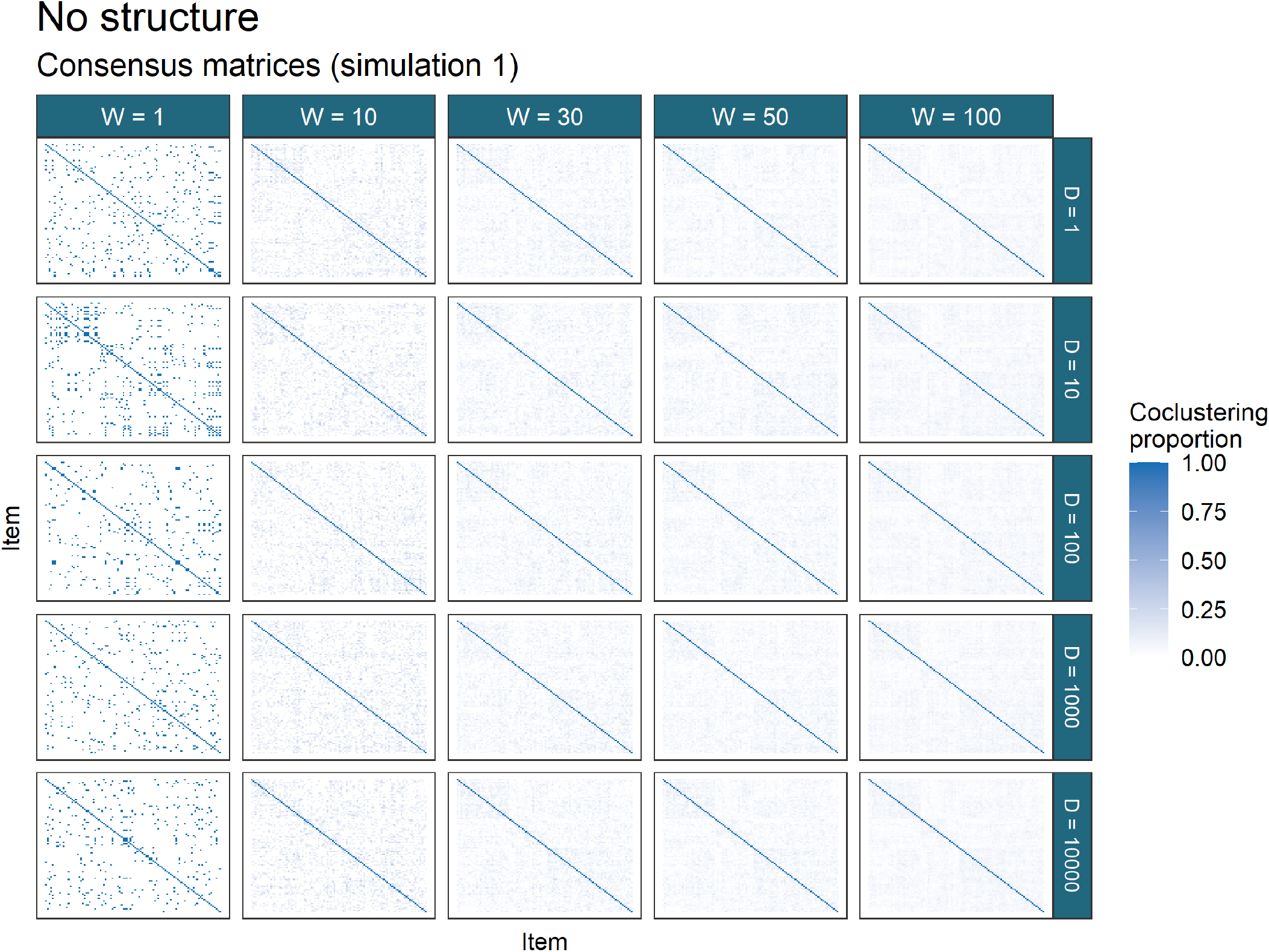
Consensus matrices for simulation 1 of the *No structure* scenario. Each item is allocated to a singleton in many of the Consensus matrices.

**Figure 10:**
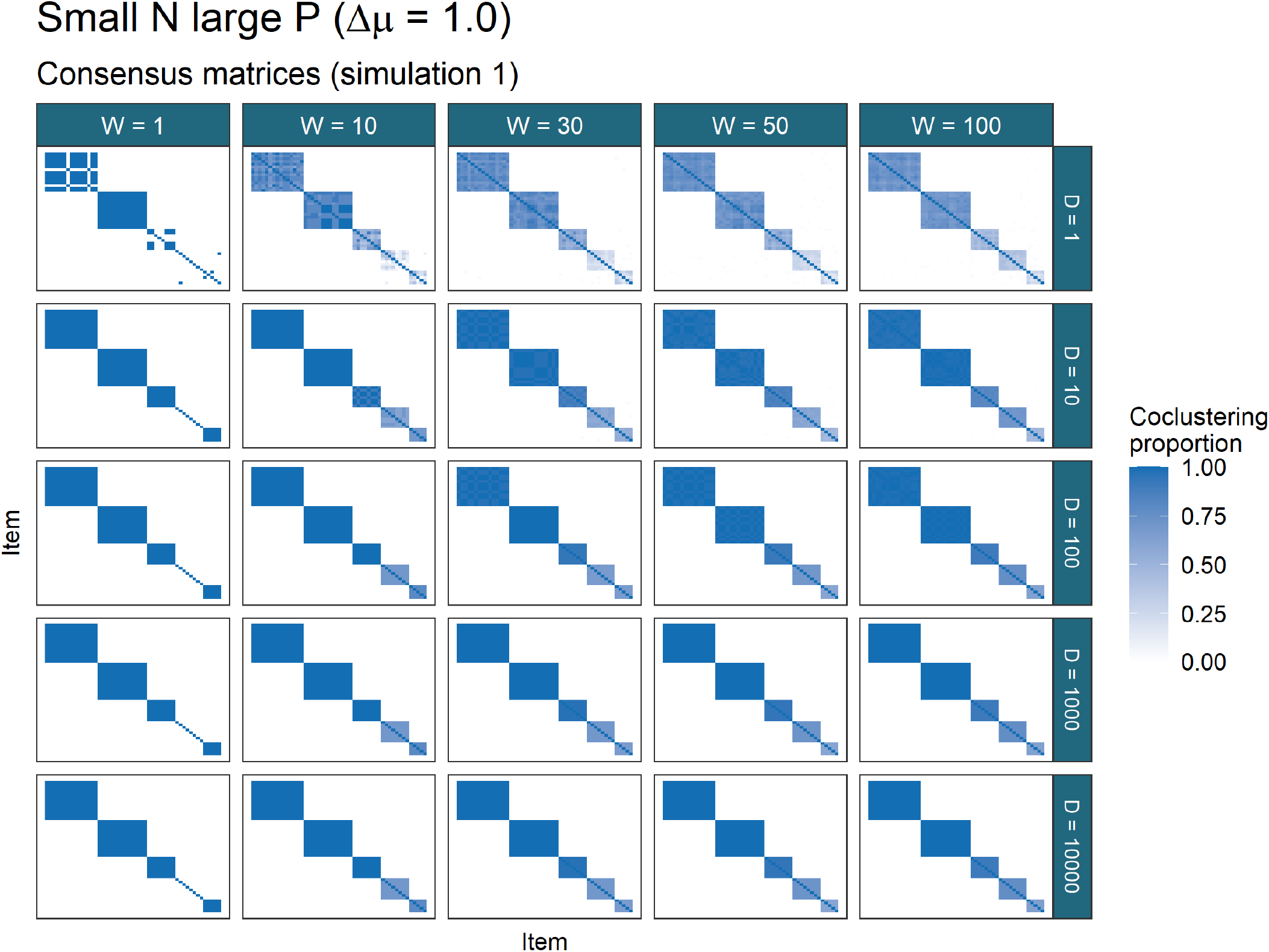
Consensus matrices for simulation 1 of the first *Large N, small P* scenario. One can see that by iteration ten the sample being drawn is from the mode (for *W* = 1, *D* = 10), and that an ensemble of chains does find structure that recalls the generating labels (see figure 11, the ARI for *CC*(10, *s*) is 1.0 for *s >* 1, meaning that the true labels perfectly align with those predicted by the consensus matrix).

**Figure 11:**
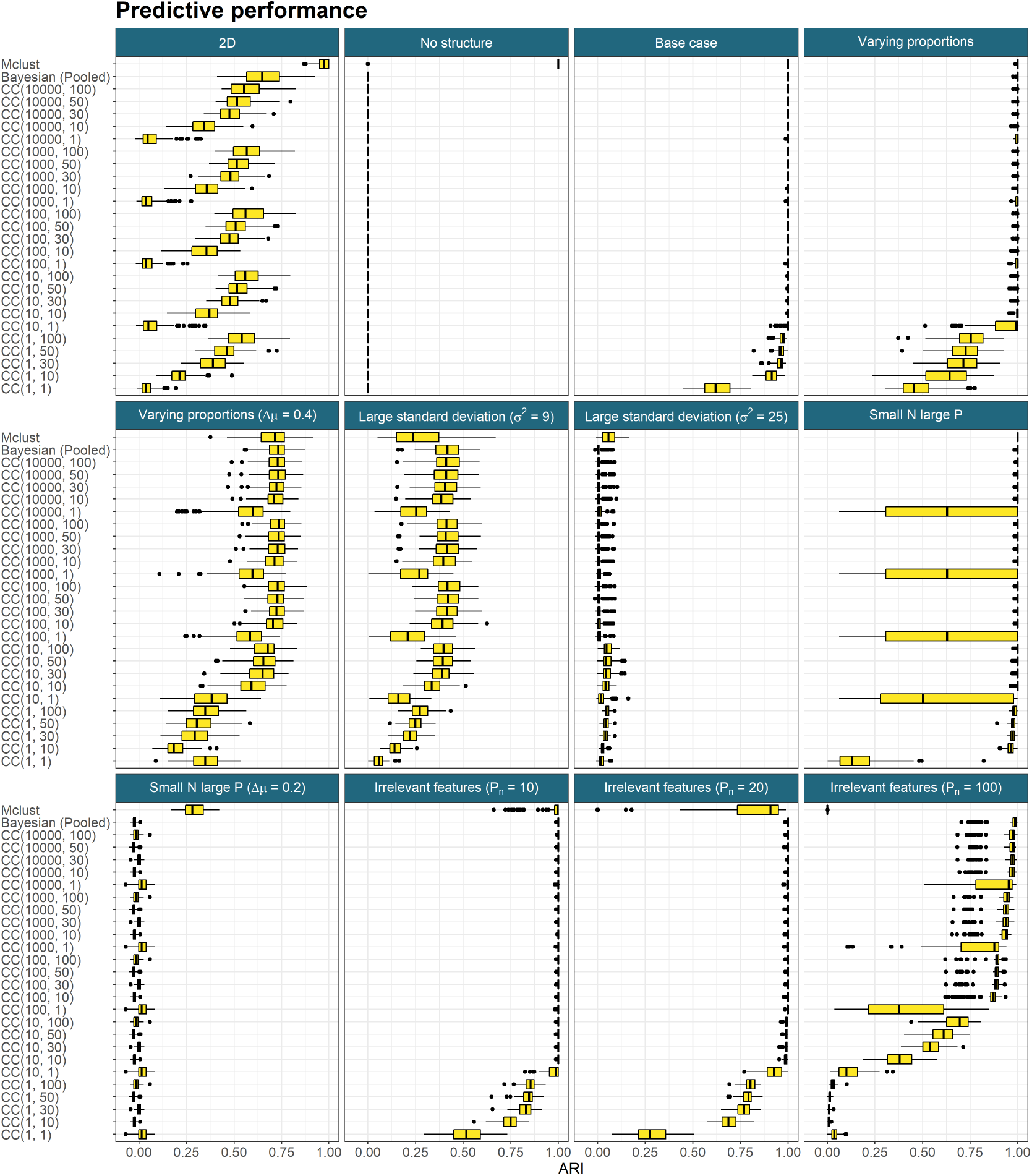
Predictive performance across all simulations. *CC*(*D, W*) denotes consensus clustering using the *D*^*th*^ sample from *W* different chains. In the cases where the generating structure is not exactly found, increasing *D* and *W* sees some improvement in the ARI between the truth and the predicted clusterings before some limiting behaviour emerges and and further increase appears to have no change in the performance.

**Figure 12:**
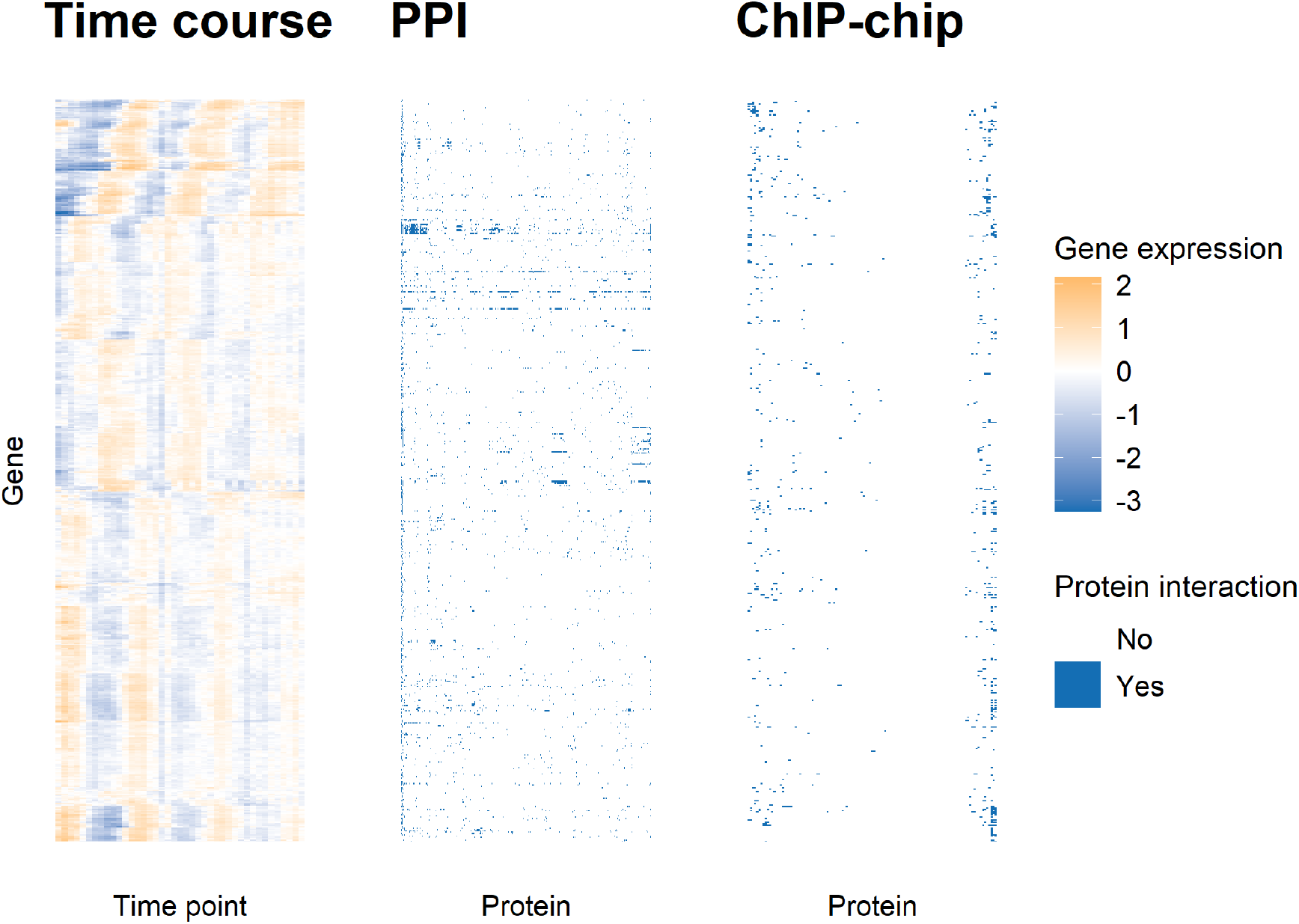
Heatmap of the yeast datasets. Each plot has a common row order corresponding to the gene products being clustered. This order was decided by a hierarchical clustering of the rows of the time course expression matrix. The time course data is associated with the “Gene expression” legend and the ChIP-chip and PPI data with “Protein interaction” legend.

## 5 Multi-omics analysis of the cell cycle in budding yeast

We chose our three datasets (shown in figure 12) to perform an integrative analysis as many of the protein encoding genes in the mitotic cell cycle have well studied genomic binding sites with mapped transcription factors (**TFs**) that control phase-specific expression (Cho et al., 1998; Spellman et al., 1998); thus the inclusion of the ChIP-chip data means that the clusters that align across the datasets should include well studied regulatory proteins and thus be of biological interest. If a cluster of genes are similarly expressed in the time course, share associated regulatory protein in the ChIP-chip and are associated with common protein complexes in the PPI data, than this implies a gene set with strong biological significance.

In contrast, if we cluster the time course dataset alone, any clusters that we find are defined by correlation across time. This might be assumed to be driven by shared regulatory mechanisms, but other sources of structure might be encouraging this, even experimental error. However, if a cluster aligns across both the time course dataset and the ChIP-chip dataset we can be more certain that these genes are part of some regulatory network; if this cluster also emerges in the PPI dataset we might believe that the genes are co-regulated as part of the formation of some protein complex. Furthermore, this integrative aspect means that clusters that might merge in the time course dataset due to similar periodicity in a standalone analysis might remain separate due to different associated transcription factors in the ChIP-chip dataset.

Thus we performed an integrative analysis using MDI to avoid agressive assumptions about either the biology defining any clusters and modelling assumptions about the latent structure.

We expect that the complexity of this data and model means that the time required for convergence of the MCMC algorithm would be very large. We avoid this problem by using consensus clustering of MDI, instead basing our final ensemble choice on the stopping rule described in the main paper.

The datasets were modelled using a mixture of Gaussian processes in the time course dataset and Multinomial distributions in the ChIP-chip and PPI datasets. To ensure that our mixture model is initially overfitted we set 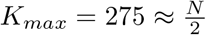, and following from this the point estimate was inferred from the consensus matrix using `maxpear` as in the simulated data except we set `k.max = 275`.

### 5.1 Consensus clustering analysis

We include the consensus matrices for each dataset for a range of ensembles for further evidence that the ensemble was stable for the 10, 000^*th*^ iteration from 1,000 chains in figures 13, 14 and 15. In these figures, there is no strong change between the consensus matrices for *D* = 5001 and *D* = 10001.

We wish to identify groups of genes that tend to be grouped together in multiple datasets. We focus upon the genes that tend to have the same cluster label in multiple datasets, those which have a common label across some set of datasets in more than half of the observed clusterings, or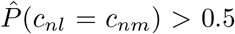, where *c*_*nl*_ denotes the cluster label of gene *n* in dataset *l*. This based upon the the concept of *fused genes* proposed by Savage et al. (2010) and used by Kirk et al. (2012), but to avoid confusion due to other possible ideas of fused genes (e.g. those that contribute to a common protein complex, the behaviour of TFs upon a gene) we avoid this term. These genes with common clustering across datasets are those most affected by the integrative aspect of the analysis and therefore we focus upon these in the our cluster analysis. In our case we have the possible sets of:

- {Time course}, {ChIP-chip}, {PPI},
- {Time course, ChIP-chip}, {Time course, PPI}, {ChIP-chip, PPI}, and
- {Time course, ChIP-chip, PPI}.

The number of genes meeting this criteria between any two datasets is indicative of how strongly they influence each other and is expected to align with the *ϕ*_*lm*_ parameters from the MDI model. We find the following number of unique genes integrated between each combination of datasets:

**Figure 13:**
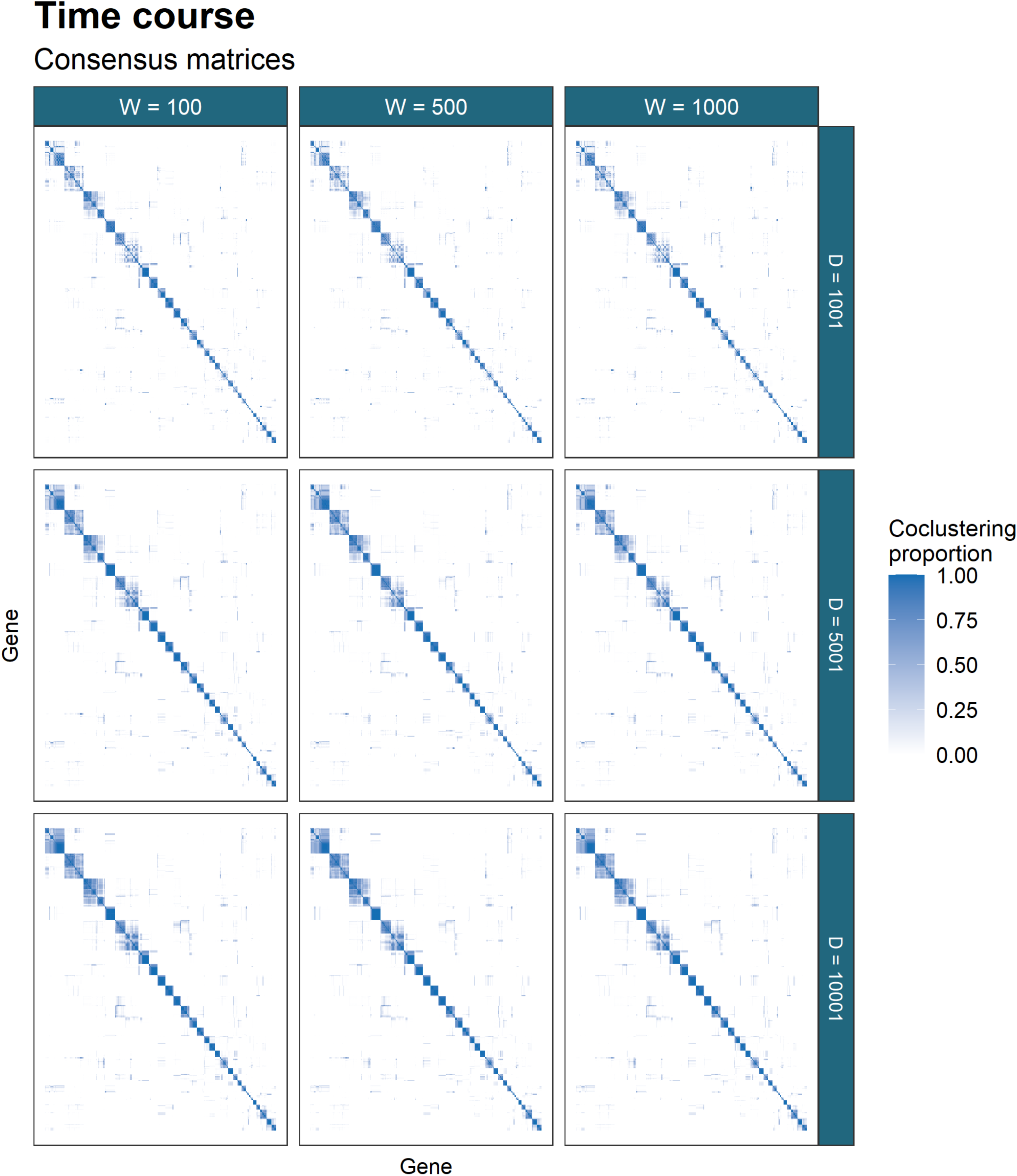
Consensus matrices for different ensembles of MDI for the time course data. This dataset has stable clustering across the different choices of number of chains, *W*, and chain depth, *D*, with some components merging as the chain depth increases.

**Figure 14:**
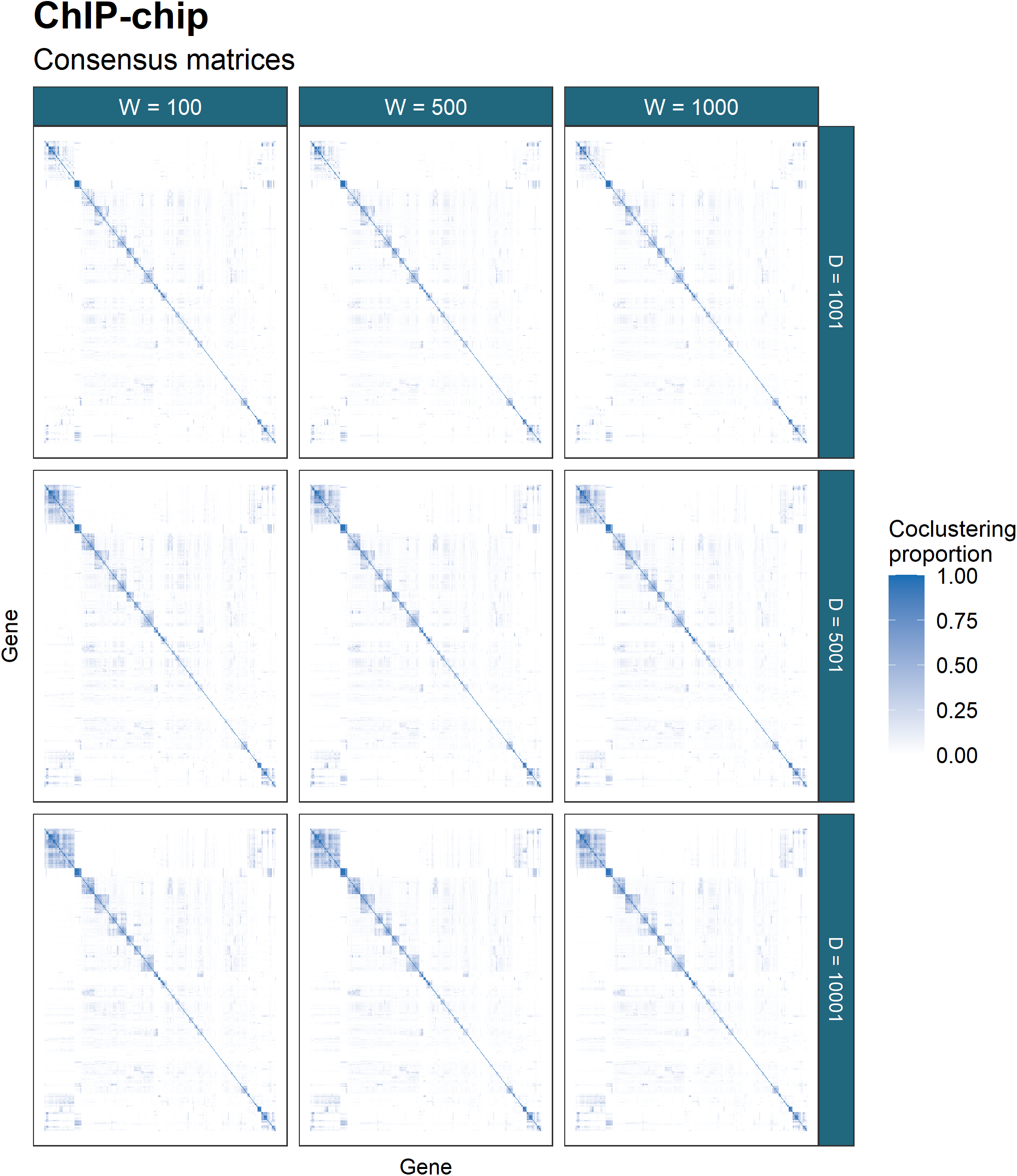
The ChIP-chip dataset is more sparse than the time course data. In keeping with the results from the simulations for mixture models, deeper chains are required for better performance. It is only between *D* = 5, 001 and *D* = 10, 001 that no change in the clustering can be observed and the result is believed to be stable. In this dataset the number of chains used, *W*, appears relatively unimportant, with similar results for *W* = 100, 500, 1000.

**Figure 15:**
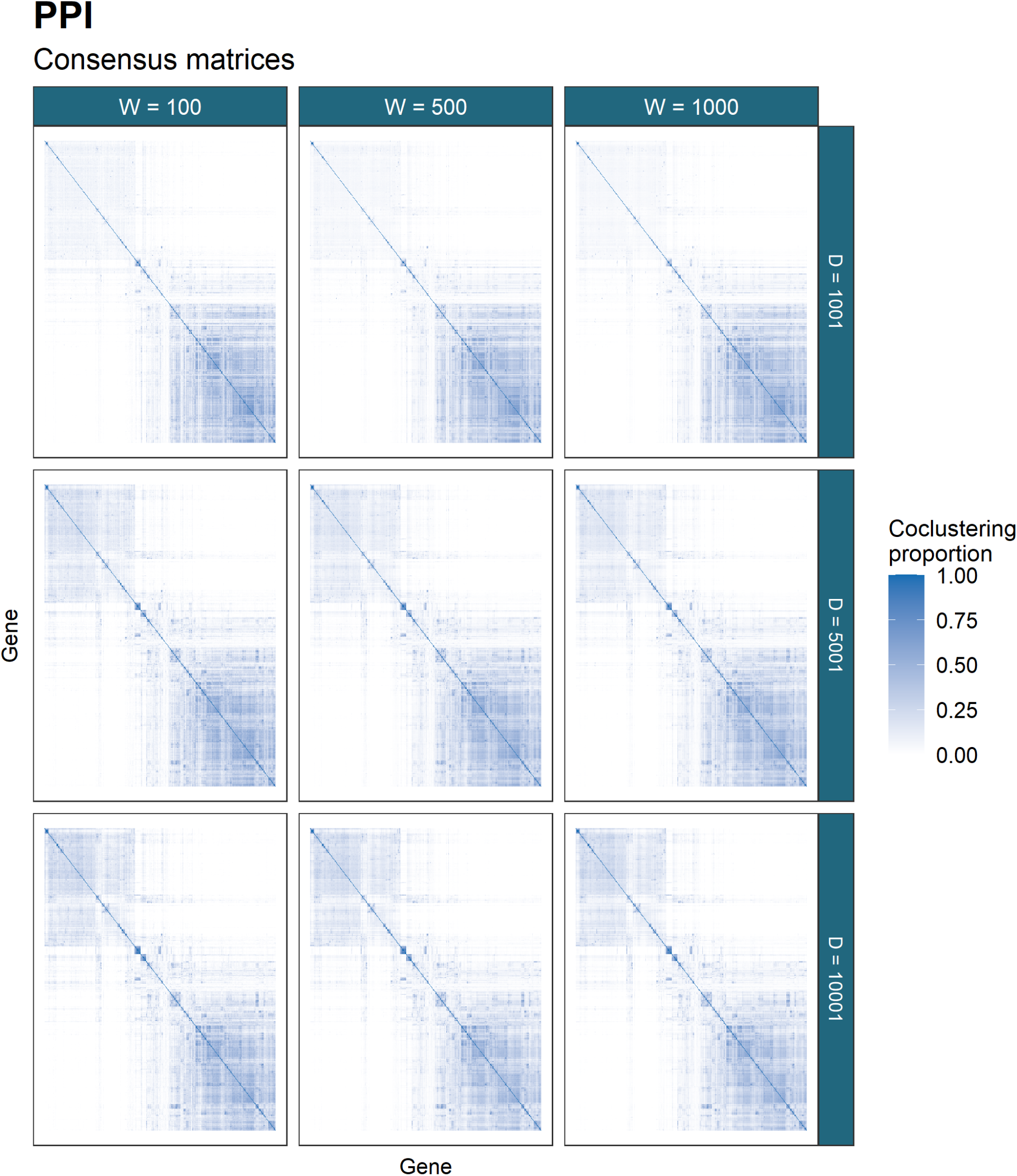
The PPI dataset has awkward characteristics for modelling. A wide, sparse dataset it is chain depth that we found to be the most important parameter for the ensemble. Similar to the results in figure 14, the matrices only stabilise from *D* = 5001 to *D* = 10001.

- Time course + ChIP-chip + PPI: 56,
- Time course + ChIP-chip: 205 (261 including the 56 integrated across all datasets),
- Time course + PPI: 12 (68),
- ChIP-chip + PPI: 43 (99)..

This shows that the time course and ChIP-chip datasets contain very similar structure, the ChIP-chip and PPI datasets have some similarity but significantly less and the time course and PPI datasets have less shared signal again.

Compare this to the original analysis of this data in Kirk et al. (2012), where the number of such genes in each combination is:

- Time course + ChIP-chip + PPI: 16,
- Time course + ChIP-chip: 32 (48),
- Time course + PPI: 16 (32),
- ChIP-chip + PPI: 15 (31).

Our analysis has found significantly more shared structure.

#### 5.1.1 Time course ChIP-chip analysis

We focus upon the dataset pairing of time course + ChIP-chip within the integrative analysis as the combination with the greatest number of genes with shared clustering. We show these genes grouped by their inferred cluster in figure 16. In this plot we exclude the 15 clusters where more than half of the member genes have no interactions in the ChIP-chip data and any clusters of one. We find that a small number of transcription factors dominate, with different combinations emerging across the 10 clusters shown here in table 2. Many of these 10 correspond to transcription factors that are well known to regulate cell cycle expression, namely MBP1, SWI4, SWI6, MCM1, FKH1, FKH2, NDD1, SWI5, and ACE2 (Simon et al., 2001).

**Table S2:**
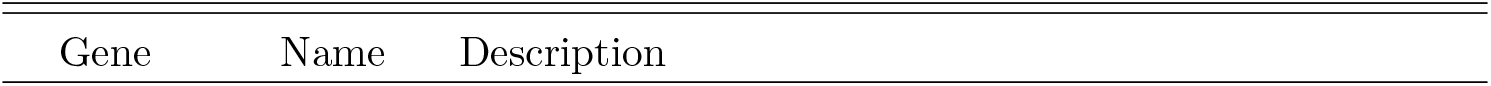
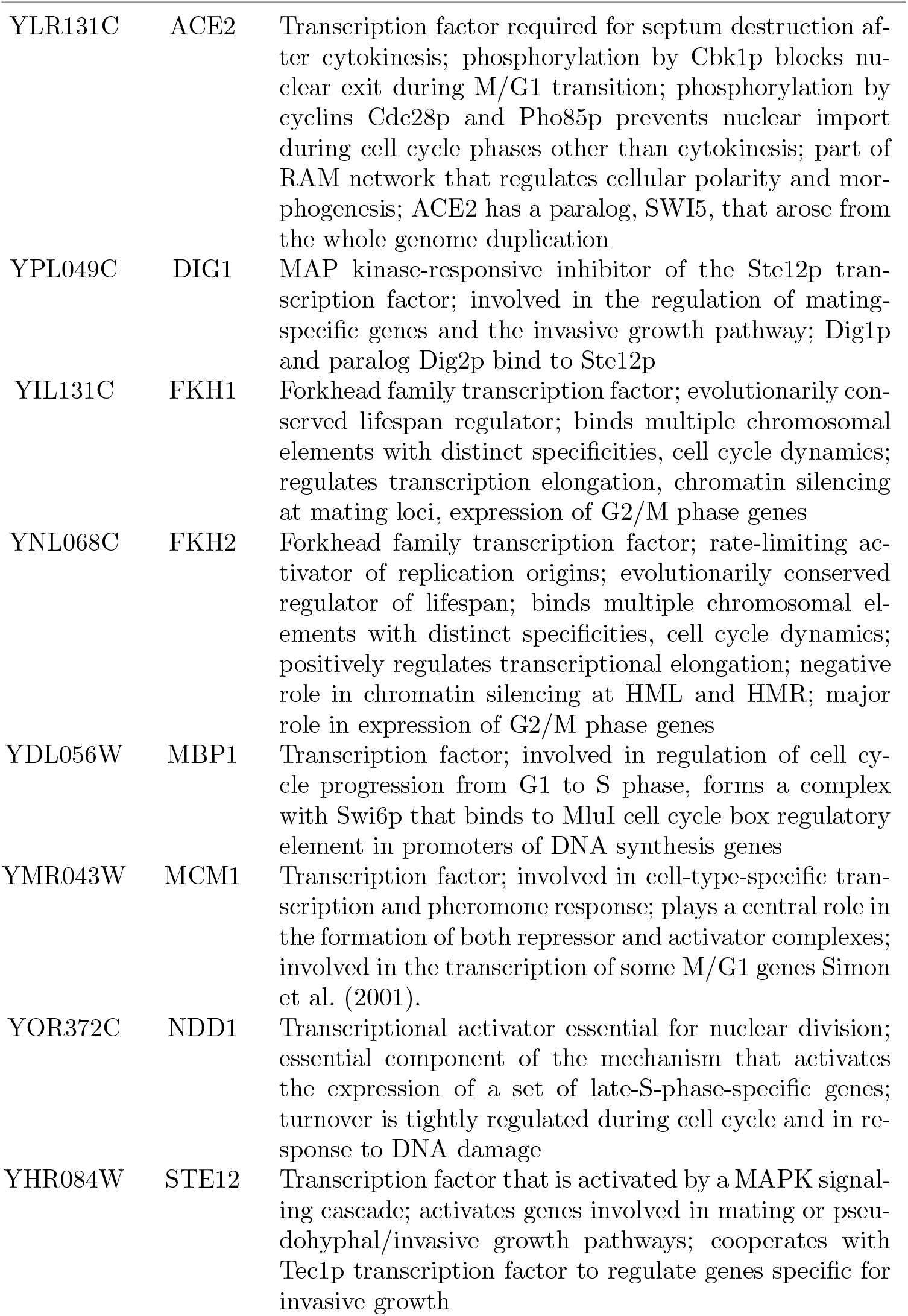
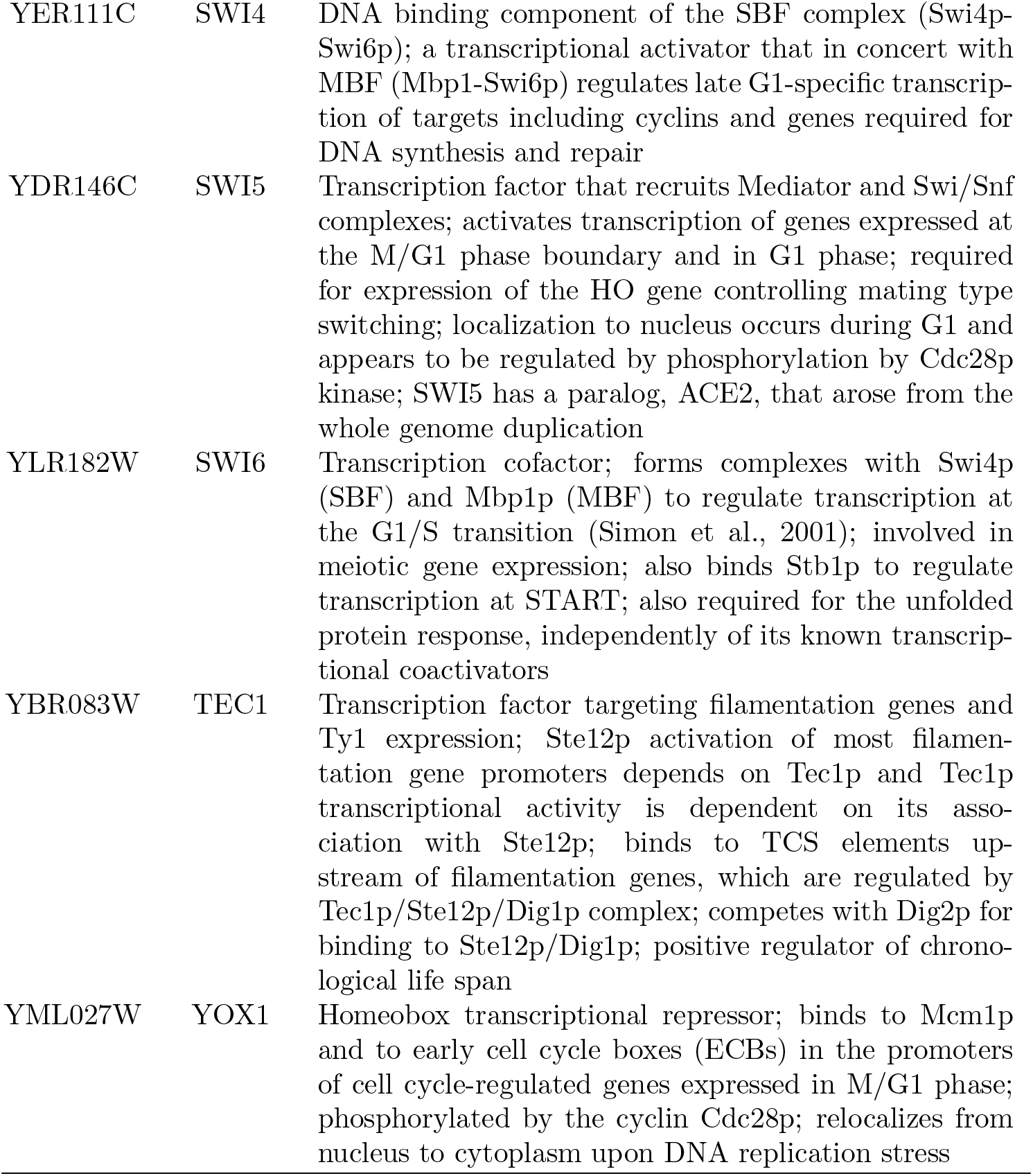
Table of transcription factors prominent in clusters of genes with shared labels for a majority of samples for the time course and ChIP-chip datasets.

|  |  |  |
| --- | --- | --- |
| YLR131C | ACE2 | Transcription factor required for septum destruction after cytokinesis; phosphorylation by Cbk1p blocks nuclear exit during M/G1 transition; phosphorylation by cyclins Cdc28p and Pho85p prevents nuclear import during cell cycle phases other than cytokinesis; part of RAM network that regulates cellular polarity and morphogenesis; ACE2 has a paralog, SWI5, that arose from the whole genome duplication |
| YPL049C | DIG1 | MAP kinase-responsive inhibitor of the Ste12p transcription factor; involved in the regulation of mating-specific genes and the invasive growth pathway; Dig1p and paralog Dig2p bind to Ste12p |
| YIL131C | FKH1 | Forkhead family transcription factor; evolutionarily conserved lifespan regulator; binds multiple chromosomal elements with distinct specificities, cell cycle dynamics; regulates transcription elongation, chromatin silencing at mating loci, expression of G2/M phase genes |
| YNL068C | FKH2 | Forkhead family transcription factor; rate-limiting activator of replication origins; evolutionarily conserved regulator of lifespan; binds multiple chromosomal elements with distinct specificities, cell cycle dynamics; positively regulates transcriptional elongation; negative role in chromatin silencing at HML and HMR; major role in expression of G2/M phase genes |
| YDL056W | MBP1 | Transcription factor; involved in regulation of cell cycle progression from G1 to S phase, forms a complex with Swi6p that binds to MluI cell cycle box regulatory element in promoters of DNA synthesis genes |
| YMR043W | MCM1 | Transcription factor; involved in cell-type-specific transcription and pheromone response; plays a central role in the formation of both repressor and activator complexes; involved in the transcription of some M/G1 genes Simon et al. (2001). |
| YOR372C | NDD1 | Transcriptional activator essential for nuclear division; essential component of the mechanism that activates the expression of a set of late-S-phase-specific genes; turnover is tightly regulated during cell cycle and in response to DNA damage |
| YHR084W | STE12 | Transcription factor that is activated by a MAPK signaling cascade; activates genes involved in mating or pseudohyphal/invasive growth pathways; cooperates with Tec1p transcription factor to regulate genes specific for invasive growth |
| YER111C | SWI4 | DNA binding component of the SBF complex (Swi4p-Swi6p); a transcriptional activator that in concert with MBF (Mbp1-Swi6p) regulates late G1-specific transcription of targets including cyclins and genes required for DNA synthesis and repair |
| YDR146C | SWI5 | Transcription factor that recruits Mediator and Swi/Snf complexes; activates transcription of genes expressed at the M/G1 phase boundary and in G1 phase; required for expression of the HO gene controlling mating type switching; localization to nucleus occurs during G1 and appears to be regulated by phosphorylation by Cdc28p kinase; SWI5 has a paralog, ACE2, that arose from the whole genome duplication |
| YLR182W | SWI6 | Transcription cofactor; forms complexes with Swi4p (SBF) and Mbp1p (MBF) to regulate transcription at the G1/S transition (Simon et al., 2001); involved in meiotic gene expression; also binds Stb1p to regulate transcription at START; also required for the unfolded protein response, independently of its known transcriptional coactivators |
| YBR083W | TEC1 | Transcription factor targeting filamentation genes and Ty1 expression; Ste12p activation of most filamentation gene promoters depends on Tec1p and Tec1p transcriptional activity is dependent on its association with Ste12p; binds to TCS elements upstream of filamentation genes, which are regulated by Tec1p/Ste12p/Dig1p complex; competes with Dig2p for binding to Ste12p/Dig1p; positive regulator of chronological life span |
| YML027W | YOX1 | Homeobox transcriptional repressor; binds to Mcm1p and to early cell cycle boxes (ECBs) in the promoters of cell cycle-regulated genes expressed in M/G1 phase; phosphorylated by the cyclin Cdc28p; relocates from nucleus to cytoplasm upon DNA replication stress |

These regulatory proteins are found in different combinations across the clusters. Based upon these combinations we associate each cluster with phases of the cell cycle and or some specific processes.

- Cluster 1: both ACE2 and SWI5 emerge. These regulate specific genes at the end of M and early G1 (McBride et al., 1999; Simon et al., 2001).
- Cluster 2: SWI5. This is similar to cluster 1, as ACE2 is a paralog of SWI5; therefore associated with M/G1. Furthermore, inspection of the expression in the timecourse data shows that the members of cluster 2 largely differentiate from those of cluster 1 based upon amplitude, not periodicity, suggesting that these clusters could be merged.

- Cluster 5: MBP1, SWI4 and SWI6. The SBF complex (Swi4p-Swi6p) is a transcriptional activator that in concert with MBF (Mbp1-Swi6p) regulates late G1-specific transcription of targets including cyclins and genes required for DNA synthesis and repair, controlling the transition to S phase (Simon et al., 2001; Iyer et al., 2001; Aligianni et al., 2009).
- Cluster 9: MBP1 and SWI6. These combine to form MBF, which regulates DNA replication and repair (Iyer et al., 2001).
- Cluster 11: DIG1, SWI4, SWI6, and STE12 emerge in all members with some having associations with TEC1. TEC1 and STE12, controls development, including cell adhesion and filament formation and is negatively regulated by DIG1 and DIG2 (van der Felden et al., 2014).
- Cluster 12: MBP1, SWI4 and SWI6. Similar to cluster 5 in both the time course and ChIP-chip datasets and thus G1/S phase.
- Cluster 16: some MBP1, SWI4 and SWI6. The constituents of this cluster are largely associated with proteins contributing to histones H1, H2A, H2B, H3 and H4, suggesting an S-phase cluster (Ewen, 2000).
- Cluster 17: FKH1 and FKH2. Fkh1p and Fkh2p are required for cell-cycle regulation of transcription during G2/M (Kumar et al., 2000).
- Cluster 20: NDD1 and MCM1 with some FKH2. Mcm1, together with Fkh1 or Fkh2, recruits the Ndd1 protein in late G2, and thus controls the transcription of G2/M genes (Simon et al., 2001; Koranda et al., 2000).
- Cluster 26: YOX1 and MCM1. YOX1 binds to Mcm1p and to early cell cycle boxes (ECBs) in the promoters of cell cycle-regulated genes expressed in M/G1 phase (Pramila et al., 2002).

**Table S3:** Description of the genes with common labelling across the time course and ChIP-chip datasets from clusters 9 and 16.

| Gene | Name | Cluster | Description |
| --- | --- | --- | --- |
| YJL115W | ASF1 | 9 | Nucleosome assembly factor; involved in chromatin assembly, disassembly; required for buffering mRNA synthesis rate against gene dosage changes in S phase |
| YLR103C | CDC45 | 9 | DNA replication initiation factor; recruited to MCM pre-RC complexes at replication origins; recruits elongation machinery; binds tightly to ssDNA, which disrupts interaction with the MCM helicase and stalls it during replication stress; mutants in human homolog may cause velocardiofacial and DiGeorge syndromes |
| YPL241C | CIN2 | 9 | GTPase-activating protein (GAP) for Cin4p; tubulin folding factor C involved in beta-tubulin (Tub2p) folding; mutants display increased chromosome loss and benomyl sensitivity; human homolog RP2 complements yeast null mutant |
| YPR175W | DPB2 | 9 | Second largest subunit of DNA polymerase II (DNA polymerase epsilon); required for maintenance of fidelity of chromosomal replication; essential motif in C-terminus is required for formation of the four-subunit Pol epsilon; expression peaks at the G1/S phase boundary; Cdc28p substrate |
| YIL026C | IRR1 | 9 | Subunit of the cohesin complex; which is required for sister chromatid cohesion during mitosis and meiosis and interacts with centromeres and chromosome arms |
| YCL061C | MRC1 | 9 | S-phase checkpoint protein required for DNA replication; couples DNA helicase and polymerase; defines a novel S-phase checkpoint with Hog1p that coordinates DNA replication and transcription upon osmostress; protects uncapped telomeres; Dia2p-dependent degradation mediates checkpoint recovery; mammalian claspin homolog; subunit of a replication-pausing checkpoint complex, Tof1p-Mrc1p-Csm3p; checkpoint-mediator protein that functions during DNA replication and activates the effector kinase Rad53 (Bando et al., 2009); human ATR homolog (Lao et al., 2018) |
| YDR097C | MSH6 | 9 | Protein required for mismatch repair in mitosis and meiosis; forms a complex with Msh2p to repair both single-base and insertion-deletion mispairs; also involved in interstrand cross-link repair; potentially phosphorylated by Cdc28p |
| YNL102W | POL1 | 9 | Catalytic subunit of the DNA polymerase I alpha-primase complex; required for the initiation of DNA replication during mitotic DNA synthesis and premeiotic DNA synthesis |
| YBL035C | POL12 | 9 | B subunit of DNA polymerase alpha-primase complex; required for initiation of DNA replication during mitotic and premeiotic DNA synthesis; also functions in telomere capping and length regulation |
| YKL113C | RAD27 | 9 | 5' to 3' exonuclease, 5' flap endonuclease; required for Okazaki fragment processing and maturation, for long-patch base-excision repair and large loop repair (LLR), ribonucleotide excision repair |
| YPL153C | RAD53 | 9 | DNA damage response protein kinase; required for cell-cycle arrest, regulation of copper genes in response to DNA damage; human homolog CHEK2 implicated in breast cancer can complement yeast null mutant |
| YAR007C | RFA1 | 9 | Subunit of heterotrimeric Replication Protein A (RPA); RPA is a highly conserved single-stranded DNA binding protein involved in DNA replication, repair, and recombination; RPA protects against inappropriate telomere recombination, and upon telomere uncapping, prevents cell proliferation by a checkpoint-independent pathway; role in DNA catenation/decatenation pathway of chromosome disentangling; relocates to the cytosol in response to hypoxia |
| YNL312W | RFA2 | 9 | Subunit of heterotrimeric Replication Protein A (RPA); RPA is a highly conserved single-stranded DNA binding protein involved in DNA replication, repair, and recombination; RPA protects against inappropriate telomere recombination, and upon telomere uncapping, prevents cell proliferation by a checkpoint-independent pathway; in concert with Sgs1p-Top2p-Rmi1p, stimulates DNA catenation/decatenation activity of Top3p; protein abundance increases in response to DNA replication |
| YAR008W | SEN34 | 9 | Subunit of the tRNA splicing endonuclease; tRNA splicing endonuclease (Sen complex) is composed of Sen2p, Sen15p, Sen34p, and Sen54p; Sen complex also cleaves the CBP1 mRNA at the mitochondrial surface; Sen34p contains the active site for tRNA 3' splice site cleavage and has similarity to Sen2p and to Archaeal tRNA splicing endonuclease |
| YJL074C | SMC3 | 9 | Subunit of the multiprotein cohesin complex; required for sister chromatid cohesion in mitotic cells; also required, with Rec8p, for cohesion and recombination during meiosis; phylogenetically conserved SMC chromosomal ATPase family member |
| YNL273W | TOF1 | 9 | Subunit of a replication-pausing checkpoint complex; Tof1p-Mrc1p-Csm3p acts at the stalled replication fork to promote sister chromatid cohesion after DNA damage, facilitating gap repair of damaged DNA; interacts with the MCM helicase; checkpoint-mediator protein that functions during DNA replication and activates the effector kinase RAD53 (Bando et al., 2009); human ATM homolog (Lao et al., 2018) |
| YMR215W | GAS3 | 16 | Putative 1,3-beta-glucanosyltransferase; has similarity to other GAS family members; low abundance, possibly inactive member of the GAS family of GPI-containing proteins; localizes to the cell wall; mRNA induced during sporulation |
| YBR009C | HHF1 | 16 | Histone H4; core histone protein required for chromatin assembly and chromosome function; one of two identical histone proteins (see also HHF2); contributes to telomeric silencing; N-terminal domain involved in maintaining genomic integrity |
| YNL030W | HHF2 | 16 | Histone H4; core histone protein required for chromatin assembly and chromosome function; one of two identical histone proteins (see also HHF1); contributes to telomeric silencing; N-terminal domain involved in maintaining genomic integrity |
| YPL127C | HHO1 | 16 | Histone H1, linker histone with roles in meiosis and sporulation; decreasing levels early in sporulation may promote meiosis, and increasing levels during sporulation facilitate compaction of spore chromatin; binds to promoters and within genes in mature spores; may be recruited by Ume6p to promoter regions, contributing to transcriptional repression outside of meiosis; suppresses DNA repair involving homologous recombination |
| YBR010W | HHT1 | 16 | Histone H3; core histone protein required for chromatin assembly, part of heterochromatin-mediated telomeric and HM silencing; one of two identical histone H3 proteins (see HHT2); regulated by acetylation, methylation, and phosphorylation; H3K14 acetylation plays an important role in the unfolding of strongly positioned nucleosomes during repair of UV damage |
| YNL031C | HHT2 | 16 | Histone H3; core histone protein required for chromatin assembly, part of heterochromatin-mediated telomeric and HM silencing; one of two identical histone H3 proteins (see HHT1); regulated by acetylation, methylation, and phosphorylation; H3K14 acetylation plays an important role in the unfolding of strongly positioned nucleosomes during repair of UV damage |
| YDR225W | HTA1 | 16 | Histone H2A; core histone protein required for chromatin assembly and chromosome function; one of two nearly identical subtypes (see also HTA2); DNA damage-dependent phosphorylation by Mec1p facilitates DNA repair; acetylated by Nat4p; N-terminally propionylated in vivo |
| YBL003C | HTA2 | 16 | Histone H2A; core histone protein required for chromatin assembly and chromosome function; one of two nearly identical (see also HTA1) subtypes; DNA damage-dependent phosphorylation by Mec1p facilitates DNA repair; acetylated by Nat4p |
| YDR224C | HTB1 | 16 | Histone H2B; core histone protein required for chromatin assembly and chromosome function; nearly identical to HTB2; Rad6p-Bre1p-Lge1p mediated ubiquitination regulates reassembly after DNA replication, transcriptional activation, meiotic DSB formation and H3 methylation |
| YBL002W | HTB2 | 16 | Histone H2B; core histone protein required for chromatin assembly and chromosome function; nearly identical to HTB1; Rad6p-Bre1p-Lge1p mediated ubiquitination regulates reassembly after DNA replication, transcriptional activation, meiotic DSB formation and H3 methylation |
| YNR009W | NRM1 | 16 | Transcriptional co-repressor of MBF-regulated gene expression; Nrm1p associates stably with promoters via MCB binding factor (MBF) to repress transcription upon exit from G1 phase |
| YDR113C | PDS1 | 16 | Securin; inhibits anaphase by binding separin Esp1p; blocks cyclin destruction and mitotic exit, essential for meiotic progression and mitotic cell cycle arrest; localization is cell-cycle dependent and regulated by Cdc28p phosphorylation |

### 5.2 Bayesian analysis

We wished to compare our results from consensus clustering to a conventional Bayesian approach. We ran 10 chains of MDI for 36 hours saving every thousandth sample. This resulted in chains of varying length. We reduced the chains to 666 samples as this was the number of samples achieved by the shortest chain. Similar to section 4.3 these chains were then investigated for

- within-chain stationarity using the Geweke convergence diagnostic (Geweke et al., 1991), and
- across-chain convergence using 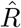 (Gelman et al., 1992) and the Vats-Knudson extension (*stable* 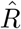, Vats and Knudson, 2018).

Again we focus upon stationarity of the continuous variables. In the implementation of MDI we used, the recorded continuous variables are the concentration parameters of the Dirichlet distribution for the dataset-specific component weights and the *ϕ*_*ij*_ parameter associated with the correlation between the *i*^*th*^ and *j*^*th*^ datasets.

We plot the Geweke-statistic for each chain in figure 17. No chain is perfectly behaved; as we cannot reduce to the set of stationary chains we thus exclude the most poorly behaved chains. Our lack of belief in the convergence of these chains is fortified by the behaviour of 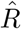 (which can be seen in figure 18) and the different distributions sampled for the *ϕ*_*lm*_ parameters shown in figure 19.

**Figure 16:**
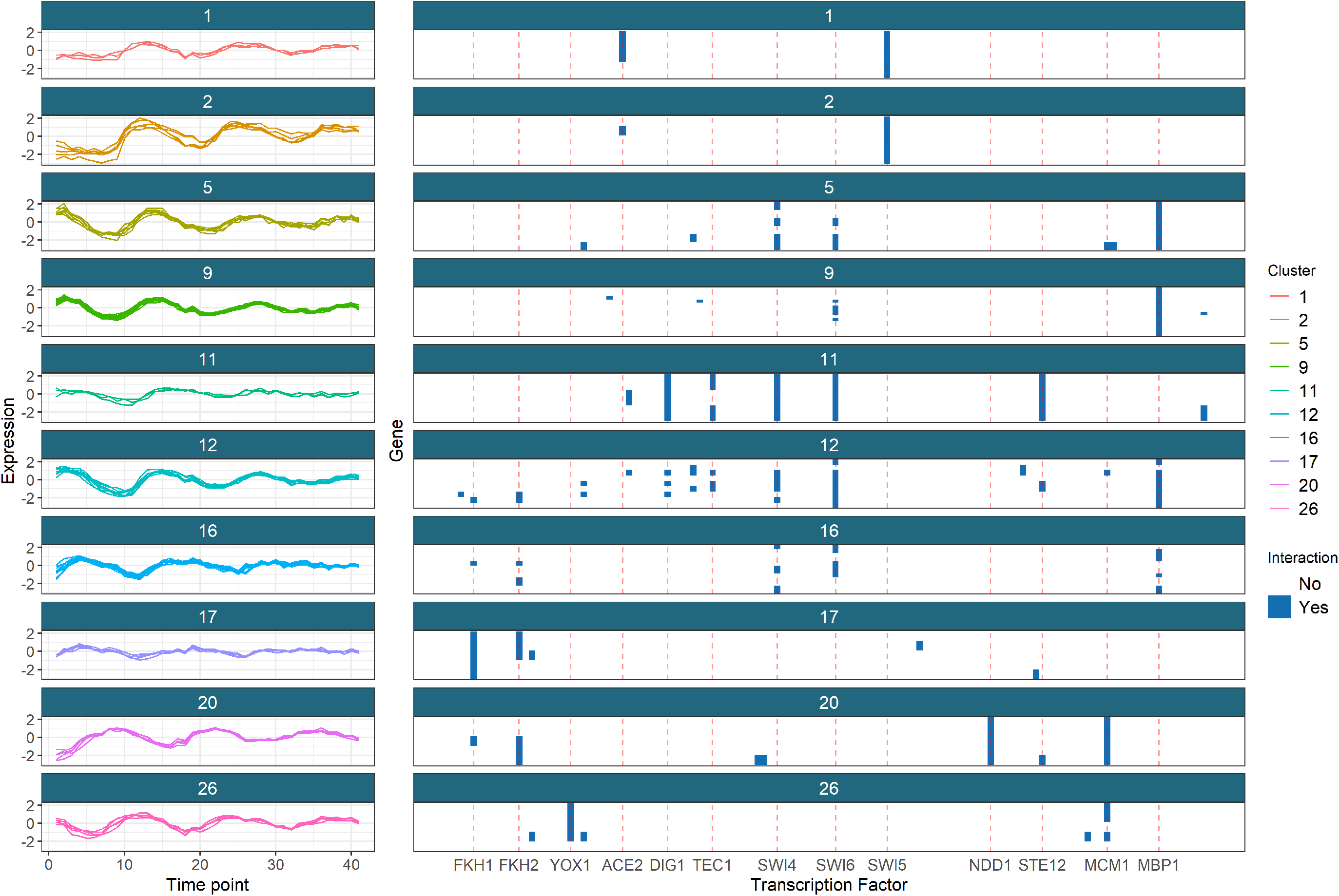
The clusters of genes with common labels across the time course and ChIP-chip datasets (as described in tbale **??**). We exclude the clusters with no interactions in the ChIP-chip dataset and include a red line for the Transcription factors that dominate the clustering structure in the ChIP-chip dataset.

**Figure 17:**
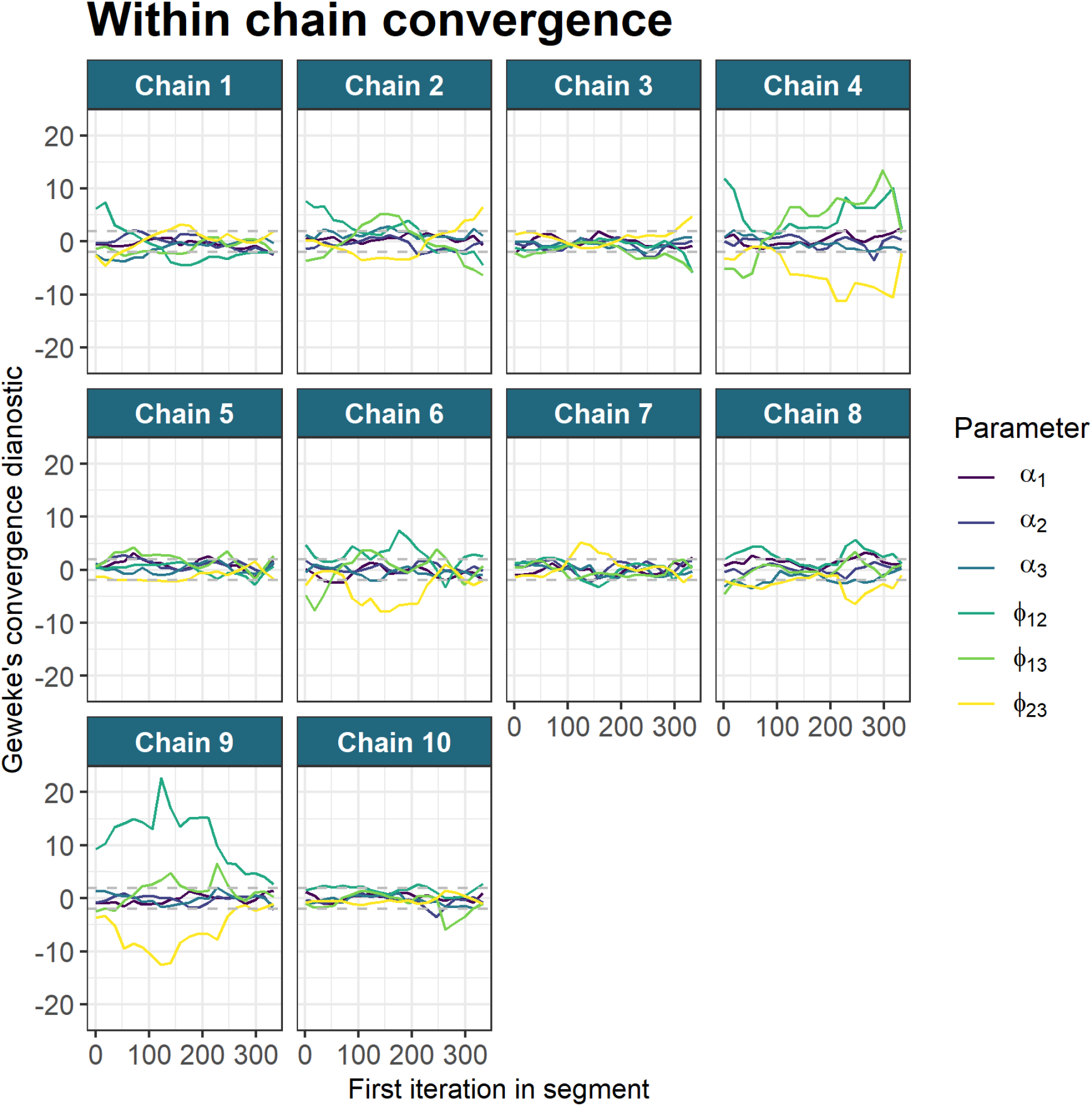
Chain 9 can be seen to have the most extreme behaviour in the distribution of the Geweke diagnostic for the parameters. We remove this chain from the analysis. Of the remaining chains we believe that 1, 2, 4 and 6 express the distributions furthest removed from the desired behaviour and are dropped from the analysis.

**Figure 18:**
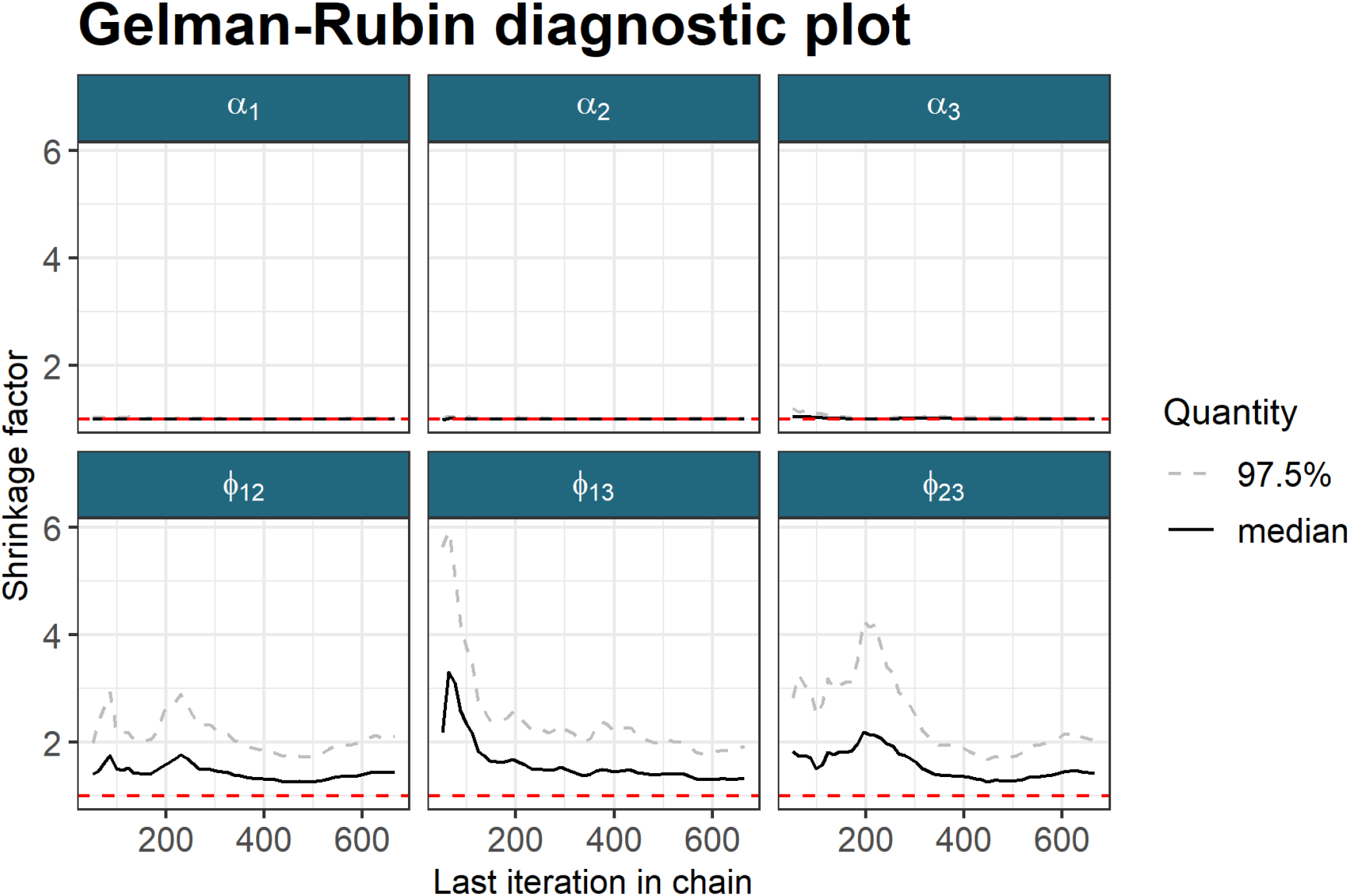
The chains still appear to be unconverged with 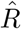 remaining above 1.25 for the *ϕ*_12_, *ϕ*_13_ and *ϕ*_23_ parameters. Stable 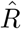 is also too high with values of 1.049, 1.052 and 1.057 for *ϕ*_12_, *ϕ*_13_ and *ϕ*_23_ respectively. The values of *α*_*l*_ cannot be seen due to the scaling of the *y*-axis.

We visualise the the PSMs for each dataset in figure 20.

If we compare the distribution of sampled values for the *ϕ* parameters for the Bayesian chains that we keep based upon their convergence diagnostics, the final ensemble used (*D* = 10001, *W* = 1000) and the pooled samples from the 5 long chains, then we see that the ensemble consisting of the long chains (which might be believed to sampling different parts of the posterior distribution) is closer in its appearance to the distributions sampled by the consensus clustering than to any single chain.

**Figure 19:**
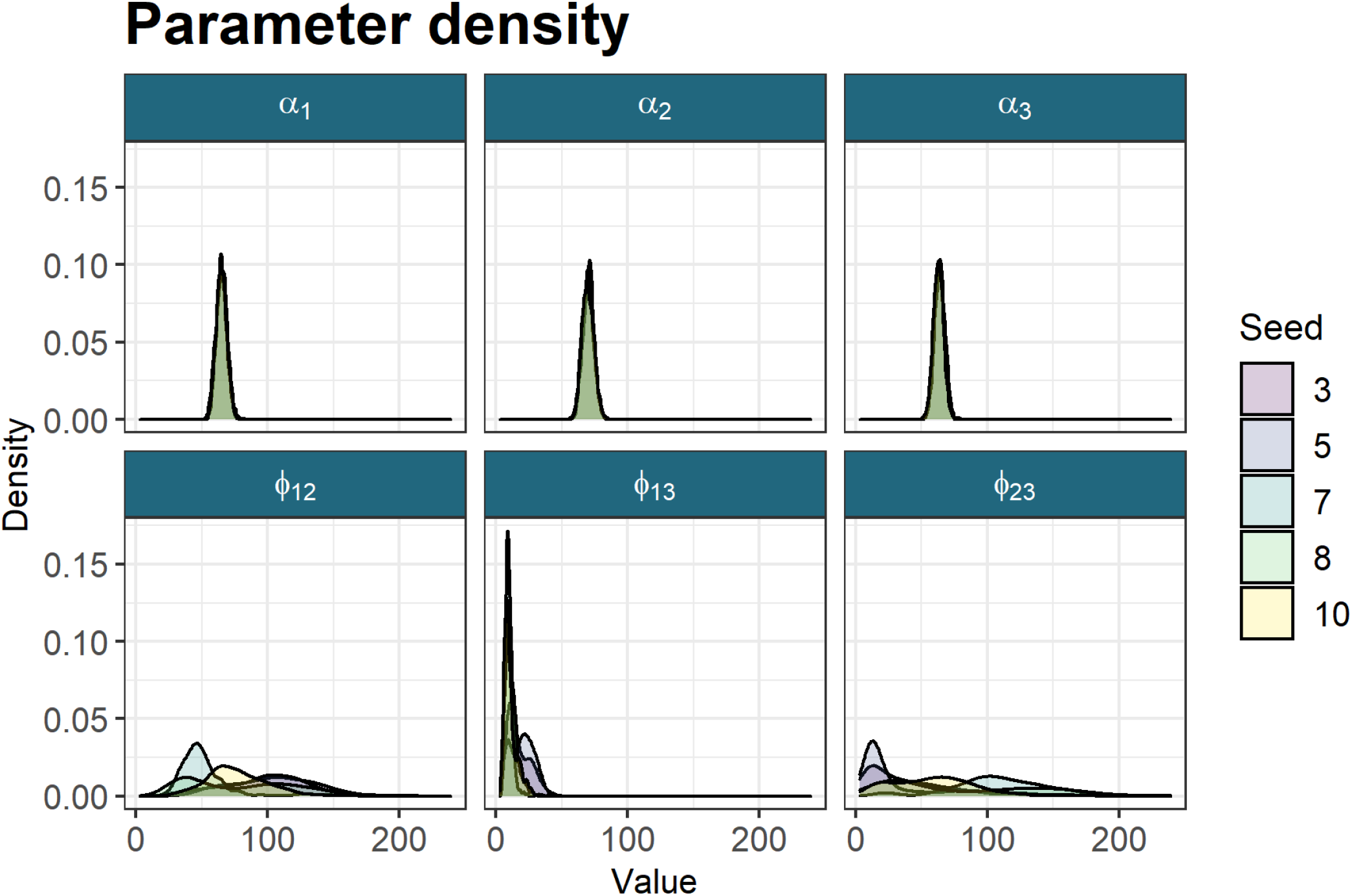
The densities of the continuous variables across the 5 chains kept for analysis. The mean sampled values are *α*_1_ = 64.84, *α*_2_ = 69.85, *α*_2_ = 63.22, *ϕ*_12_ = 81.76, *ϕ*_13_ = 13.87, and *ϕ*_23_ = 65.03. It can be seen that different modes are being sampled for the *ϕ* parameters in each chain.

**Figure 20:**
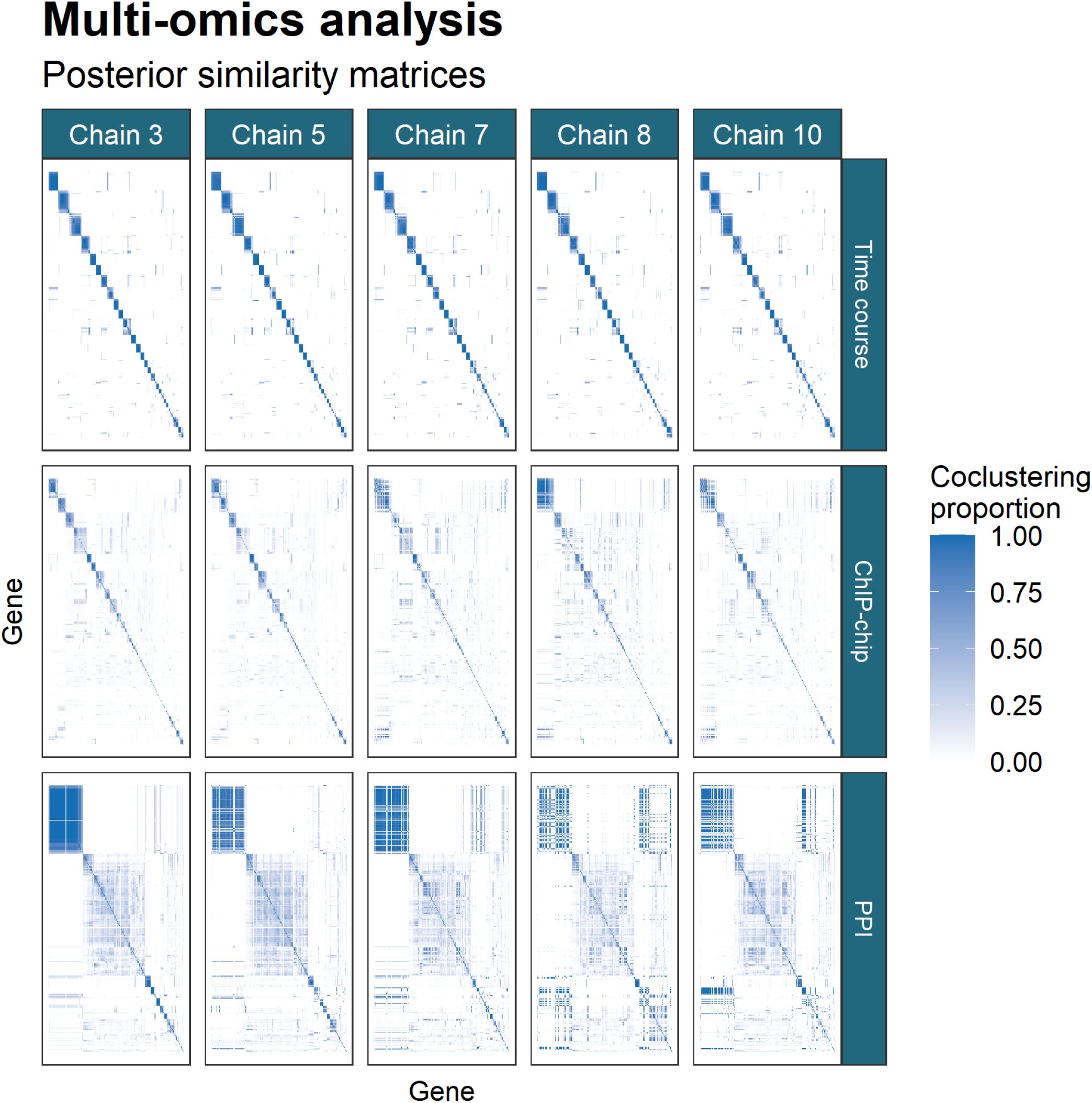
PSMs for each chain within each dataset. The PSMs are ordered by hierarchical clustering of the rows of the PSM for chain 3 in each dataset. There is no marked difference between the matrices for the time course data with disagreement becoming more prominent in the ChIP-chip data and more so again in the PPI dataset.

**Figure 21:**
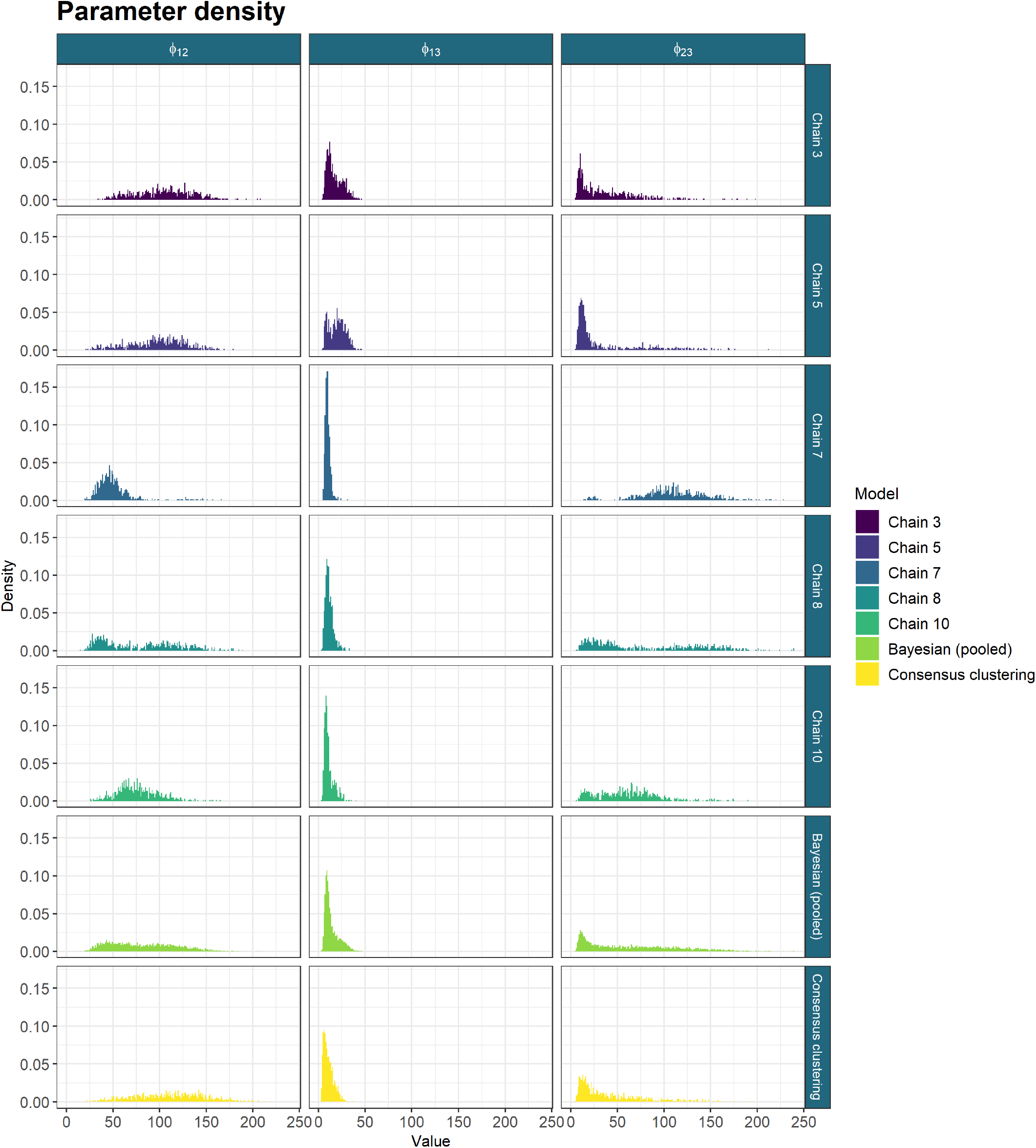
The sampled values for the *ϕ* parameters from the long chains, their pooled samples and the consensus using 1000 chains of depth 10,001. The long chains display a variety of behaviours. Across chains there is no clear consensus on the nature of the posterior distribution. The samples from any single chain are not particularly close to the behaviour of the pooled samples across all three parameters. It is the consensus clustering that most approaches this pooled behaviour.

### 5.3 GO term over-representation

We further show the lack of disagreement between the long chains from section 5.2 in a Gene Ontology (GO) term over-representation analysis. We estimated clusterings from the PSMs of the chains kept from section 5.2 visualised in figure 20 and the consensus matrix of the largest ensemble run (i.e. *CC*(10001, 1000)) using the `maxpear` function from the R package `mcclust` Fritsch (2012) using default settings except for `k.max` which was set to 275 ≈*N/*2. To perform the GO term over-representation analysis we used the `Bioconductor` packages `clusterProfiler` (Yu et al., 2012), `biomaRt` (Durinck et al., 2009) and the annotation package `org.Sc.sgd.db` (Carlson et al., 2014).

We conditioned the test on the background set of the 551 yeast genes in the data. The gene labelled YIL167W was not found in the annotation database and was dropped from the analysis leaving a background universe of 550 genes. A hypergeometric test was used to check if the number of genes associated with specific GO terms within a cluster was greater than expected by random chance. We corrected the *p*-values using the Benjamini-Hochberg correction (Benjamini and Hochberg, 1995) and defined significance by a threshold of 0.01. We plotted the over-represented GO terms for the different clusterings within each dataset using the three different ontologies of “Molecular function” (**MF**), “Biological process” (**BP**) and “Cellular component” (**CC**) (figures 22, 23 and 24 respectively).

As we expect based upon the disagreement shown in figure 21, we find that the Bayesian chains have very significant disagreements between each other; there is no consensus on the results with many terms enriched in one or two chains. However, the consensus clustering finds many of the terms common to all of the long chains. This is what we would expect based upon the similarity of the *ϕ*_*lm*_ distribution in the ensemble and the pooled long chains. Consensus clustering also finds some terms with low *p*-values common to a majority of chains (such as DNA helicase activity in the MF ontology for the time course dataset) and a small number of GO terms unique to itself. These terms that no long chain find are normally related to other terms already over-represented within either the consensus clustering or a number of the long chains. For example, the transmembrane transporter activity and transporter activity terms uncovered by the ensemble in the time course dataset are related to terms found across 3 of the chains and by consensus clustering (specifically transferase activity and phosphotransferase).

**Figure 22:**
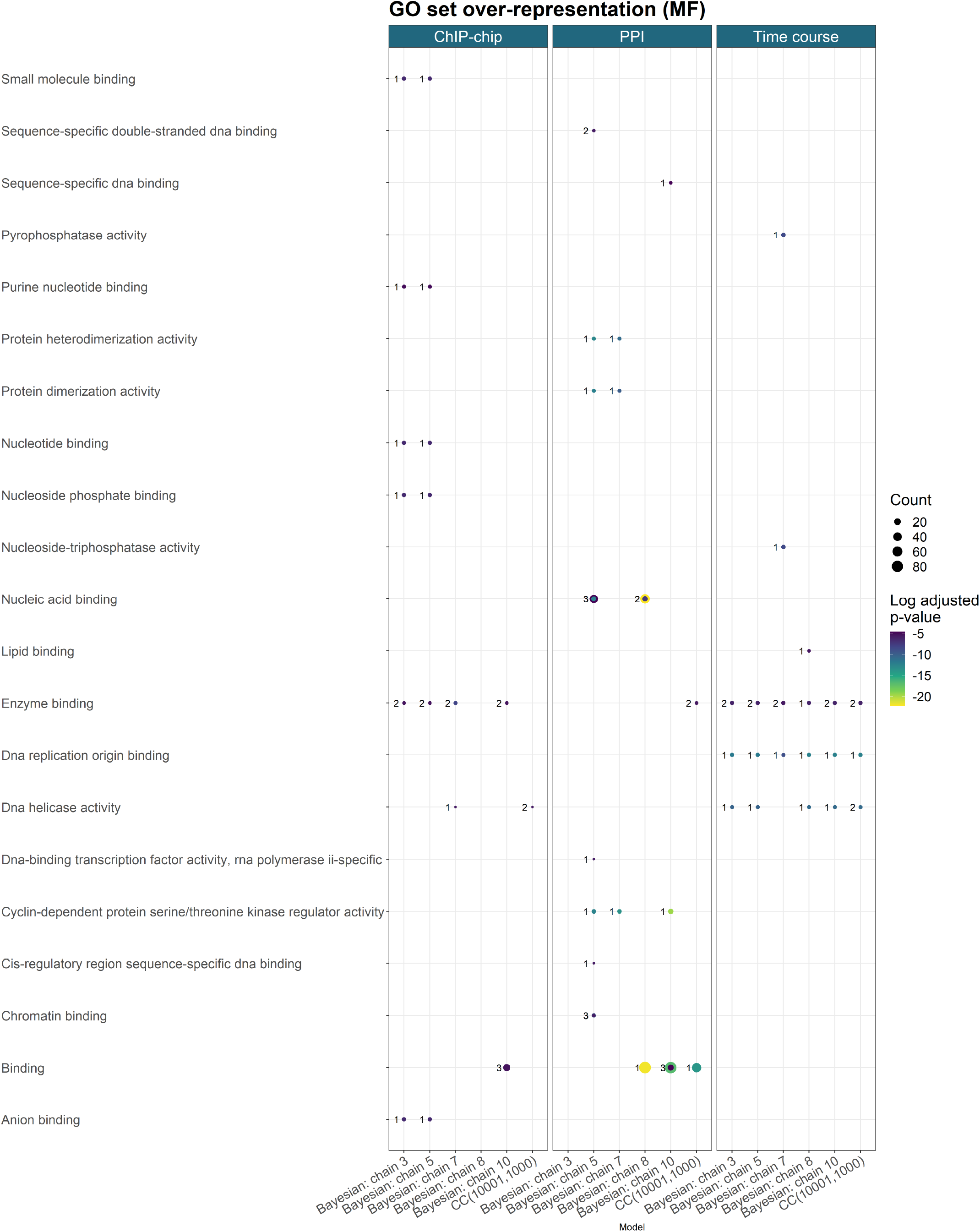
GO term over-representation for the Molecular function ontology for each dataset from the final clustering of each method.

**Figure 23:**
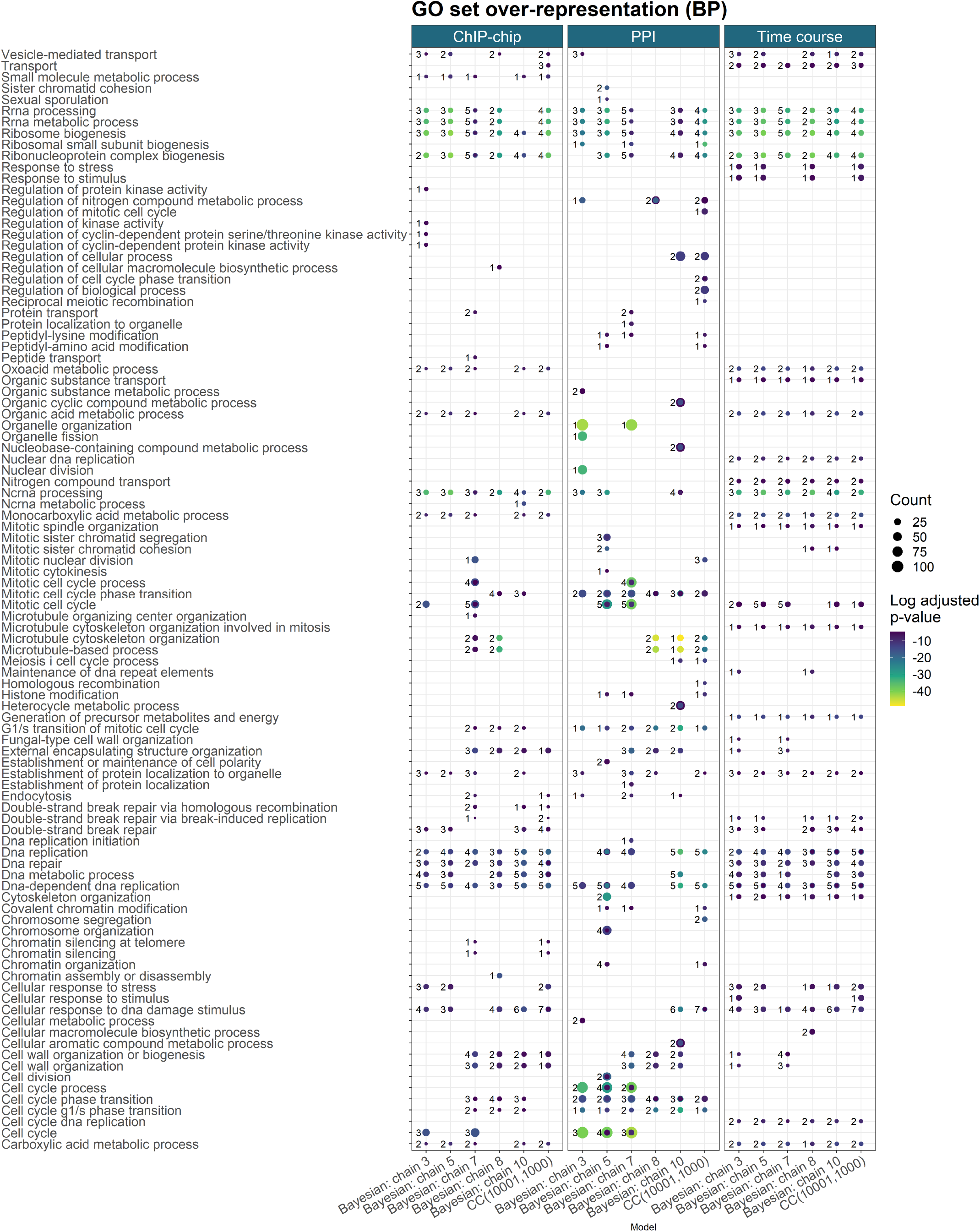
GO term over-representation for the Biological process ontology for each dataset from the final clustering of each method.

**Figure 24:**
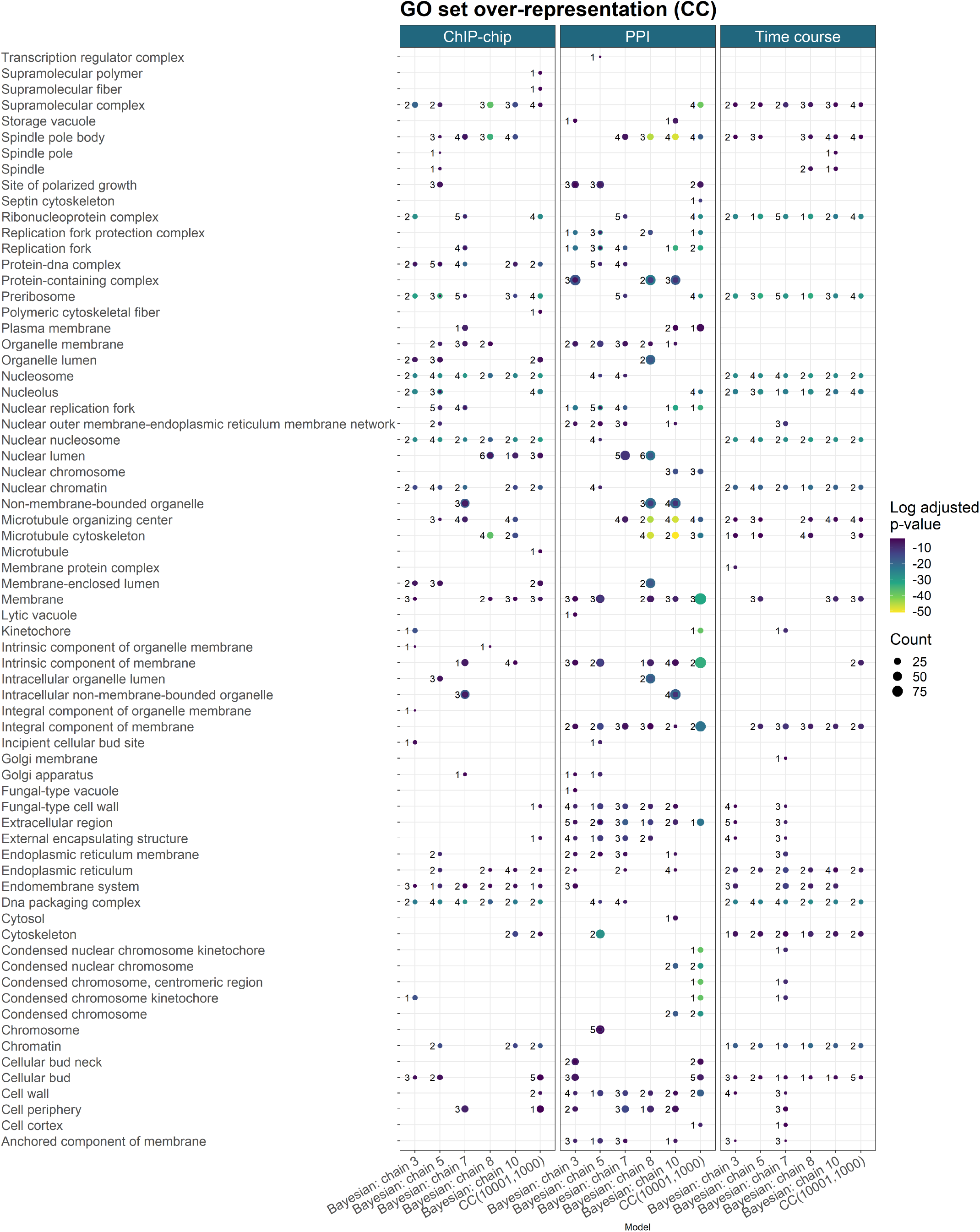
GO term over-representation for the Cellular component ontology for each dataset from the final clustering of each method.

## Notes

### Competing Interest Statement

The authors have declared no competing interest.

### Summary of Updates

Novelty and motivation highlighted more clearly. Method differentiated from similarly named method.

